# Principal component analysis of RNA-seq data unveils a novel prostate cancer-associated gene expression signature

**DOI:** 10.1101/2020.10.26.355750

**Authors:** Yasser Perera, Augusto Gonzalez, Rolando Perez

**Author notes:** Correspondence: Y.P.

## Abstract

Prostate cancer (Pca) is a highly heterogeneous disease and the second more common tumor in males. Molecular and genetic profiles have been used to identify subtypes and guide therapeutic intervention. However, roughly 26% of primary Pca are driven by unknown molecular lesions. We use Principal Component Analysis (PCA) and custom RNAseq-data normalization to identify a gene expression signature which segregates primary PRAD from normal tissues. This Core-Expression Signature (PRAD-CES) includes 33 genes and accounts for 39% of data complexity along the PC1-cancer axis. The PRAD-CES is populated by protein-coding (AMACR, TP63, HPN) and RNA-genes (PCA3, ARLN1), validated/predicted biomarkers (HOXC6, TDRD1, DLX1), and/or cancer drivers (PCA3, ARLN1, PCAT-14). Of note, the PRAD-CES also comprises six over-expressed LncRNAs without previous Pca association, four of them potentially modulating driver’s genes TMPRSS2, PRUNE2 and AMACR. Overall, our PCA capture 57% of data complexity within PC1-3. GO enrichment and correlation analysis comprising major clinical features (i.e., Gleason Score, AR Score, TMPRSS2-ERG fusion and Tumor Cellularity) suggest that PC2 and PC3 gene signatures may describe more aggressive and inflammation-prone transitional forms of PRAD. Of note, surfaced genes may entail novel prognostic biomarkers and molecular alterations to intervene. Particularly, our work uncovered RNA genes with appealing implications on Pca biology and progression.

## 1. Introduction

Prostate cancer (Pca) is the second most common cancer in men [1]. Multiple genetic and demographic factors contribute to the incidence of Pca [2]. Prostate-specific antigen (PSA) screening allows detection of nearly 90% of prostate cancers at initial stages when their surgical removal is the preferred medical intervention [3]. Of note, during their they life-time, most of these patients would never experience Pca, therefore the disease is considered over-diagnosed and over-treated [4].

The clinical outcome of Pca is highly variable, and precise prediction of disease’s course is not possible [5]. Major risk stratification systems are based on clinical and pathological parameters such as Gleason score, PSA levels, TNM system and surgical margins [6]. However, the above risk stratification systems fail to adequately predict outcome in many cases [7,8]; thus, novel serum-, urinary-, and tissue-based biomarkers are constantly tested and implemented [9]. Of note, for those tumors spreading beyond the prostatic gland (i.e., local and/or distant metastasis) the prognosis is more dismal, and effective therapies are needed [10,11]. Renewed expectations are still rooted into emerging and hopefully more tractable Pca molecular alterations [12,13].

Comprehensible genome-wide analysis of primary Prostate Adenocarcinoma (PRAD) revealed already known and novel molecular lesions for 74% of all tumors [14]. The most common alterations were fusions of androgen-regulated promoters with ERG and other members of the E26 transformation-specific (ETS) family of transcription factors. Particularly, the TMPRSS2-ERG fusion is the most representative molecular lesion, accounting for 46% of study cases. Pca also show varying degrees of DNA copy-number alteration, whereas somatic point mutations are relatively less common [15,16]. Despite this detailed molecular taxonomy of PRAD, roughly 26% of primary Pca of both, good and poor prognosis, are driven by unknown molecular lesions [14].

Principal Component Analysis (PCA) is an unsupervised analysis method providing information about major directions of data variability and structure, thus reducing the overall dimensionality of complex datasets to a few dominant components [17]. Based on global gene expression data, PCA usually reveals underlying population heterogeneity, including cell differentiation stages, malignant phenotypes and treatment-induced changes, which can be linked to phenotypes and further characterized [18]. Biological meanings are usually capture by the first 3-4 PCs, although further improvements on PCA revealed that higher dimensions may also entail biology information [19].

Recently, we used Principal Component Analysis (PCA) analysis of RNA-seq expression data to show that a relatively small number of “core genes” can segregate normal from neoplastic tissues in different tumor localizations [20]. Here, by using such PCA we analyze primary PRAD RNAseq data to uncover and characterize a novel PRAD-Core Expression Signature (PRAD-CES) which may may “describe” at expression level Pca [21,22]. The PRAD-CES segregates tumor from normal samples along what we call the cancer axis (i.e., PC1), whereas top genes populating PC2 and PC3 might reflects a more aggressive and inflammation-prone transitional forms of PRAD. Overall, the list of surfaced genes may entail novel prognostic biomarkers and/or molecular alterations to intervene. Particularly appealing, was the identification of several RNA genes with potential implications on Pca biology and progression.

## 4. Materials and Methods

### RNA-seq data

For PCA we take RNA-seq tissue expression data from the TCGA Prostate Adenocarcinoma project (TCGA-PRAD, https://portal.gdc.cancer.gov/repository, Accessed in March 2019). The data is in the number of fragments per kilo base of gene length per mega-base of reads format (FPKM). The studied cases include 499 tumor samples and 52 normal samples. At Cbioportal (https://www.cbioportal.org/) such data belong to Prostate Adenocarcinoma (TCGA, Firehose Legacy) cohort. Two other data cohorts were used in particular analysis: Prostate Adenocarcinoma (TCGA, Cell 2015) and Prostate Adenocarcinoma (MSKCC, Cancer Cell 2010).

### PCA analysis

Fig. 1a shows in a typical PRAD sample that the expression of more than 35000 genes is below 0.1. We shift the ex-pression by 0.1 in such a way that, when computed the differential expressions, genes with not statistically significant expressions are ruled out of the analysis. Then, we take the mean geometric average over normal samples in order to define the reference expression for each gene, and normalize accordingly to obtain the differential expressions, ē = e/eref. Finally, we take the base 2 logarithm, ê = Log2 (ē), to define the fold variation. Besides reducing the variance, the logarithm allows treating over- and sub-expression in a symmetrical way. The co-variance matrix is defined in terms of ê. We forced the reference for the PC analysis to be at the center of the cloud of normal samples, ê = 0. This is what actually happens in a population, where most individuals are healthy and cancer situations are rare.

**Figure 1.**
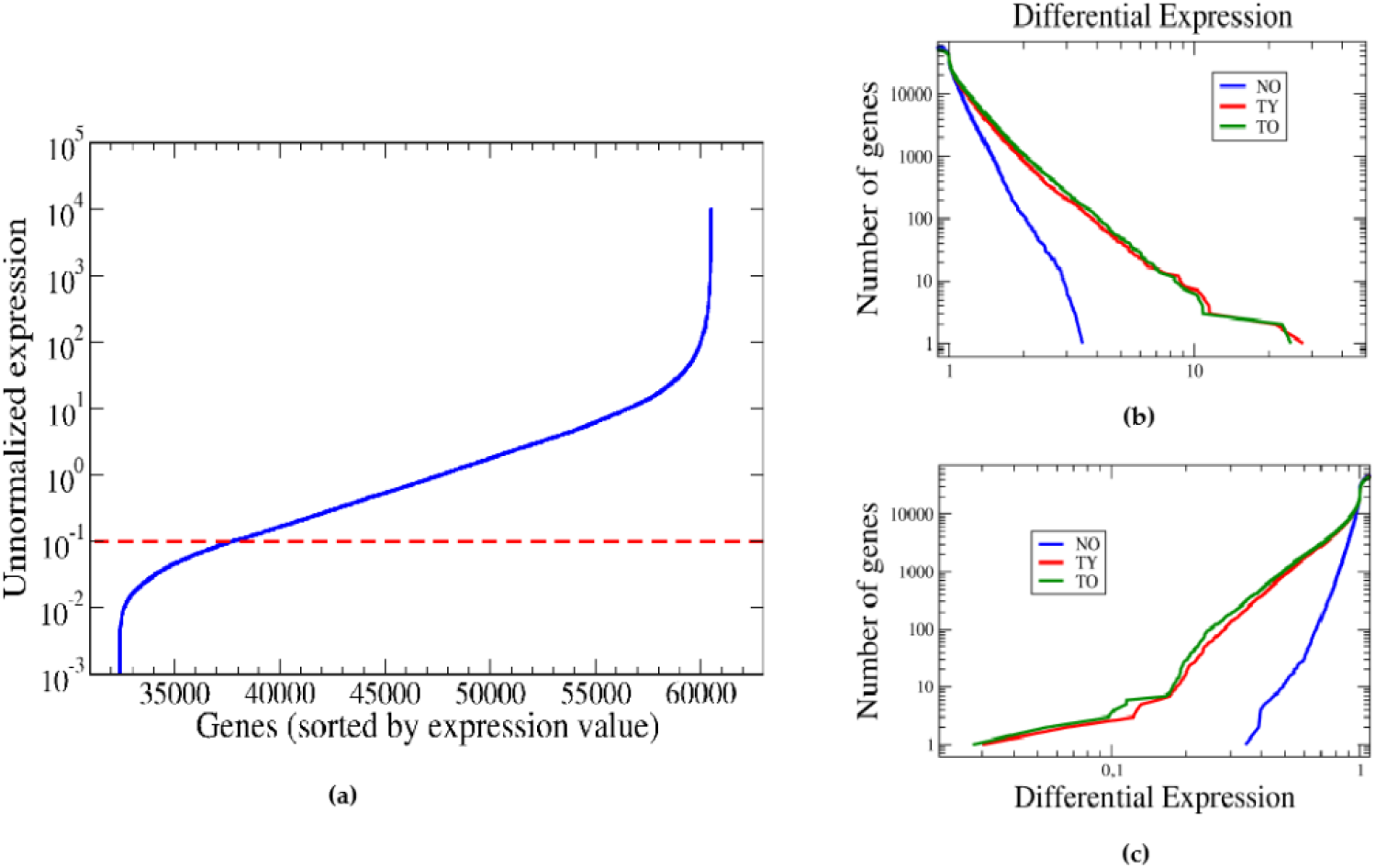
Un-normalized expression data and differential expression profiles in PRAD. (a) Typical range of (un-normalized) expression values from one representative patient (log scale). The red dashed line denotes the expression threshold for statistical significance. Genes with expression below the threshold both in normal tissue and in tumor are mapped to differential expressions very near one. (b,c) Differential-expression profiles for each of the data cohorts NO, TY and TO. The geometric average over the NY group is taken as reference. Notice that there are around 1000 genes with differential expression values below 1/2 (down-regulated) (b) and around 1000 genes with differential expression above 2 (up-regulated) (c). Notice also that the expression profiles practically coincide for the TY and TO groups, and apparently differ from the NO profile.

With these assumptions, the covariance matrix is written: σ2ij = Σ êi(s) êj(s) / (Nsamples-1), where the sum runs over the samples, s, and Nsamples is the total number of samples in the study. êi(s) is the fold variation of gene i in samples. The dimension of matrix σ2 is 60483, that is equals the number of genes in the data. By diagonalizing this matrix, we get the axes of maximal variance: The Principal Components (PCs). They are sorted in descending order of their contribution to the variance. As mentioned, PC1 captures 39% of the total data variance, PC2 11%, PC3 7%, etc. These results suggest that we may achieve a reasonable description of the main biological characteristics of PRAD using only a small number of the eigenvalues and eigenvectors of σ2. To this end, we diagonalize σ2 by means of a Lanczos routine in Python language, from which we get the first 100 eigenvalues and their corresponding eigen-vectors.

### Gene information and genome visualization

General gene information was collected from Genecards integrated data sources (www.genecards.org) including but not limited to expression, tissues-specificity, sub-cellular localization and diseases association data [23]. Genome visualizations were done with Ensembl release 100 - April 2020 (https://www.ensembl.org), Genome assembly: GRCh38.p13 (GCA_000001405.28) [24].

### LncRNA databases

To identify any previous association among identified LncRNAs and cancer, the following non-redundant databases were reviewed: Lnc2Cancer 2.0: An updated database that provides comprehensive experimentally supported associations between lncRNAs and human cancers [25]. LncRNADisease 2.0: contains experimentally and/or computationally supported data [26]. Cancer LncRNA Census (CLC): a compilation of 122 GENCODE lncRNAs with causal roles in cancer phenotypes [27]. The miRTarBase http://miRTarBase.mbc.nctu.edu.tw/ was used to uncover ceRNAs among selected LncRNAs [28].

### Enrichment analysis

The enrichment analysis was performed using the Enrich platform and the following categories: Ontologies (GO_Biological_Process_2018) and Pathways (Reactome_2016) [29].

### Cbioportal

Oncoprint visualizations for selected Genomic Profiles, Alteration Frequency, and mutations representation were obtained from Cbioportal, https://www.cbioportal.org/ [21,22].

### Cancer Driver repositories and driver prediction platforms

To search for any previous cancer association of identified genes the Cancer Gene Census and OncoKB (http://oncokb.org) databases were reviewed [30,31]. The driver prediction platforms IntoGene (https://www.intogen.org/search) and ExInAtor (https://www.gold-lab.org/cancer-driverlncrna-prediction-sof) were used to predict a potential driver role for protein-coding and non-coding genes [32,33].

### Pearson Correlation

Correlations among selected Pca clinical features and the PCs variables were performed using a Mathematica function (Pearson Correlation Test). A normal distribution of the variables is required.

## 2. Results

### 3.1. Data normalization surfaced an age-independent aberrant gene expression profile

In our analysis there are 52 samples of “normal” prostate tissues, 498 primary tumors samples, and one metastatic sample. RNA-seq data comprise expression values for 60483 independent genes, roughly 35000 of them are not transcribed at significant levels in prostate samples (Figure 1a).

Considering sample availability, we dicotomized the RNAseq data from “normal” and “neoplastic” tissues into two arbitrary age cohorts, with the “old” threshold set at ≥ 62 years (age range: 42-78, median=62) (Supplementary Figure 1). Thus, “normal” patient samples were divided in “young” samples (n=28, NY) and “old” patient samples (n=24, NO); whereas primary tumors samples were divided in “young” tumor samples (n=249, TY) and “old” ones (n=250, TO). While such distribution seems arbitrary and dictated by data availability, only 1 out 4 new PRAD diagnostic cases occurs below 60 years, whereas the mean diagnosis age is 66 years [34].

The normalization of expression values for each of the data cohorts TY and TO against NY group data indicates that the neoplastic transformation entails a similar and genome-wide over- and under-expression of genes, irrespective of the age of the patients (i.e., TY vs TO) (Figure 1b, c). Overall, we found roughly 1000 genes with normalized expression values above 2 and about the same number of genes with normalized expression values below 0.5.

### 3.2 Principal Component Analysis unveils a Core Expression Signature

The eigenvectors of the covariance matrix defined the PCs axes: PC1, PC2, etc., and projection over them define the new state variables. By definition, PC1 captures the highest fraction of the total variance in the sample set (i.e., PC1=39%), whereas the rest of components are sorted in descending order of their contribution to the variance 11% (PC2), 7% (PC3), 5% (PC4) and so on. Overall, the 8 first PCs comprised 74% of the data variance. Of note, 50% of data variance can be captured by the two major Principal Components (i.e., PC1 and PC2).

The PCA reveals that a Core Expression Signature composed of 33 genes from PC1 (hereafter, PRAD-CES33) can segregate primary neoplastic samples from normal prostatic tissues with roughly 4% and 8% of false positives and false negatives, respectively (Figure 2). Beyond such 33 genes, the addition of subsequent genes only slightly improves the ratio of false positives and the segregation of neoplastic from normal samples along the PC1 axis.

**Figure 2.**
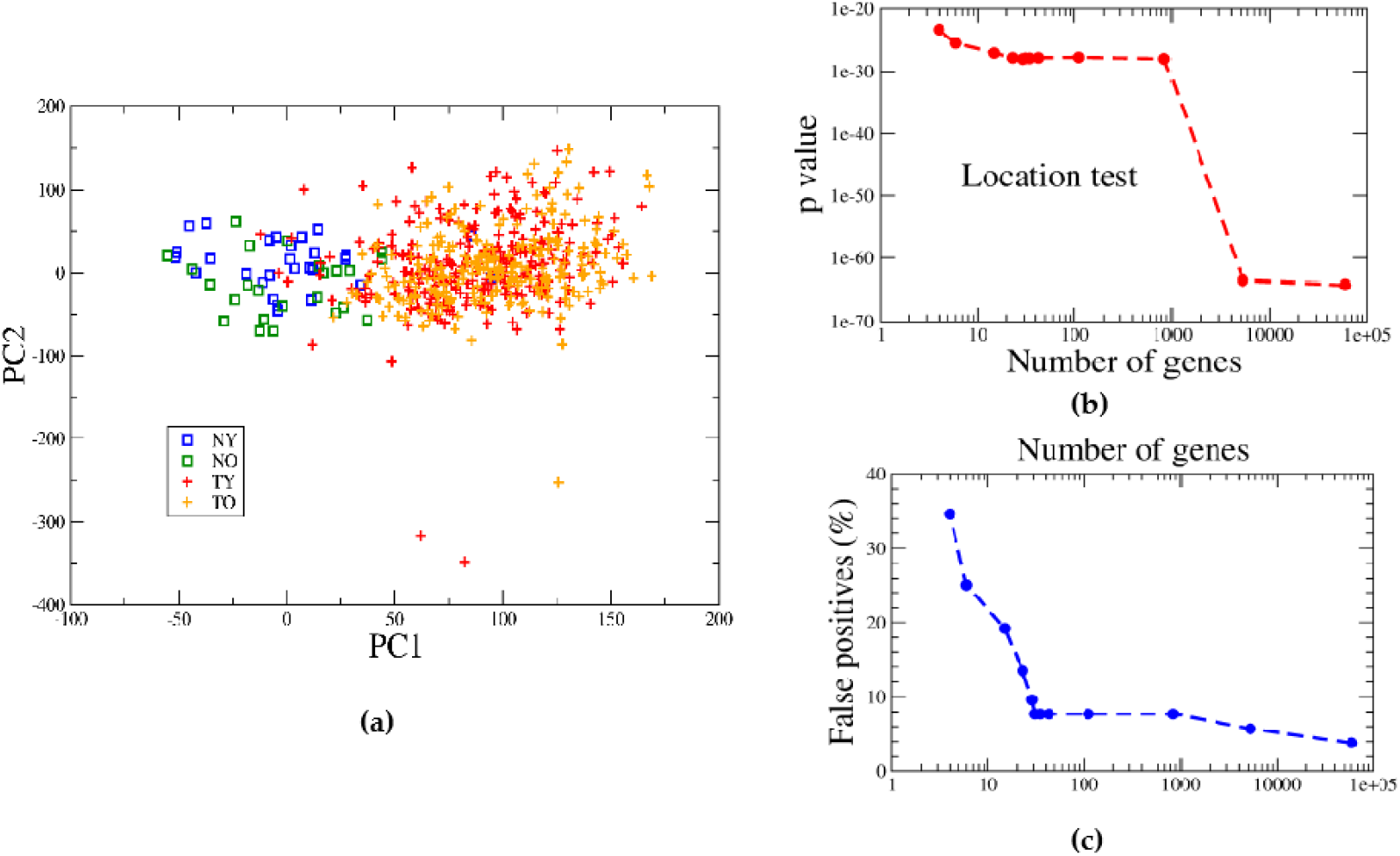
Principal Component Analysis (PCA) of RNAseq-based expression data from PRAD patients. (a) The tumor samples (cloud mean=+91.3) fall apart the distribution of “normal” ones (cloud mean= 0.0) along the PC1 axis defined here as the “cancer axis” (p-value = 10^-65, Mann-Whitney test). (b) Selecting a PC1 value of 45 as a frontier, 2/52 (3.8%) of normal samples are false positives, whereas 38/499 (7.7%) deemed as false negatives. (b, c) The optimal number of “core” genes within the PC1 gene subset is selected according to the ratio of False Positives and the Location Test.

The position along the PC1 axis of a sample is computed as x1 = Σ êi v1i, where v1i are the components of the unitary vector along this axis. A bardcode-like representation of the amplitudes for such 33 genes is represented in Supplementary Figure S2. The greatest value (i.e., over-expression) corresponds to a well know driver and biomarker gene in PRAD, the Prostate Cancer Associated 3 (PCA3) antisense [35,36]. Otherwise, the most under-expressed genes within this PRAD-CES are the protein coding gene SEMG1 [37]. Further bardcode-like analysis of top-100 genes contributing to PC1 axis shown a similar profile (Supplementary Figure S2). Detailed information about the 33 genes included in the core signature are described in Supplementary Table 1.

Notice that a picture like Figure 2b is drawn by recomputing the positions of samples along PC1, the ratio of false positives, etc. by using only the first n genes, ordered according to the module of their amplitudes in vector v1.

Finally, the distribution of tumor samples according to PRAD-CES on the PC1-PC2 plane was similar, irrespective of the age range (i.e., TY cloud median=87, TO cloud median=64) (Figure 2a). These results imply that not only the global normalized gene expression profile is similar among TY and TO in PRAD cases; rather, than a small number of core genes could become a molecular signature of the neoplastic state, irrespective of the age of the patient (i.e., PRAD-CES33).

### 3.3 Protein coding and RNA-genes compose the PRAD-CES33

The surfaced PRAD molecular signature its composed by protein coding (70%), as well as RNA-genes, including antiSense, pseudogene, and LncRNA (30%). The expression of the corresponding proteins was observed for 9/23 coding genes, whereas 6/10 RNA genes were detected in malignant prostate tissues (Supplementary Table 2). Of note, 20/23 (87%) protein coding genes have been previously associated to cancer, 18 (78%) particularly to Pca. Otherwise, 3/10 RNA genes have been connected to Pca (33%) (Supplementary Table 2).

PRAD-CES genes displayed low mutational burden with less than 5% of all samples having mutations (Supplementary Figure S3). Otherwise, roughly 15% of primary PRAD samples harbor CNV on PRAD-CES genes, being predominant deep deletions. The overall alteration frequency of PRAD-CES genes is roughly half of Pca driver genes annotated in the CGC (i.e., 21% vs 42% of cumulative alteration frequency, respectively).

### 3.4 Core expression signature includes emerging drivers and biomarkers

A text-mining indicated at least 18 surfaced genes may play driver roles in PRAD (Supplementary Table 2). However, only TP63 is enlisted in the CGC database as Tier 1 driver for NSCLC, HNSCC and DLBCL cancers, but not Pca. None of the remaining 23 protein coding genes populate CGC or OncoKB databases, nor two orthogonal driver prediction tools (i.e. IntoGene and ExInAtor) found further drivers among PRAD-CES genes.

Otherwise, we search for non-coding genes that might be predicted as drivers by ExInAtor. PCA3 was the only significantly mutated LncRNA predicted as a driver, despite four of the six LncRNAs were analyzed (i.e., PCA3, AP006748.1, AP001610.2, ARLNC1) (data not shown in details).

To further investigate RNA genes previously associated with cancer, we search three LncRNA databases Lnc2Cancer, LncRNADisease and Cancer LncRNA Census. Three (i.e., ARLNC1, PCA3, PCAT-14), six (i.e., AC092535.4, AP001610.2, AP002498.1, AP006748.1, PCA3 and PCAT14), and one RNA gene (i.e., PCA3), respectively; were previously associated with cancer (Table 1). Of note, the PRAD driver TMPRSS2 was predicted as mRNA target for AP001610.2 and AP006748.1 LncRNAs according to LncRNADisease database.

**Table 1.**
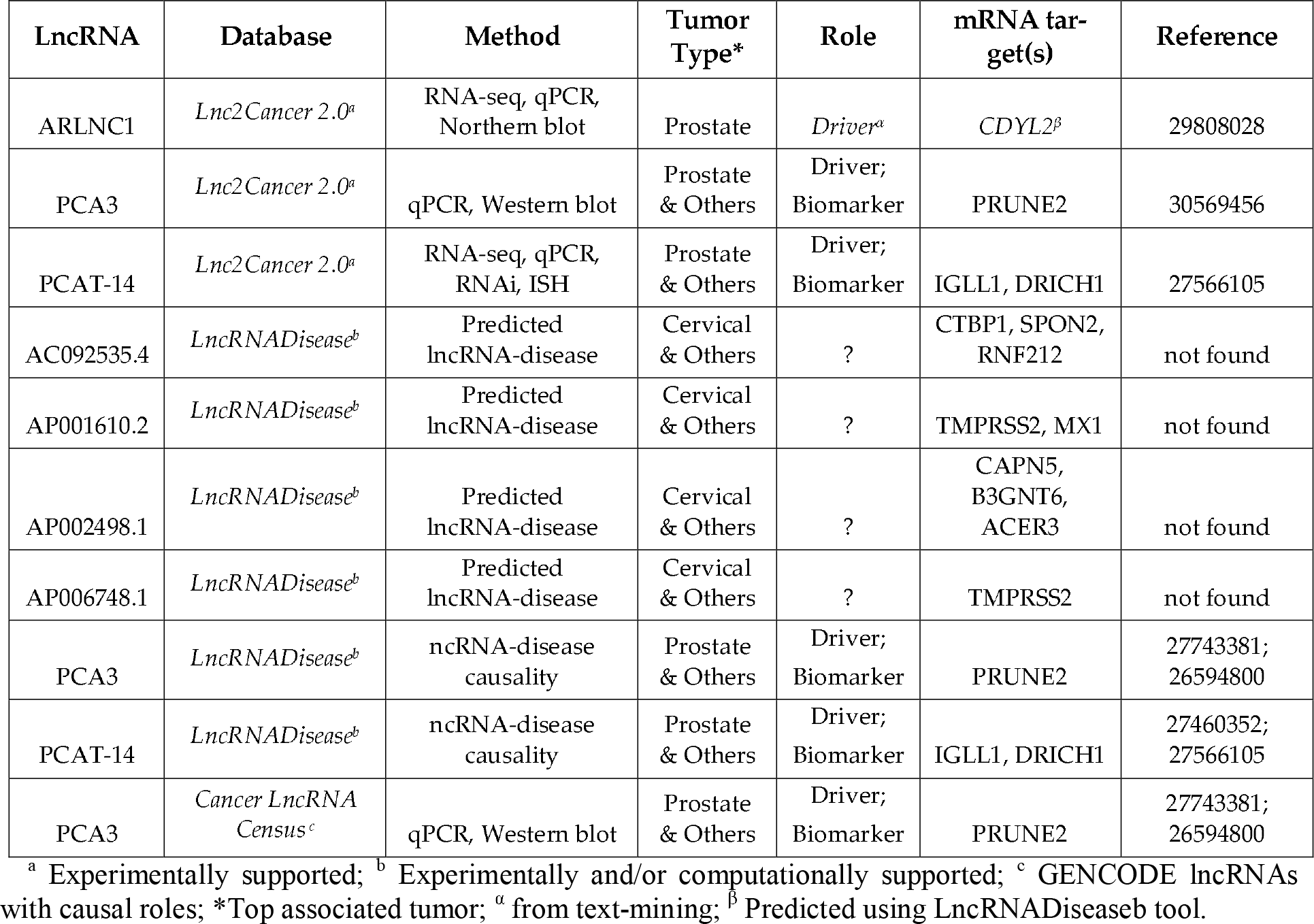
LncRNA included in PRAD-CES and their association with cancer according to indicated databases.

Finally, two others surfaced LncRNAs may impinge on Pca relevant genes according to a genomic inspection. The LncRNA AL359314.1 overlap with PCA3 and may reinforce the negative regulation of PCA3 over PRUNE2 [36]; whereas AC139783.1 is transcribed within the AMACR protein coding gene (Figure 4a, b).

**Figure 4.**
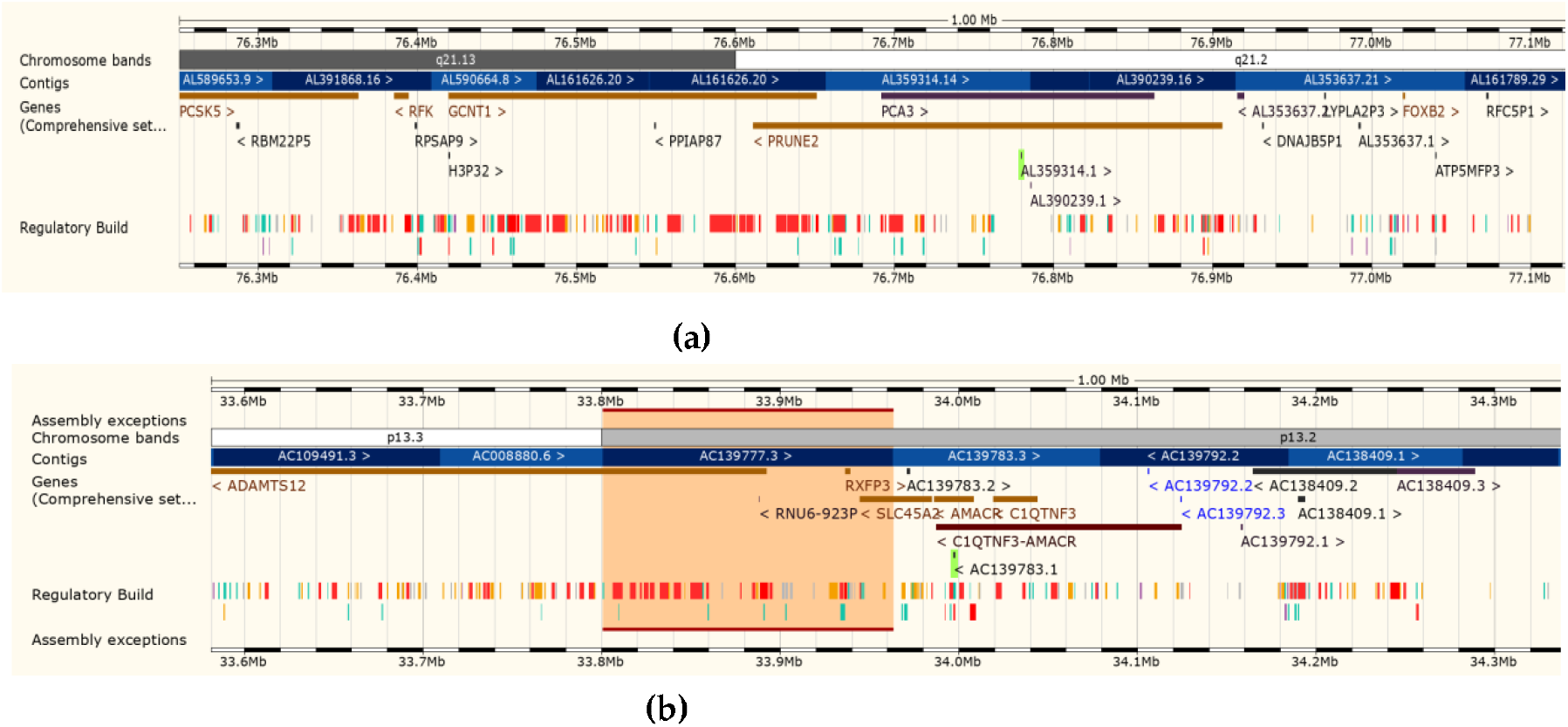
Genome location of the surfaced LncRNAs AL359314.1 (a) and AC139783.1 (b) indicating sequence overlap with protein coding genes PRUNE2 (anti-sense direction) and AMACR (sense direction). Representation of the regions of interest using Ensembl release 100.

### 3.5 Aberrant expression of PRAD-CES genes on independent datasets

The expression of PRAD-CES genes were further analyzed on three independent prostate cancers studies from Lapointe et al., 2004, Taylor et al., 2010 and Ross-Adams et al., 2015 [38–40]. Three putative emerging drivers in PRAD were consistently deregulated across the analyzed datasets. AMACR, SIM2 and GPX2 protein-coding genes were significantly up-regulated (AMACR, SIM2) or down-regulated (GPX2) in both primary and metastatic samples from lymph node or multiple sites (Figure 5, Supplementary Figure 4).

**Figure 5.**
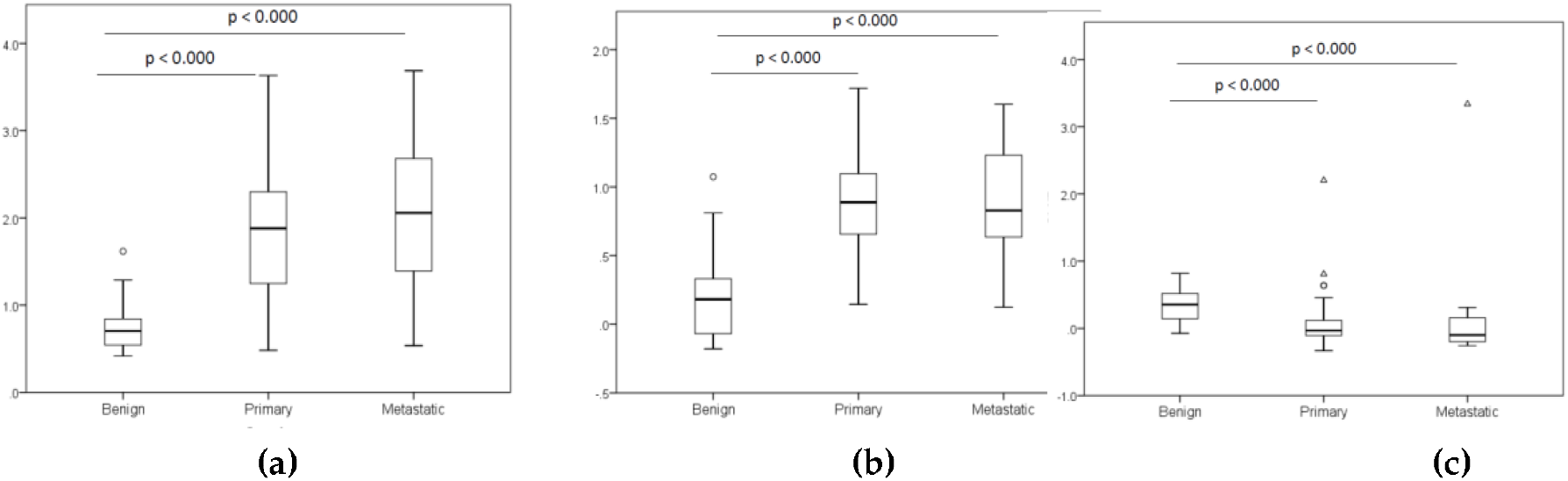
The figures shown Box plots of z-scores of Benign vs malignant tissues for AMACR (a), SIM2 (b) and GPX2 (c) genes. For statistics analysis a Kruskal-Wallis test with Bonferroni correction for multiple tests was conducted. Data taken from (Taylor et al., 2010)[40].

Overall, 14 of 33 PRAD-CES genes were included in the Lapointe dataset (Supplementary Figure 4). Whereas, the expression of 11 of them were consistently up- or down-regulated in this dataset, three showed no statistical differences (i.e., COMP, SEMG1 and SEMG2). On the other hand, in the Taylor dataset 17 of 33 PRAD-CES genes were detected. The expression of 11 genes were found consistently up- or down-regulated in primary tumors vs. benign tissues in agreement with our RNAseq-data, whereas no significant differences were found for 6 genes (i.e., GSTM1, SERPINA5, COMP, SLC39A2, SEMG1 and SEMG2). Finally, in the Ross-Adams dataset 19 of the 33 PRAD-CES genes were detected. The expression of 17 genes were found consistently up- or down-regulated in primary tumors vs. benign tissues, whereas no significant differences were found for 2 genes (i.e., SEMG1 and SEMG2) (Supplementary Figure 4).

### 3.6 PCs: Enriched Biological Processes and correlation with major clinical features

To seek for biological meanings beyond that of the individual genes populating the PCs, the top 33 genes from PC1, PC2 and PC3 were submitted to enrichment analysis to identify associated Biological Process. Of note, the top 33 genes populated PC1 (i.e., PRAD-CES) were mainly associated with tumor-intrinsic processes (GO:1900003, GO:0010950, GO:0007283, GO:0048232; p<0.01); whereas the Biological Process related to PC2 (GO:0006958, GO:0002455, GO:2000257, GO:0030449; p<0.001) and PC3 (GO:0050864, GO:0099024, GO:0051251, GO:0006911; p<0.001) suggested involvement of the Innate and adaptive Immune System (Figure 6a-c, Supplementary Table 3).

**Figure 6.**
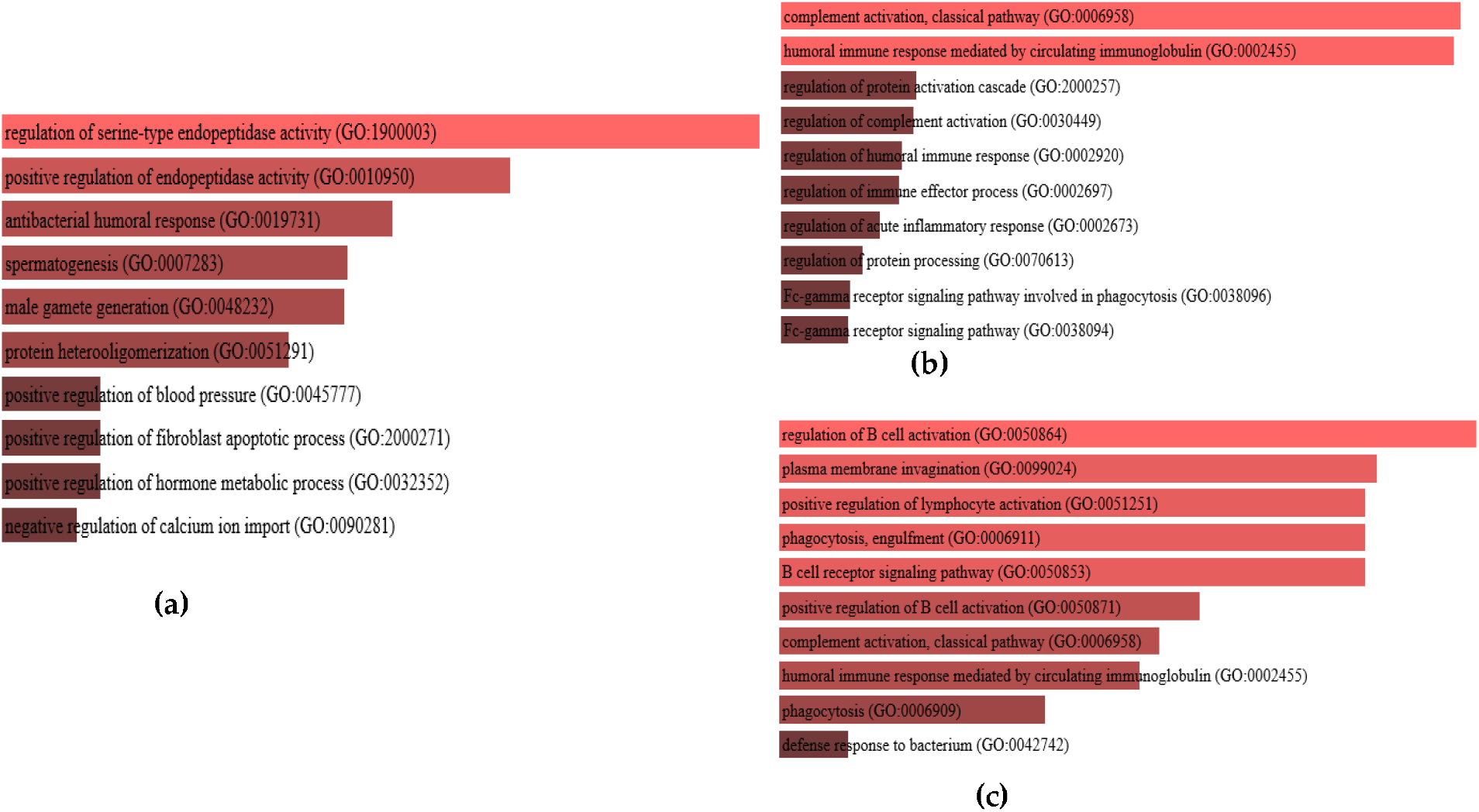
Enrichment analysis for Biological Process using the tool Enrich. PRAD_CES composite genes from PC1(a) and top 33 genes from PC2 (b) and PC3 (c) were included in the analysis. Statistical significance is in accordance with color from light (highly significant) to dark tones (less significant) (See Supplementary Table 3 for details).

Overall, the PRAD-CES genes (PC1) participate in more diverse BP and pathways compared to genes populated PC2 and PC3 (Supplementary Table 3 and 4). Otherwise, PC2 and PC3 populated genes seemed mainly involved in the complement activation, humoral immune response, regulation of B cell activation, phagocytosis, engulfment and regulation of acute inflammatory response.

To analyze the underlying distribution of major PRAD clinical features across PCs 1-3, a correlation analysis between each PC and the Gleason-Score, AR-Score, TMPRSS2-ERG and tumor cellularity were performed (Figure 7, Table 2).

**Figure 7.**
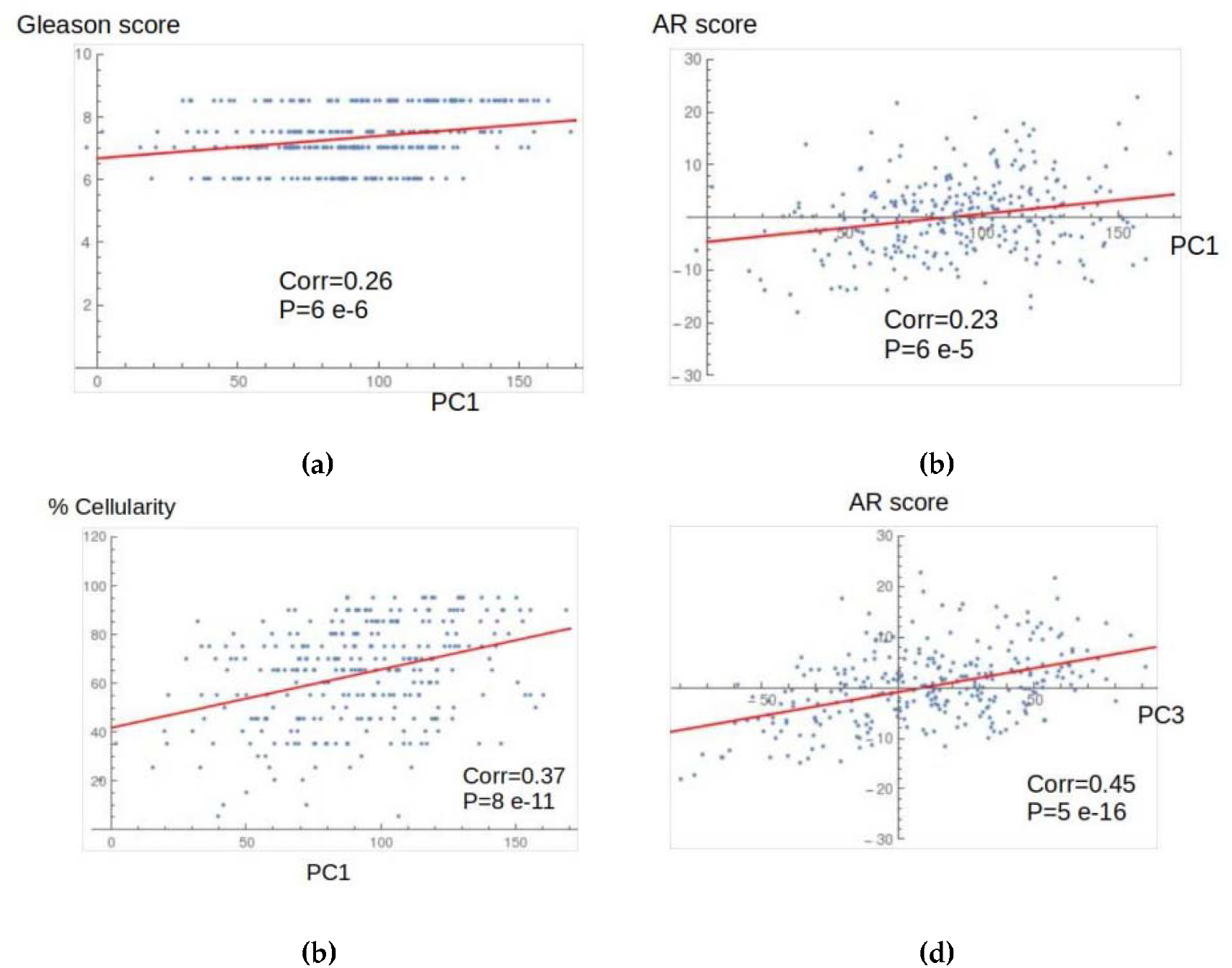
Correlation between PC1 and PC3 and major clinical features of PRAD using the data cohort Prostate Adenocarcinoma (TCGA, Cell 2015), comprising 333 primary tumors. Major features include Gleason-Score (a), AR-Score (b,d), TMPRSS2-ERG and tumor cellularity (c) (see Table 2 for details).

**Table 2.** Correlations among PCs and selected clinical features of PRAD (TCGA, Cell 2015). A Pearson Correlation Test was performed. **Bold** numbers indicate significant correlation p<0.05. See also Supplementary Table 5.

| Clinical Features | PC1 | PC2 | PC3 |
| --- | --- | --- | --- |
| Gleason Score | **0.26** | **-0.16** | 0.04 |
| TMPRSS2-ERG | 0.02 | **-0.18** | **0.24** |
| AR Score | **0.23** | **0.32** | **0.45** |
| Cellularity | **0.37** | **0.14** | **0.19** |

Our analysis revealed that PC1 values shown a weak-yet positive correlation with Gleason (R<0.30, p=6.0E-06), and AR Score (R<0.30, p=5.0E-5); whereas a medium-strength positive association with Tumor cellularity (R=0.37, p=8.0E-11) was seen. Of note, independent correlations among clinical features in this dataset indicated that the Gleason score weakly correlates with Cellularity (R=0.26, p=8.0E-6) and TMPRSS2-ERG fusion anti-correlates with AR Score (R=−0.24, p=4.0E-5) (Supplementary Table 5). Therefore, the observed correlation between PC1 values and the above-mentioned clinical features may reflect the underlying PRAD biology which is in line with the fact that PC1 explain up to 39% of data complexity, being a more “general” expression signature.

Concerning PC2, we observed an anti-correlation among TMPRSS2-ERG and AR Score which goes along the underlying PRAD biology; however, in this PC the Gleason Score anti-correlated with Tumor Cellularity. Finally, the genes included in PC3 showed positive correlations with TMPRSS2-ERG, AR Score and Tumor Cellularity (Table 2, Supplementary Table 5).

## 3. Discussion

Here, we use Principal Component Analysis (PCA) to surface a gene expression signature which may “describe” primary PRAD, providing new putative biomarkers and/or molecular targets to intervene. Such dimensionality reduction algorithm clearly segregates tumor from normal samples, with eight PCs capturing roughly 3/4 of data complexity. The RNA-seq input data was obtained from the Prostate Adenocarcinoma cohort TCGA_Firehose Legacy, which comprised a significant number of tumor and normal samples, the ultimate required to perform our custom-made normalization. Furthermore, considering that PCA lose resolution on highly heterogeneous and pooled data, we selected only this Pca data cohort to perform our PCA [13].

Our custom-made normalization revealed a long-tail distribution of expression values which might reflect global deregulation events associated with aging and/or malignant transformation [41]. Since we used “Normal Young” data as reference, the obtained pattern may suggest that neoplastic transformation over-impose on already age-adjusted global expression profile (i.e., similar TY and TO distribution). However, this notion needs to be verified by using larger and better dichotomized age-based patient cohorts. Of note, the observed long-tail distribution is independent of the type of expression data (i.e. RNA-seq), since similar global gene-expression patterns emerged after analyzing micro-array data using our normalization procedure (data not shown).

The PCA allow us to identify a Core-Expression Signature (PRAD-CES) composed of 33 genes which accounts for 39% of data variance along what we call the cancer axis (PC1). The biological meaning of PC2 and PC3 seems more elusive, accounting for an additional 18% of variability. The PRAD-CES includes validated, emerging and putative PRAD drivers and/or biomarkers. Although only one validated protein-coding driver was found (i.e., TP63), three RNA genes with causative roles were surfaced: ARLNC1, PCA3, and PCAT-14 [36,42–44]. Otherwise, six protein coding genes awaits further validation concerning PRAD driver roles: OR51E2, HPN, AMACR, DLX1, HOXC6 and WFDC2 [45–50]. Concerning potential or validated biomarkers, the PRAD-CES list contains 15 RNA- or protein-coding genes with such a role. Among them HOXC6, TDRD1, and DLX1 have been already proposed to identify patients with aggressive prostate cancer [51]. TDRD1 might also play an important role in prostate cancer development, and as a cancer/testis antigen, a potential therapeutic target for cancer immunotherapy [52].

Of note, cross-validation of PRAD-CES genes using independent data cohorts (i.e., Lapointe et al., 2004, Taylor et al., 2010 and Ross-Adams et al., 2015), indicated that most of these genes were consistently deregulated in primary PRAD, with notable exceptions on comp, semg1 and semg2 genes. Otherwise, the expression of 14 PRAD-CES genes could not be verified in all datasets [38–40]. Overall, the most consistent genes among those detected across all analyzed data were OR51E2, SIM2, HPN, SLC45A2, TDRD1, PCA3, DLX1, AMACR, WFDC2, and HOXC6.

On the other hand, our PCA surfaced nine over-expressed RNA genes, six of them lacking previous association with Pca. Particularly, four LncRNAs could target PRAD driver’s genes TMPRSS2, PRUNE2 and AMACR. One interesting finding was the genome proximity/overlap among PRAD-CES over-expressed genes AC139783.1, AMACR and SLC45A2 on Chromosome 5. SLC45A2-AMACR was reported as a novel fusion protein which is associated with progressive Pca disease [53]. Otherwise, among several miRNAs which may down-regulate AMACR expression in Pca, the potential sponging of hsa-miR-26a-5p by the surfaced AC139783.1, needs to be addressed. AMACR over-expression have been associated with Pca evolution towards hormone-independency, whereas AMACR inhibition seems a feasible strategy to treat hormone-refractory prostate cancer patients [47]. LncRNA over-expression in Pca has been related with disease progression, used as prognostic factor, or proposed as therapeutic targets [54–56].

The most frequent molecular abnormalities in PRAD involved gene-fusions, copy-number alterations and epigenetic deregulation [14]. As a matter of facts, the mutational burden observed in surfaced PRAD-CES genes was low, suggesting that expression levels and not co-existing mutations determine the PCA-based segregation of tumor from normal samples. Furthermore, less than 3% of PRAD samples included in our study displayed CNV, thus suggesting that most of the observed gene expression deregulation arose from epigenetic and/or other transcription-based mechanism.

Finally, we selected four Pca molecular/clinical features to correlate with PCs 1-3. The first, Gleason score, remains as a cornerstone pathological criterion for risk-stratification and prognosis [59]. Furthermore, primary prostate cancer is androgen dependent, and androgen-mediated signaling is crucial in prostate cancer pathogenesis, driving the creation and over-expression of most ETS fusion genes [57,58]. Among such ETS fusion genes, TMPRSS2-ERG fusion accounts for 46% of cases [14]. The fourth clinical feature, i.e. tumor cellularity, was used here as a proxy for non-prostatic yet-relevant infiltrating populations [60]. The observed correlations indicated PC1 reflects the underlying primary PRAD biology with positive correlation among Gleason Score and Tumor Cellularity, as well as among this variable and AR Score. Otherwise, genes comprising PC2 and PC3 might reveal a transition towards a more aggressive and inflammation-prone phenotype, with a mixture of tumor epithelial cells and infiltrating immune cells [61]. This notion seems also supported by a weaker correlation of PC2 and PC3 genes with tumor cellularity, but also by the increasingly positive correlation among genes populating PCs 1-3 and the AR Score (i.e., from 0.23 to 0.45). Of note, only PC3 genes positively correlated with TMPRSS2-ERG fusion. Altogether, an intriguing possibility is whether PC3-populating genes may describe an inherent fraction of highly infiltrated tumor cells endowed to metastasize.

Overall, our study is limited by data availability/structure and biopsy bias as any global transcriptome inquire [62]. Primary prostate tumors are multi-focal and molecularly heterogeneous; thus, the surfaced gene expression signature may “described” only the sampled site, which also contains different cells from the tumor micro-environment [63,64]. However, our PCA indeed uncover relevant PRAD genes found dispersed across several studies, providing new putative biomarkers and/or drivers. In this sense, the inclusion of PCA3 within our PRAD-CES seems encouraging since this LncRNA is well recognized as causative, prostate-specific and feasible biomarker which is secreted to an easy-to-inquire biological fluids [65]. Finally, as therapeutic options for poorly tractable Pca are limited, the evaluation of putative novel molecular targets populating PRAD-CES seems appealing.

## Supplementary Materials

Figure S1: Histogram showing the age range for patients included in the PRAD-TCGA Firehose Legacy cohort used for PCA of RNA-seq expression data, Figure S2: A Barcode-like representation of PRAD genes comprising the unitary vectors along PC1. Top panel, contains 33 genes identified for a delta=0.027; Lower panel representing 100 genes for a delta=0.0235. Within the barcode the major value (i.e., over-expression) corresponds to PCA3, whereas the lower value (i.e., under-expression) belongs to SEMG1, Figure S3: Differential expression of PRAD-CES genes in the MSKCC, Cancer Cell 2010 cohort. The oncoprint representation tool from Cbioportal is used. Z scores>2, normal vs tumor expression values, Figure S4: Expression analysis of genes from the PRAD-CES using three independent data cohorts of primary prostate tumors from radical prostatectomy, Table S1: Detailed information about the 33 genes included in the PRAD core signature (PRAD-CES33), Table S2: Gene Classification, Disease Association and Gene Product Expression, Table S3: Enrichment analysis for Biological Process using the tool Enrich, Table S4: Enrichment analysis for Biological Pathways using the tool Enrich, Table S5: Correlations among PCs and selected clinical features of PRAD (TCGA, Cell 2015).

## Author Contributions

Conceptualization, all authors; methodology, A.G. and Y.P.; software, A.G.; formal analysis, A.G. and Y.P.; writing—original draft preparation, Y.P.; writing—review and editing, all authors; funding acquisition R.P. “All authors have read and agreed to the published version of the manuscript.”

## Data Availability Statement

The data come from the TCGA Research Network: https://www.cancer.gov/tcga and Cbioportal: https://www.cbioportal.org/ as for March, 2019.

## Acknowledgments

We would like to thanks to Dr. Simone Chevalier from the Centre for Translational Medicine, Research Institute, McGill University for expression data cross-validation. A.G acknowledges the Cuban Program for Basic Sciences and the Office of External Activities of the Abdus Salam Centre for Theoretical Physics for support.

