## Supplementary figures and images for "Principal component analysis of RNA-seq data unveils a novel prostate cancer-associated gene expression signature"

### Supplementary Fig 1

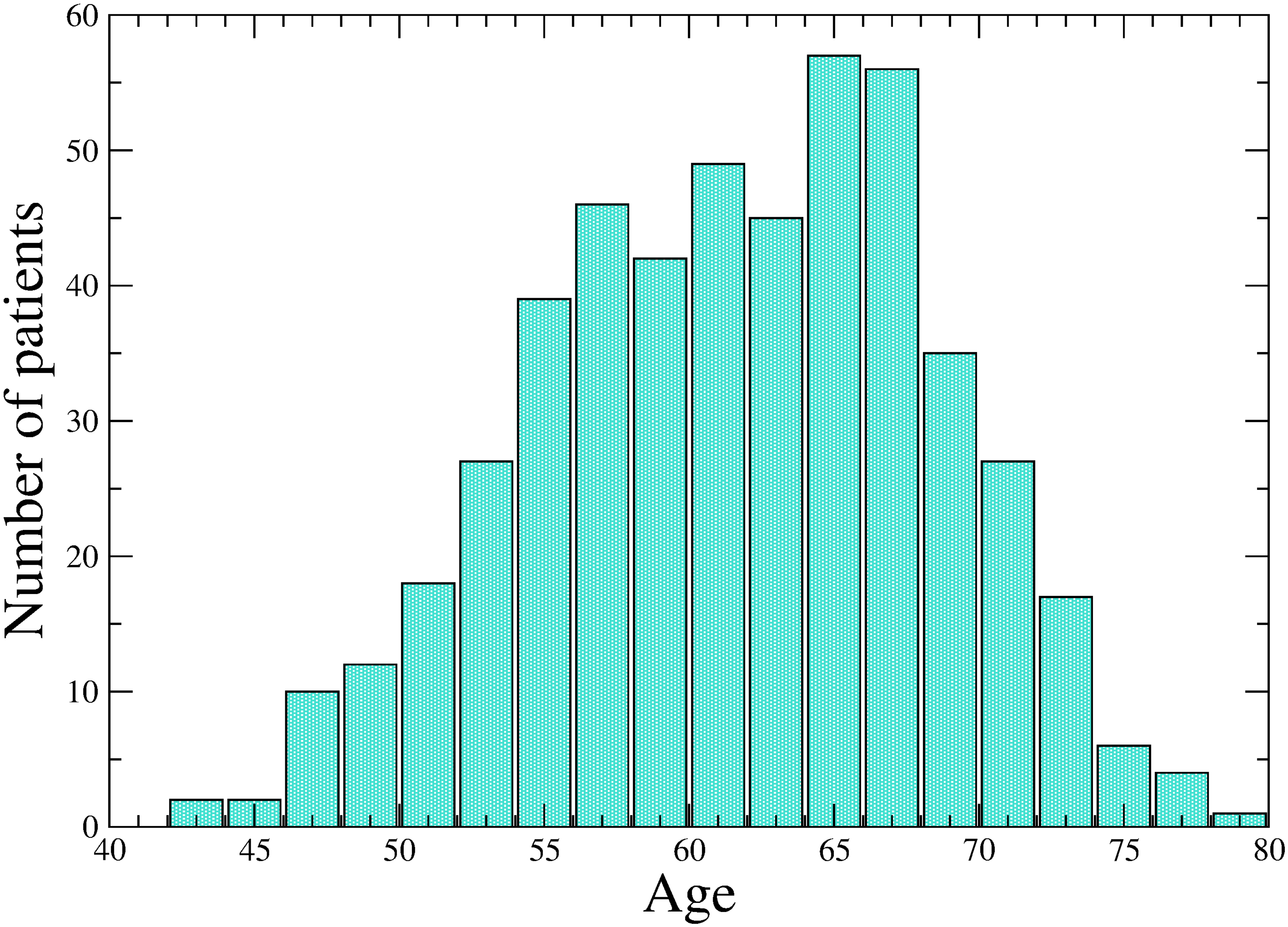

### Supplementary Fig 2

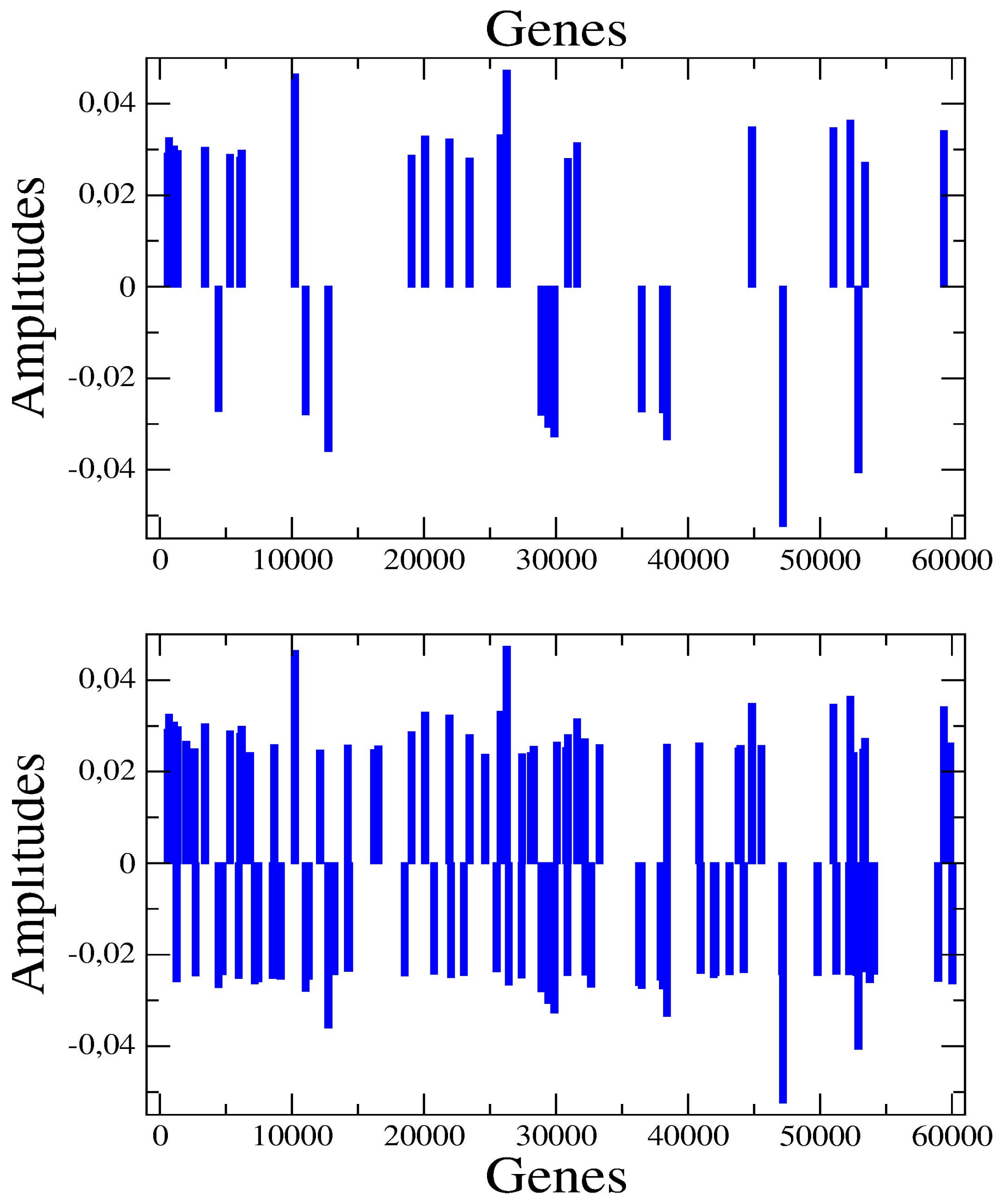
