## Supplementary Fig 3 for "Principal component analysis of RNA-seq data unveils a novel prostate cancer-associated gene expression signature"

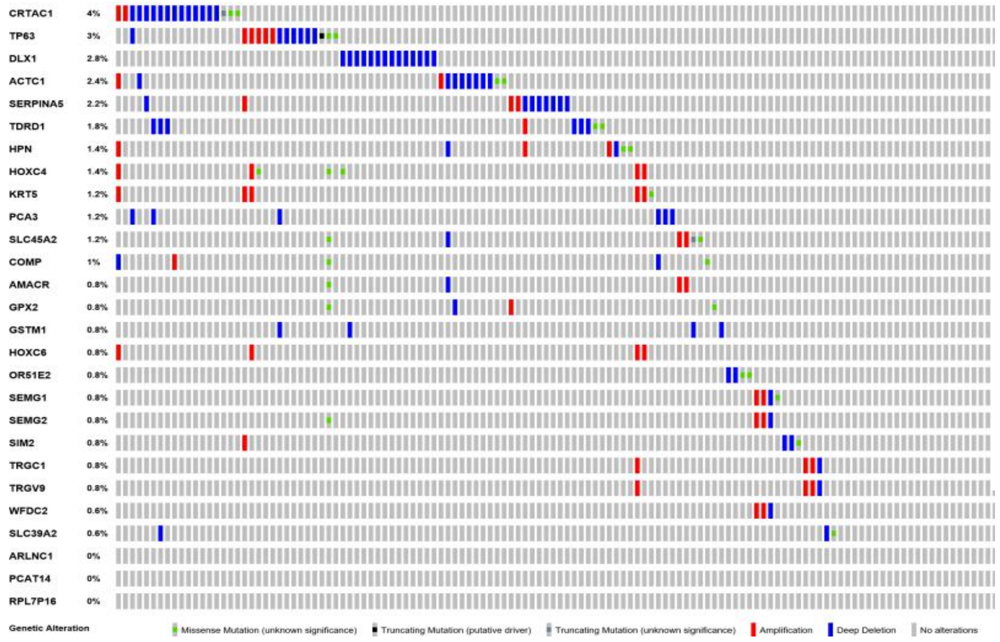


**(a)**

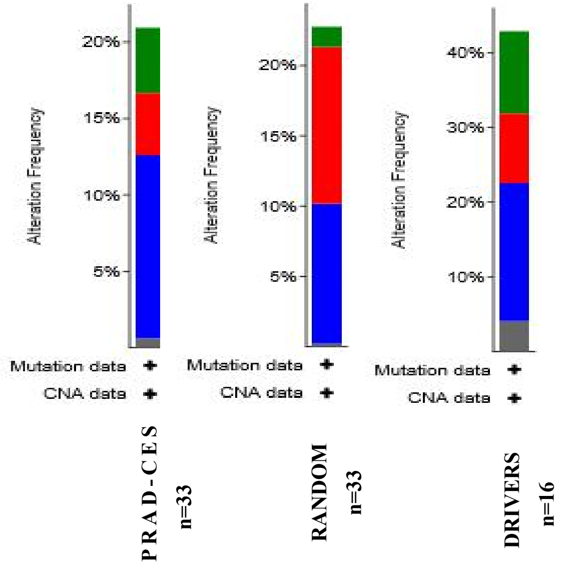


**(b)**

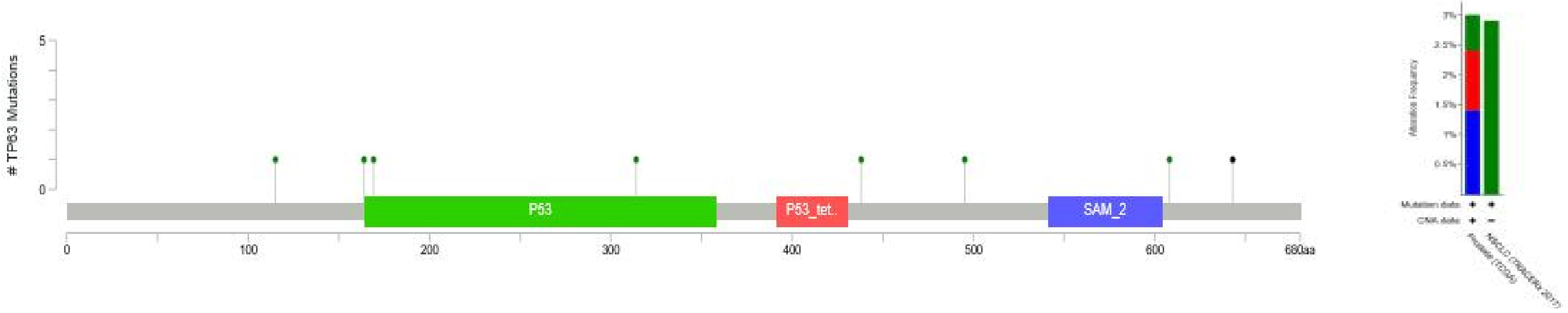


| Study | Sample ID ▼ | Cancer Type | Protein Change | Annotation | Functional Impact | Mutation Type | Copy # | COSMIC | # Mut in Sample |
|---|---|---|---|---|---|---|---|---|---|
| Prostate Adenoca... | TCGA-XK-AAIW-01 | Prostate Adenocarcinoma | *A164V* | | | Missense | Diploid | | 6525 |
| Prostate Adenoca... | TCGA-J4-A83J-01 | Prostate Adenocarcinoma | *R643** | | | Nonsense | ShallowDel | | 47 |
| Prostate Adenoca... | TCGA-HC-A6AP-01 | Prostate Adenocarcinoma | *G314A* | | | Missense | Gain | 1 | 46 |
| Non-Small Cell L... | CRUK0091-R2 | Lung Squamous Cell Carcinoma | *Q438H* | | | Missense | | | 418 |

**(c)**
