## Supplementary Fig 4 for "Principal component analysis of RNA-seq data unveils a novel prostate cancer-associated gene expression signature"

### Expression of genes of your list in reported primary prostate tumors of radical prostatectomy patients [1,2,3]


1. **Lapointe** J, Li C, Higgins JP, van de Rijn M et al. Gene expression profiling identifies clinically relevant subtypes of prostate cancer. *Proc Natl Acad Sci U S A* 2004 Jan 20;101(3):811-6.

2. **Ross-Adams** H, Lamb AD, Dunning MJ, et al. Integration of copy number and transcriptomics provides risk stratification in prostate cancer: A discovery and validation cohort study. EBioMedicine. 2015;2(9):1133–1144.

3. **Taylor** BS, Schultz N, Hieronymus H, et al. Integrative genomic profiling of human prostate cancer. Cancer Cell. 2010;18(1):11–22.

### 1- Lapointe dataset

Benign vs. primary tumour vs. lymph node metastases

Box plots of z-scores vs. benign, with genes presented in the same order for all analyses

Statistics: Kruskal-Wallis test with Bonferroni correction for multiple tests

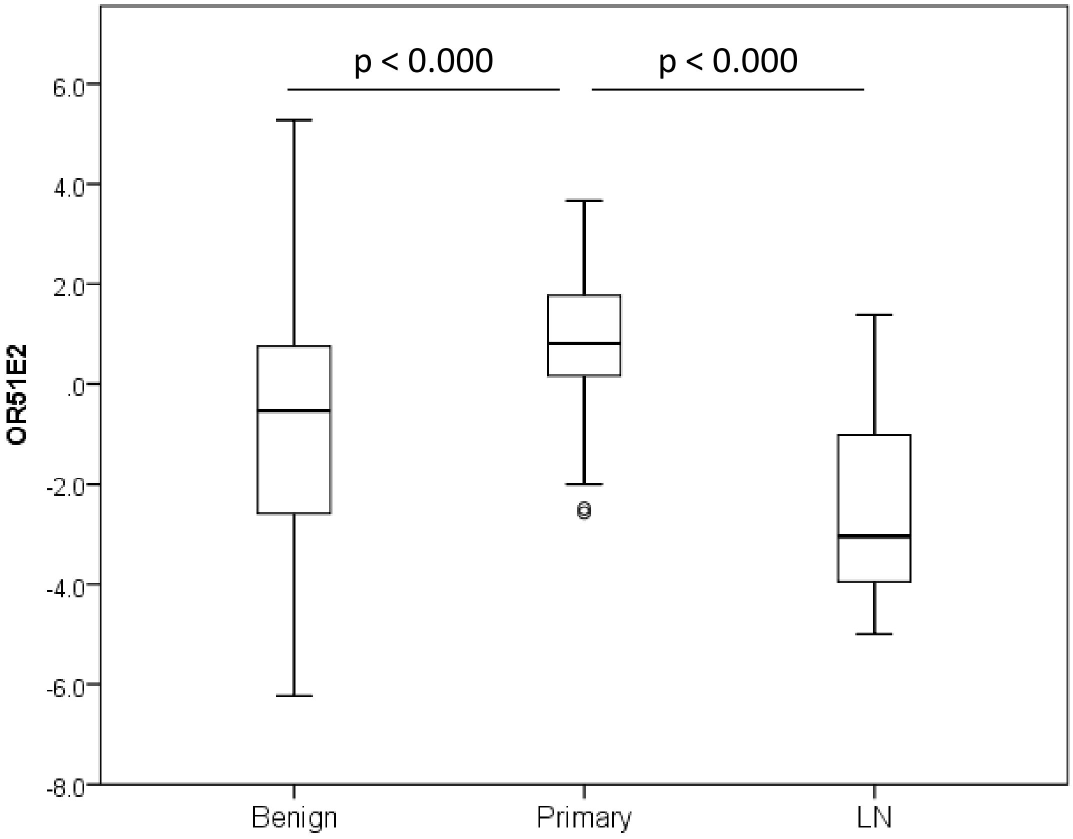
p < 0.000
p < 0.000
6.0
4.0
2.0
.0
-2.0
-4.0
-6.0
-8.0
OR51E2
Benign
Primary
LN

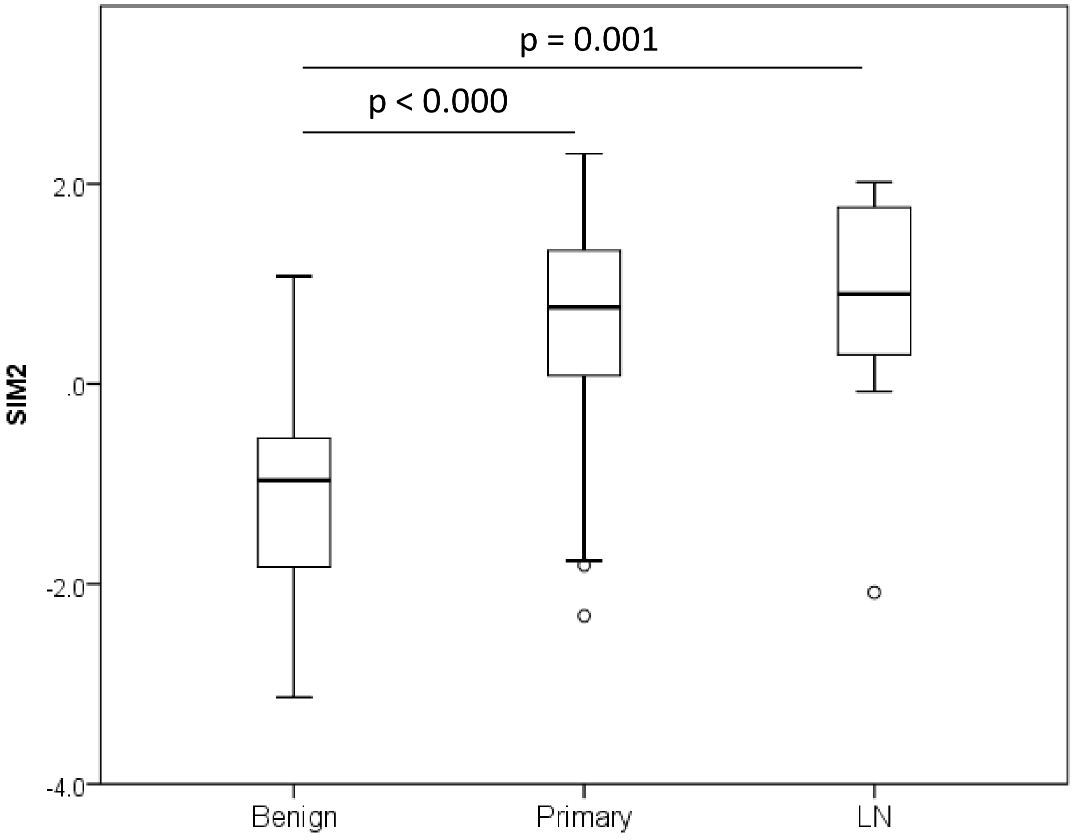
p = 0.001
p < 0.000
2.0
.0
-2.0
-4.0
SIM2
Benign
Primary
LN

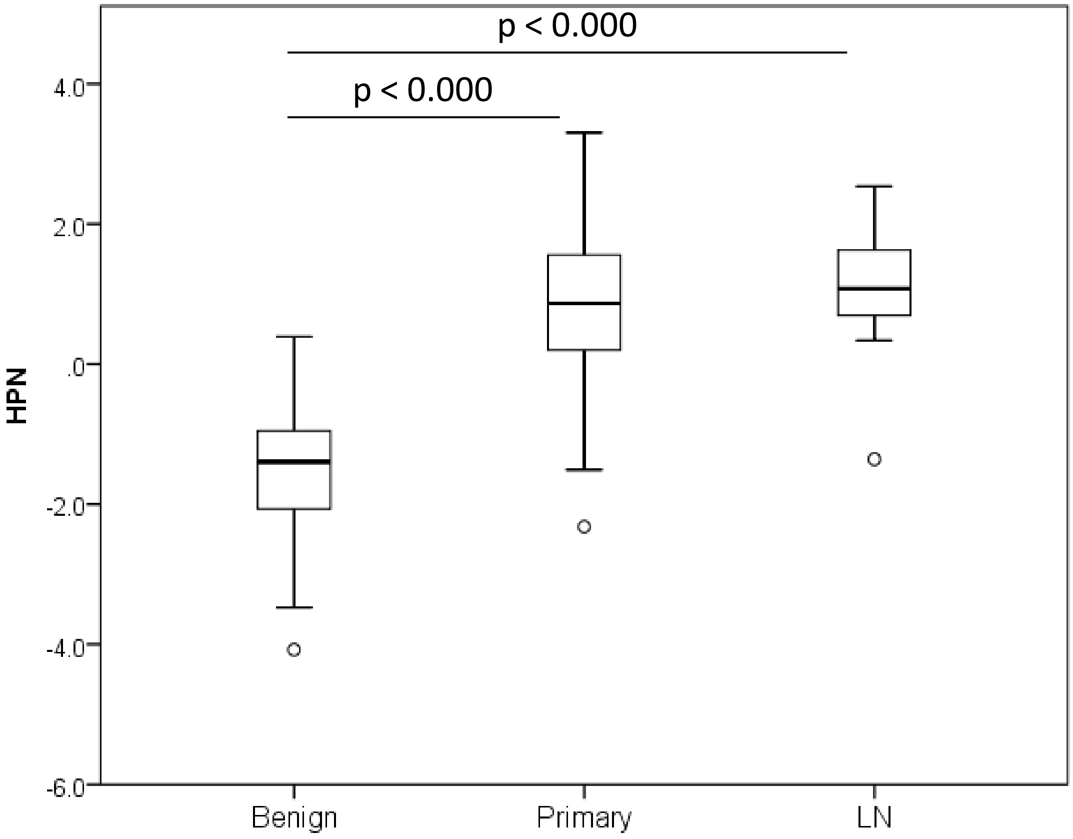
p < 0.000
p < 0.000
4.0
2.0
.0
-2.0
-4.0
-6.0
HPN
Benign
Primary
LN

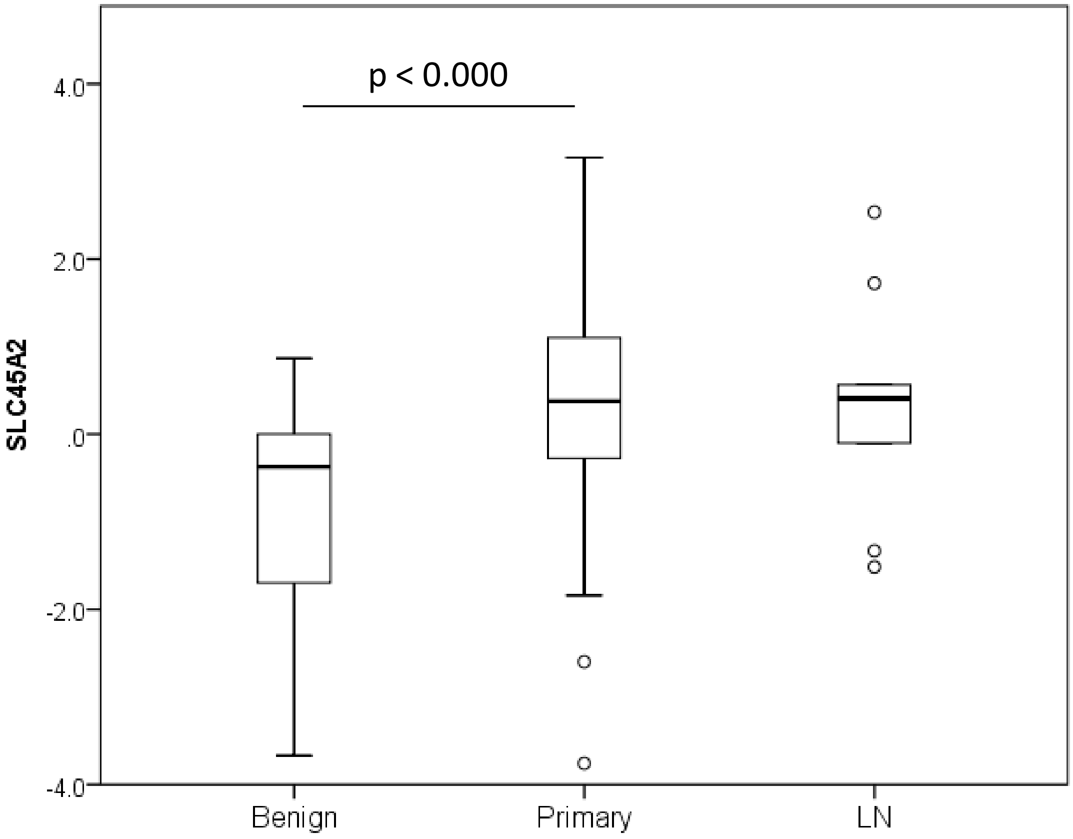
p < 0.000
4.0
2.0
.0
-2.0
-4.0
SLC45A2
Benign
Primary
LN

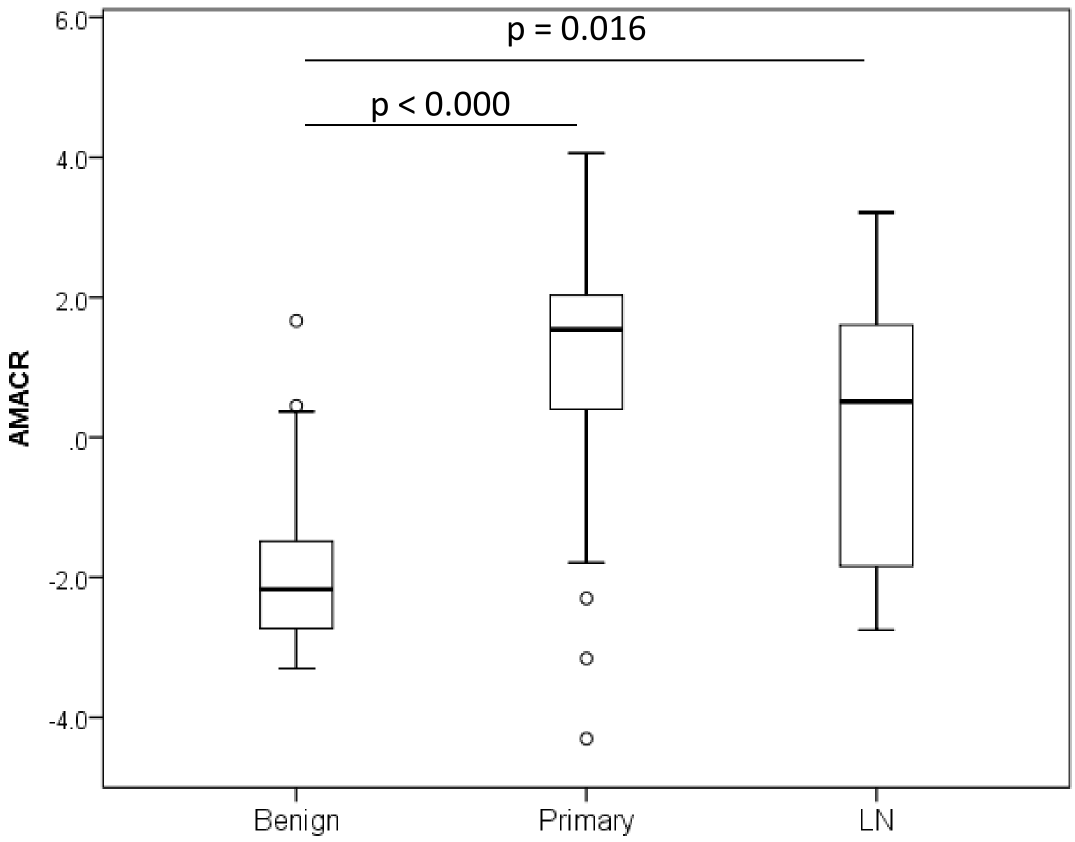

p = 0.016
p < 0.000
AMACR
6.0
4.0
2.0
.0
-2.0
-4.0
Benign
Primary
LN


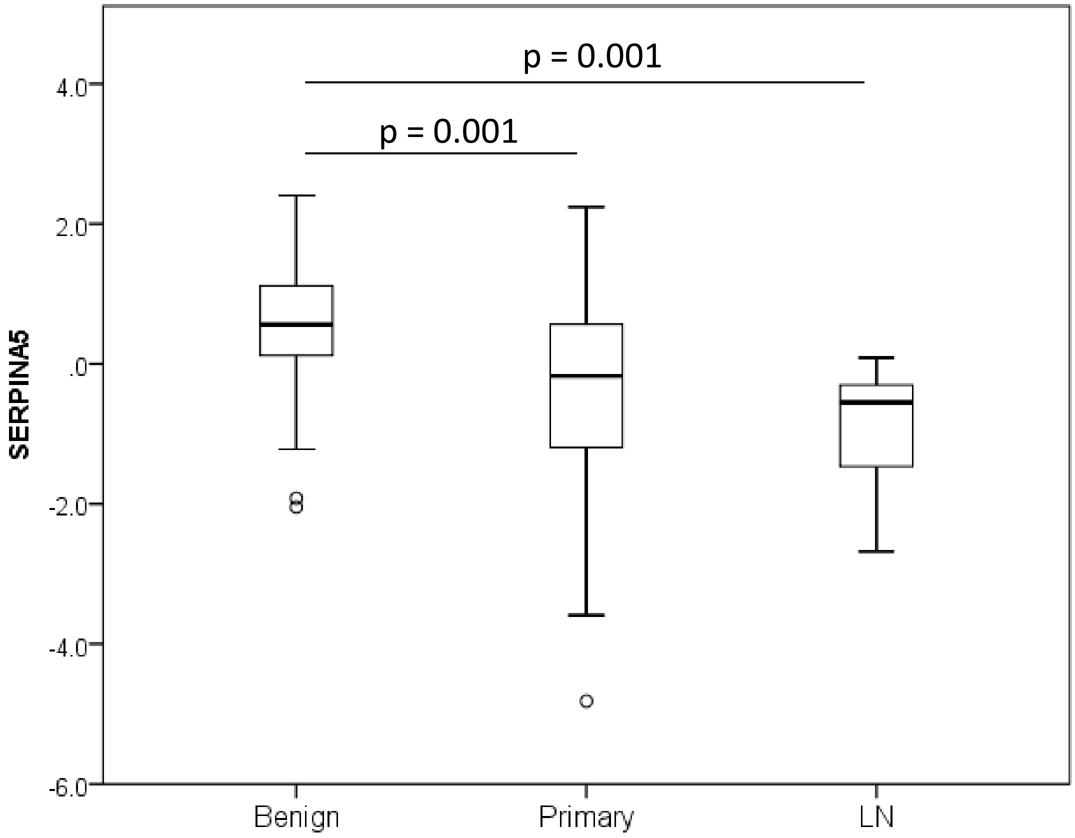

p = 0.001
p = 0.001
SERPINA5
4.0
2.0
.0
-2.0
-4.0
-6.0
Benign
Primary
LN

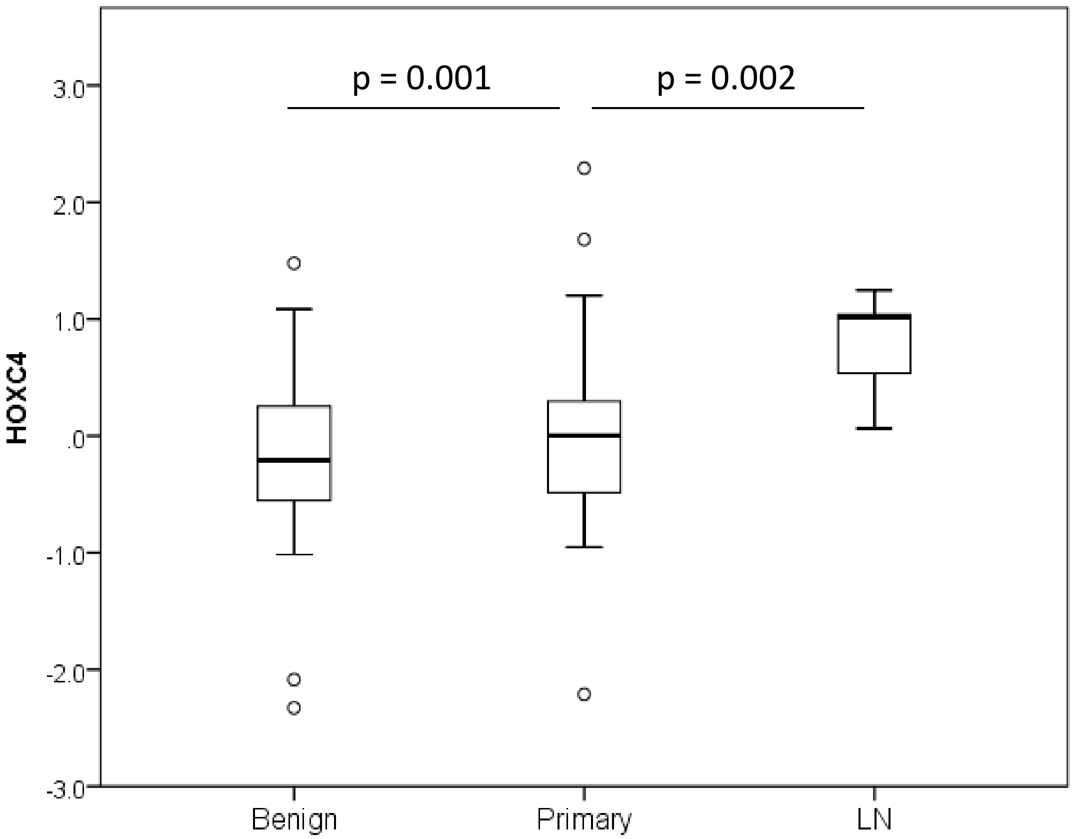
p = 0.001
p = 0.002
3.0
2.0
1.0
.0
-1.0
-2.0
-3.0
HOXC4
Benign
Primary
LN

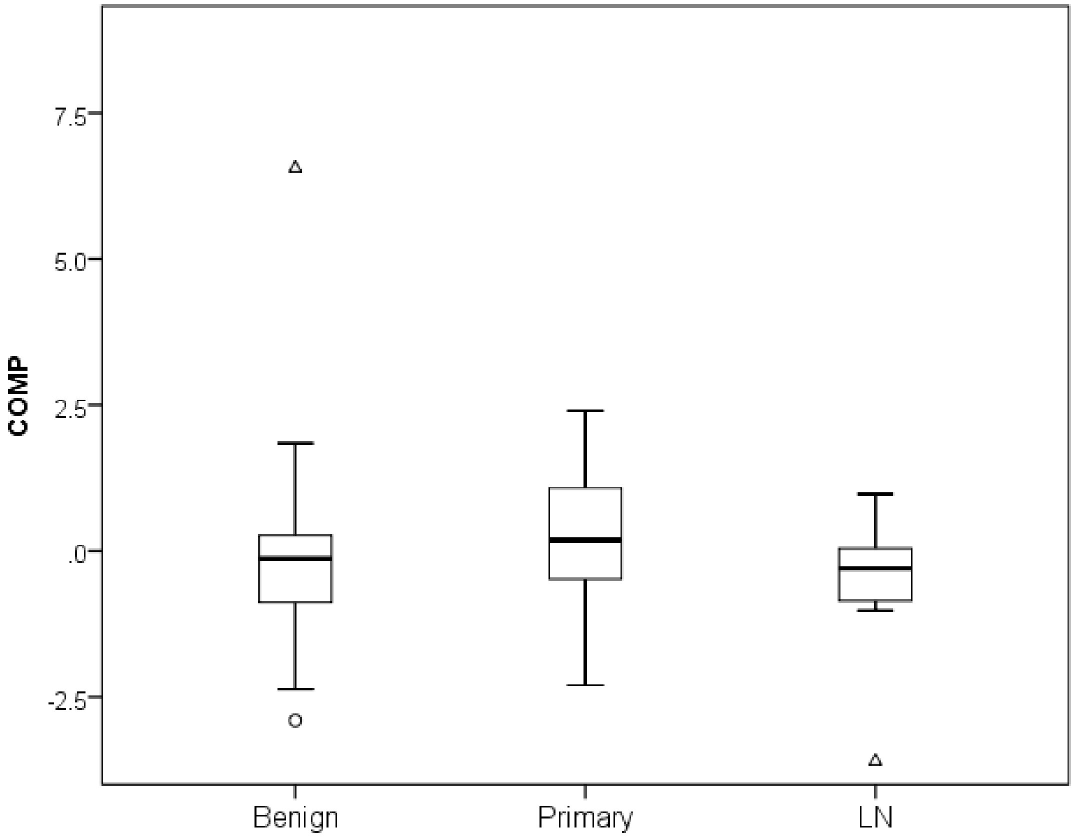
7.5
5.0
2.5
.0
-2.5
COMP
Benign
Primary
LN

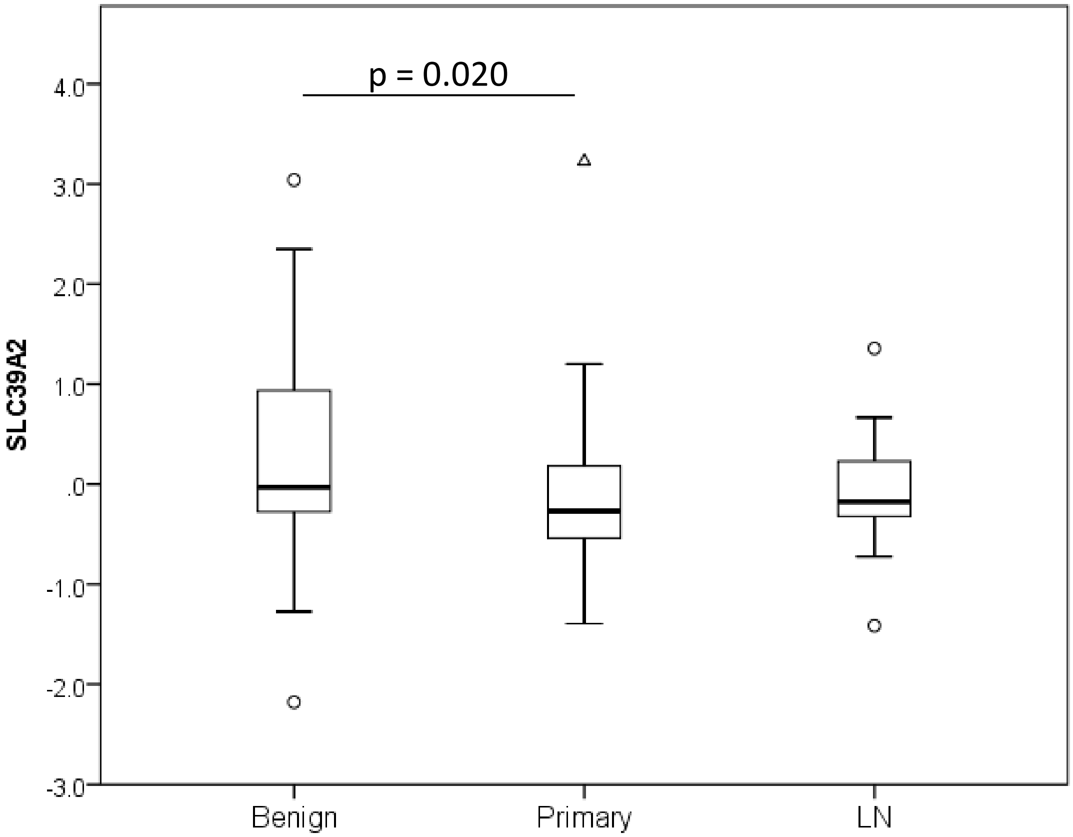
p = 0.020
SLC39A2
4.0
3.0
2.0
1.0
.0
-1.0
-2.0
-3.0
Benign
Primary
LN

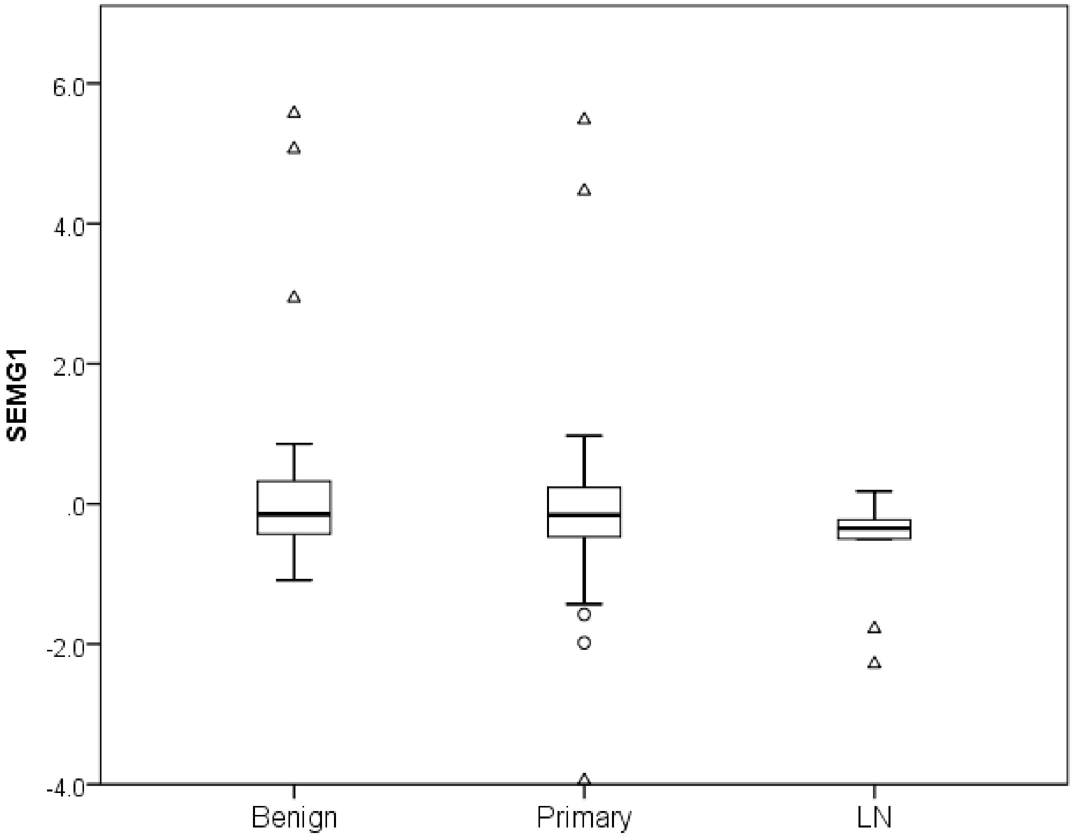
SEMG1
6.0
4.0
2.0
.0
-2.0
-4.0
Benign
Primary
LN

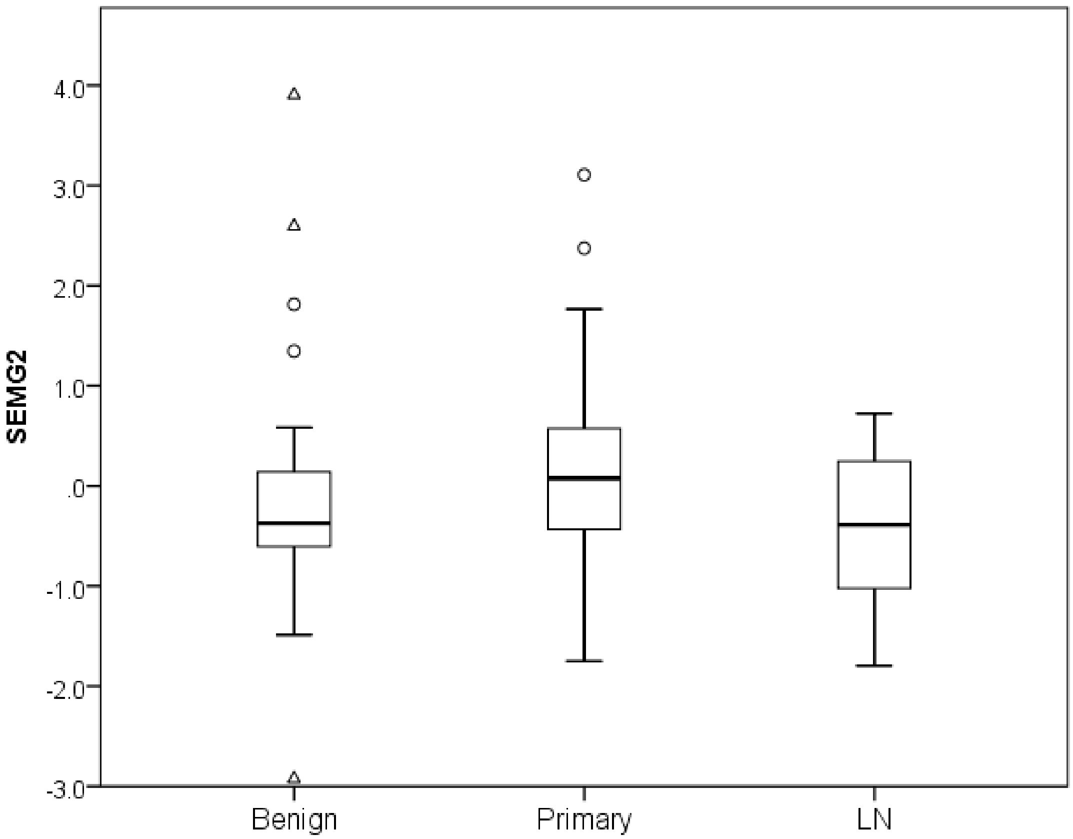
SEMG2
4.0
3.0
2.0
1.0
.0
-1.0
-2.0
-3.0
Benign
Primary
LN

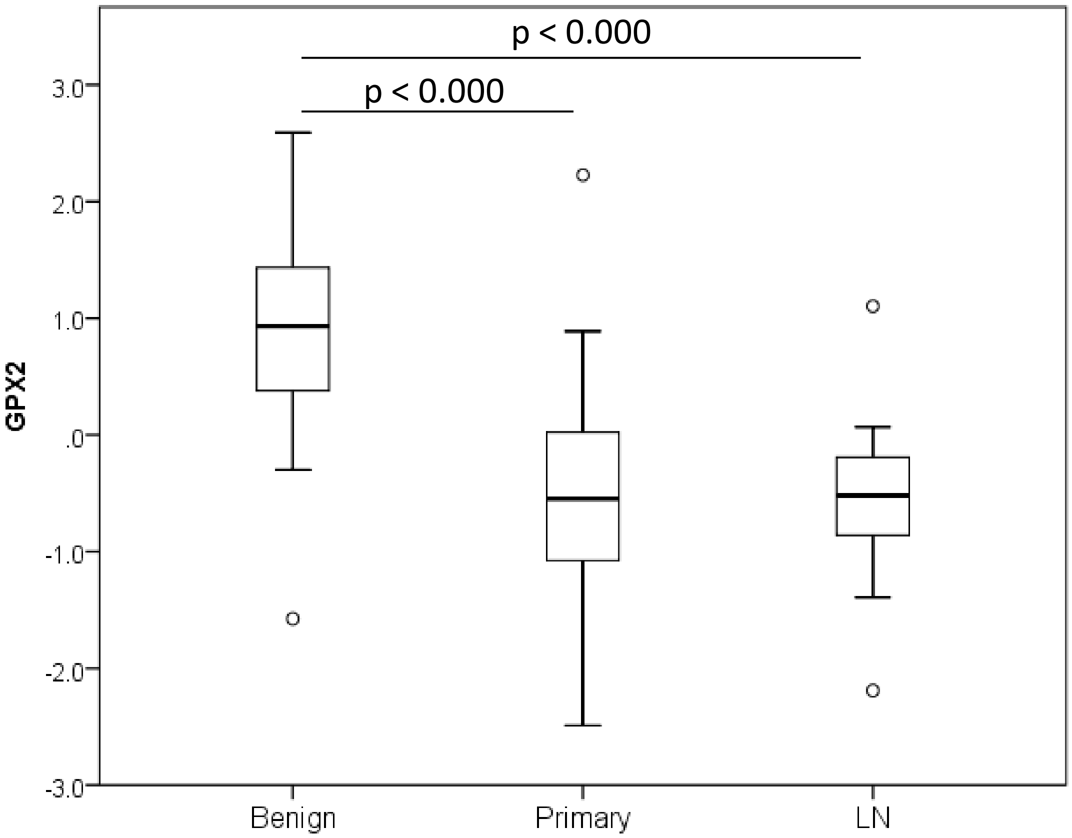
p < 0.000
p < 0.000
GPX2
3.0
2.0
1.0
.0
-1.0
-2.0
-3.0
Benign
Primary
LN

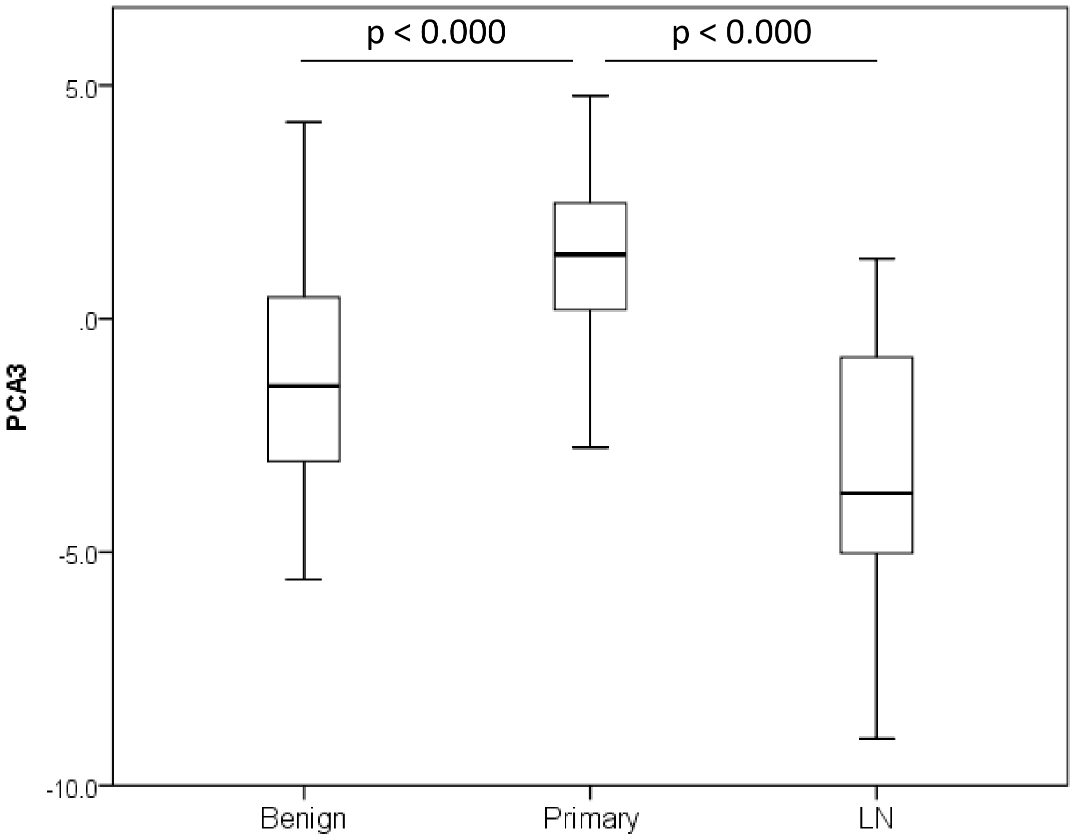

p < 0.000
p < 0.000
5.0
.0
-5.0
-10.0
PCA3
Benign
Primary
LN


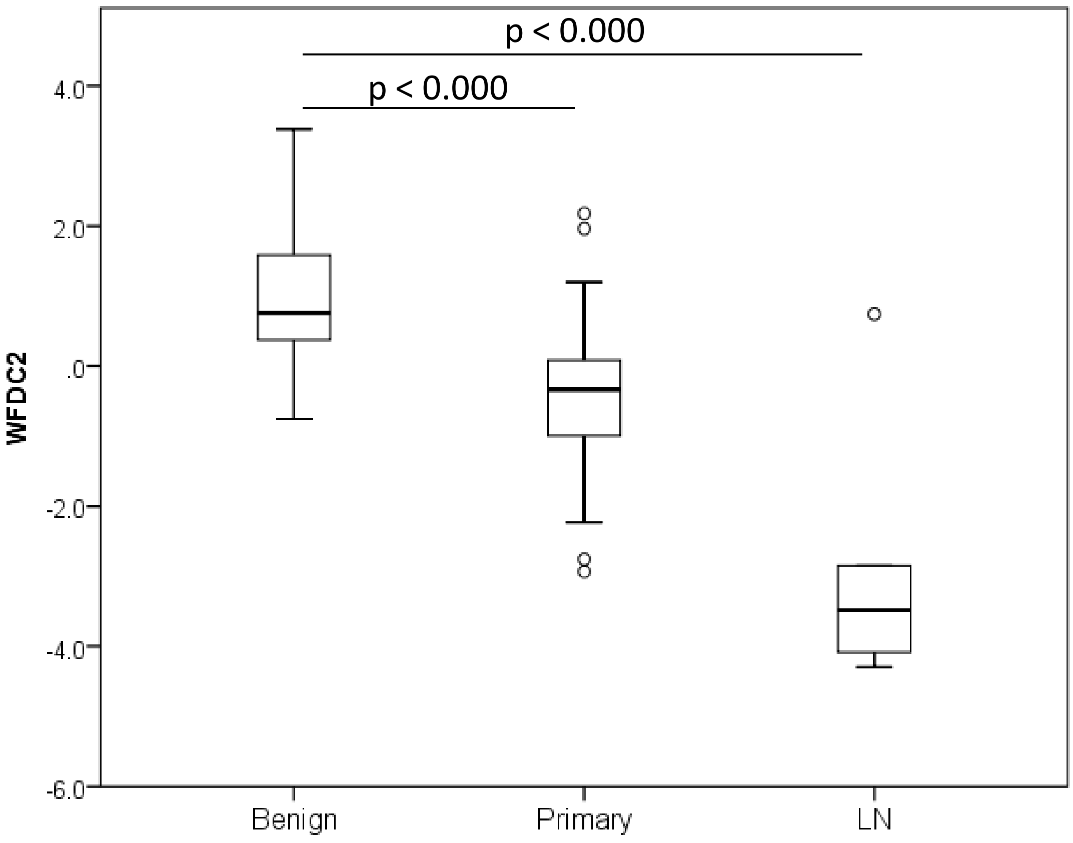

p < 0.000
p < 0.000
4.0
2.0
.0
-2.0
-4.0
-6.0
WFDC2
Benign
Primary
LN

### Lapointe Dataset
### Summary of Results

| P values | Benign vs. Primary | Benign vs. lymph node | Primary vs. Lymph node |
|---|---|---|---|
| OR51E2 | <0.000 | | <0.000 |
| SIM2 | <0.000 | 0.001 | |
| HPN | <0.000 | <0.000 | |
| SLC45A2 | <0.000 | | |
| AMARC | <0.000 | 0.016 | |
| SERPINA5 | 0.001 | 0.001 | |
| HOXC4 | 0.001 | | 0.002 |
| COMP | | | |
| SLC39A2 | 0.020 | | |
| SEMG1 | | | |
| SEMG2 | | | |
| GPX2 | <0.000 | <0.000 | |
| PCA3 | <0.000 | | <0.000 |
| WFDC2 | <0.000 | <0.000 | |

Green: Upregulation (e.g . High in primary vs. Benign)
Red: Downregulation (e.g . Low in primary vs. Benign)

### 2- Ross-Adams dataset

Benign vs. radical prostatectomy vs. TURP cases

Statistics: Kruskal-Wallis test with Bonferroni correction for multiple tests

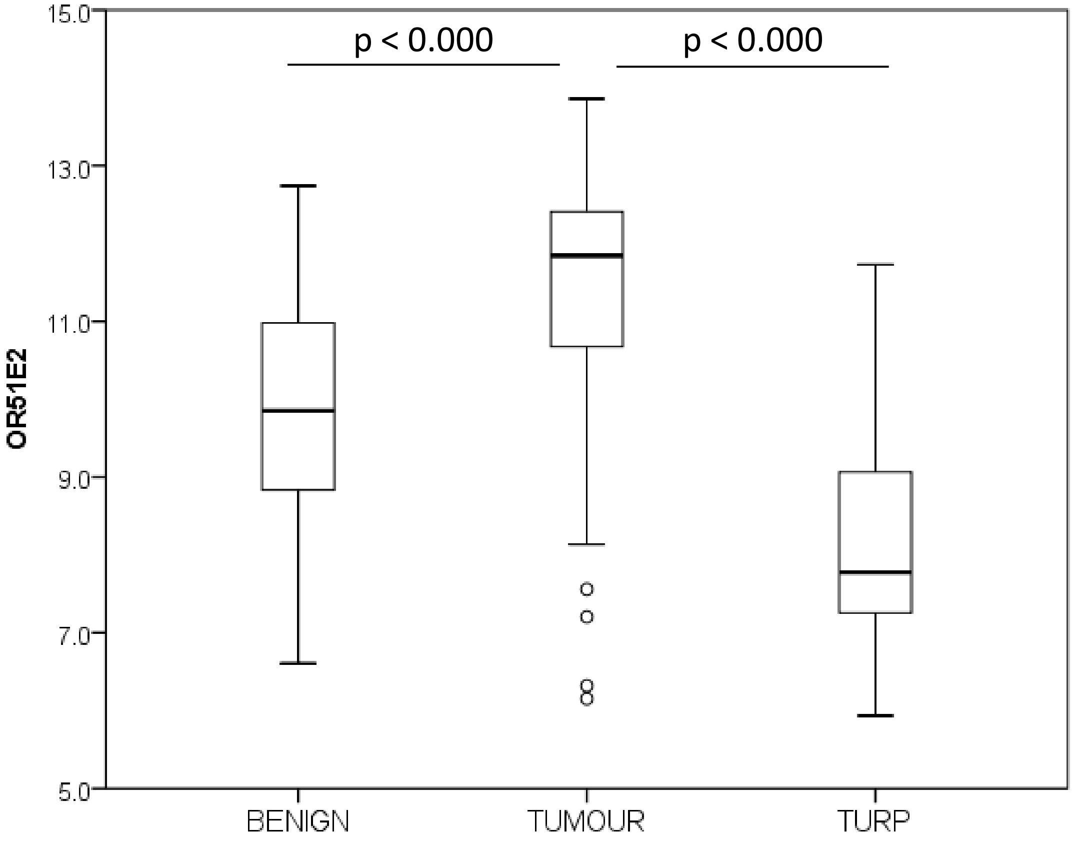
p < 0.000
p < 0.000
OR51E2
15.0
13.0
11.0
9.0
7.0
5.0
BENIGN
TUMOUR
TURP

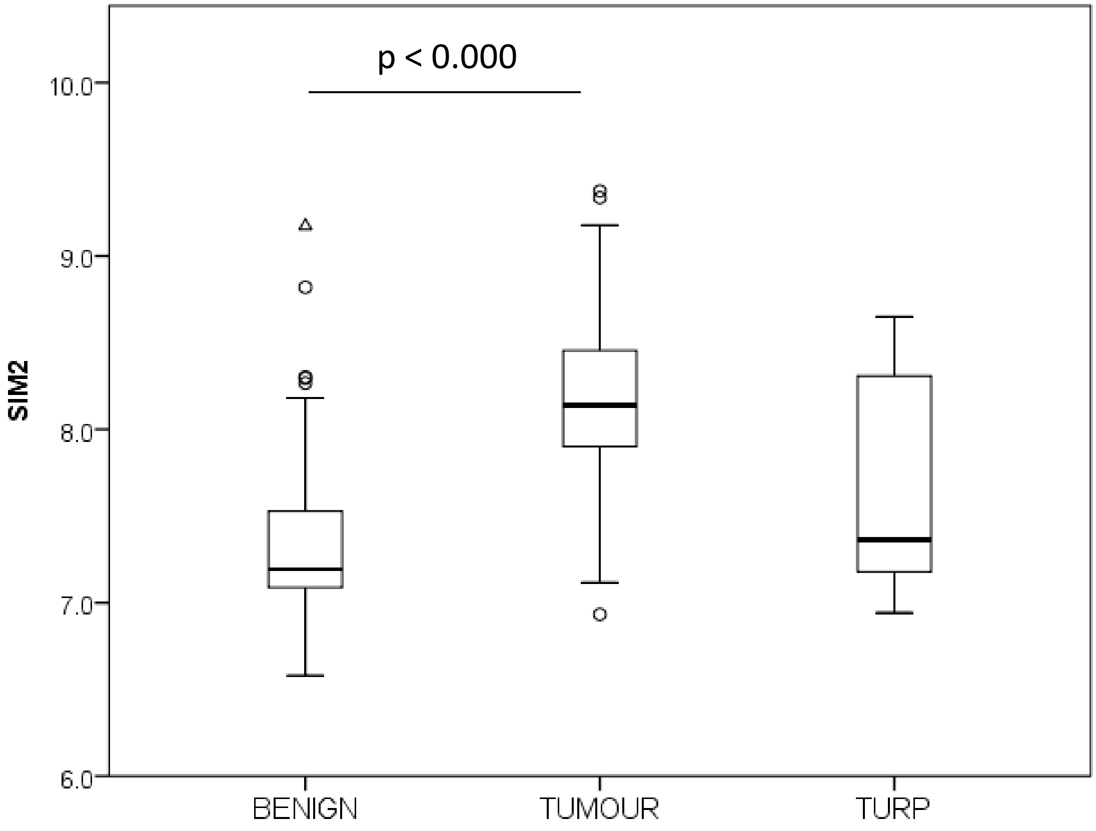
p < 0.000
SIM2
10.0
9.0
8.0
7.0
6.0
BENIGN
TUMOUR
TURP

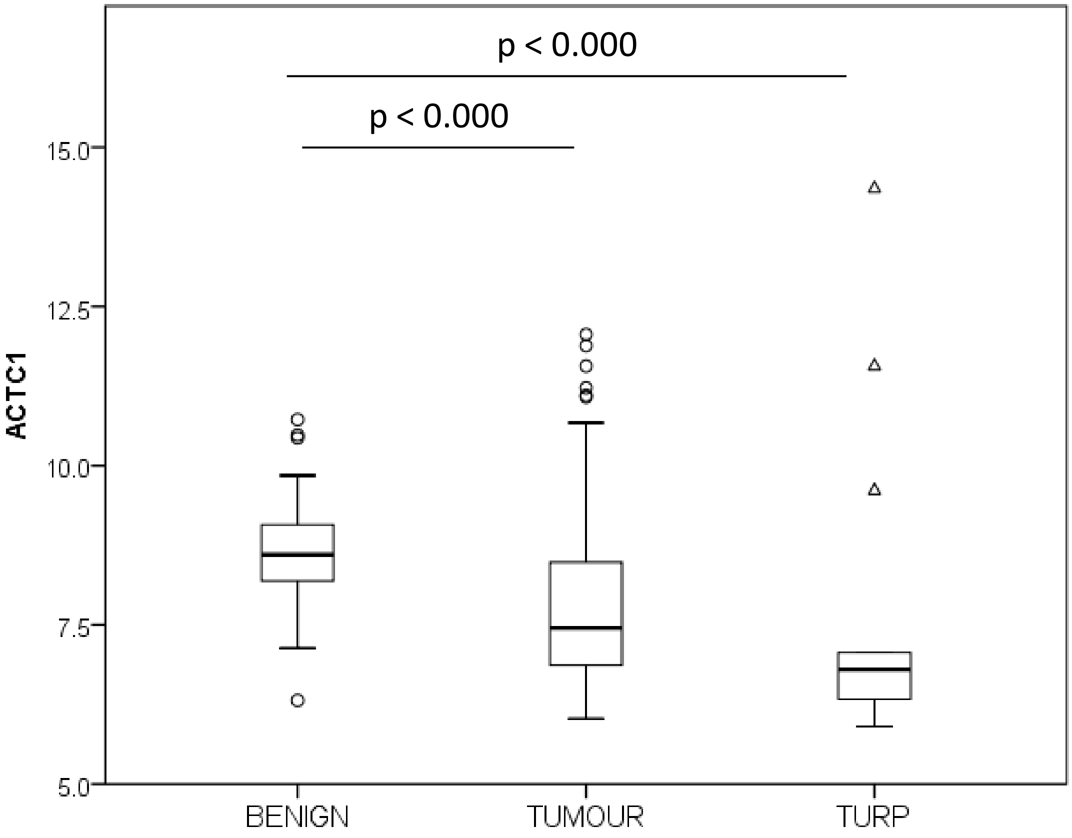

p < 0.000
p < 0.000
ACTC1
15.0
12.5
10.0
7.5
5.0
BENIGN
TUMOUR
TURP


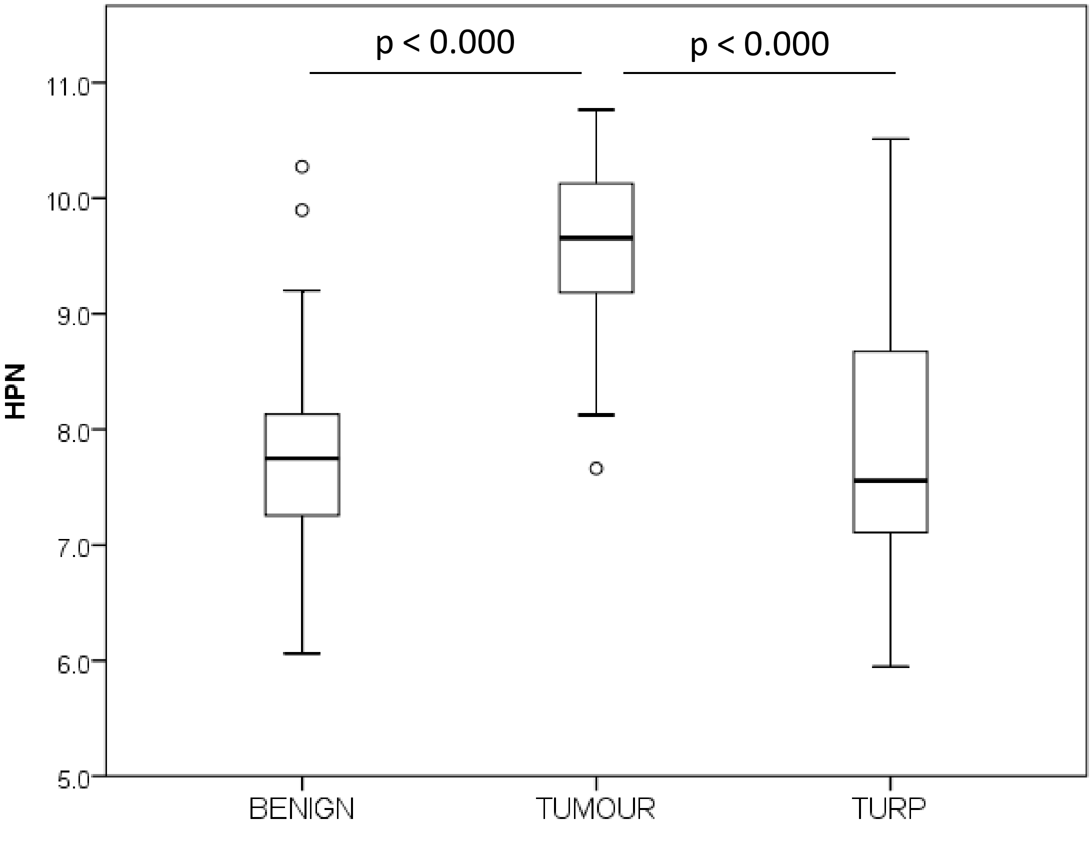

p < 0.000
p < 0.000
HPN
11.0
10.0
9.0
8.0
7.0
6.0
5.0
BENIGN
TUMOUR
TURP

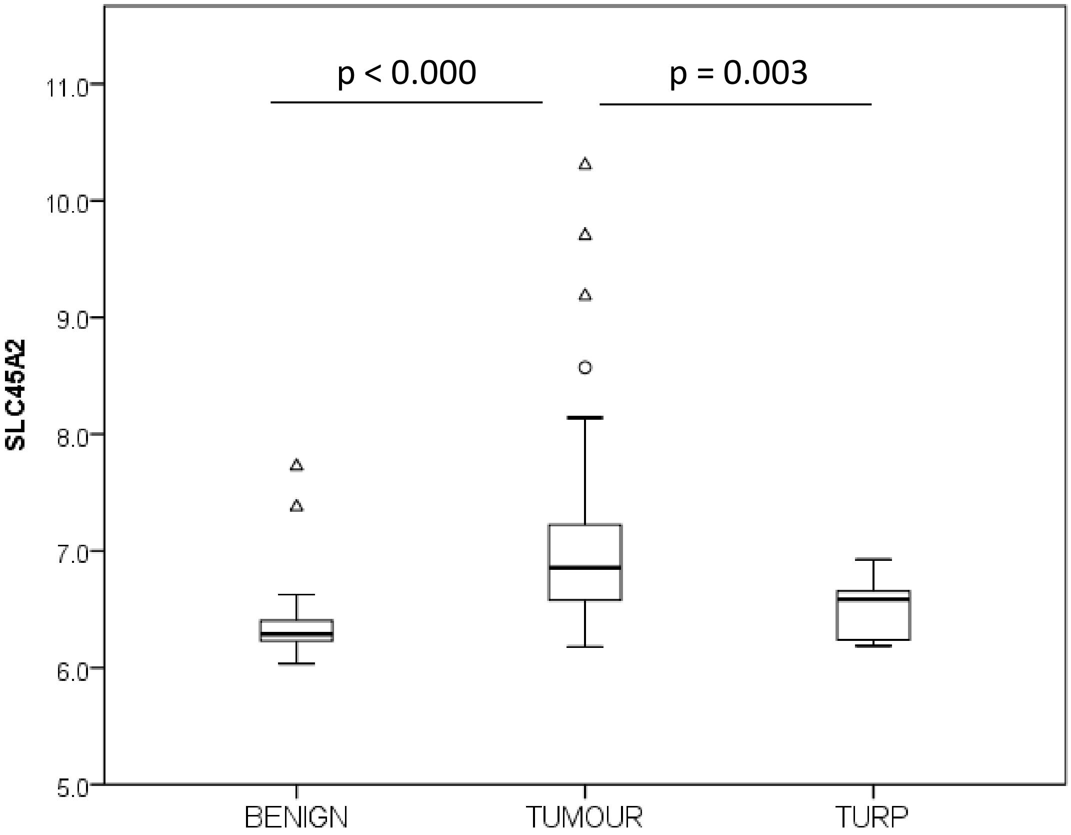
p < 0.000
p = 0.003
SLC45A2
11.0
10.0
9.0
8.0
7.0
6.0
5.0
BENIGN
TUMOUR
TURP

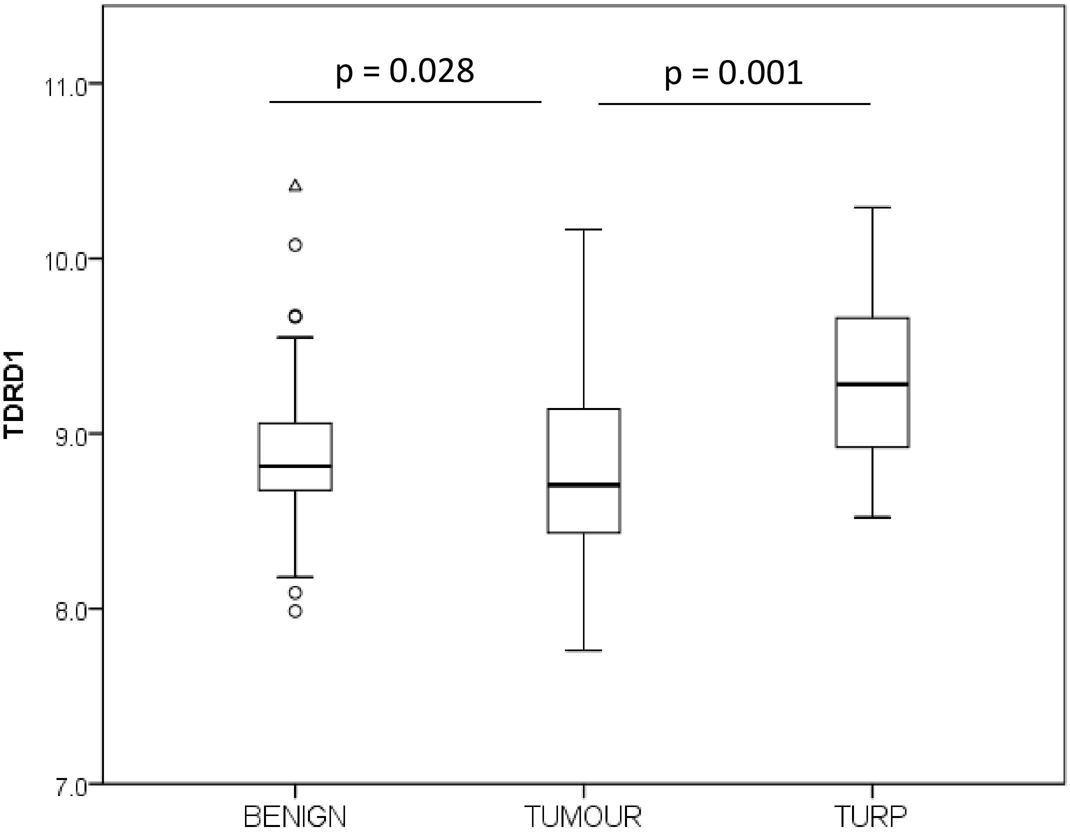
p = 0.028
p = 0.001
TDRD1
11.0
10.0
9.0
8.0
7.0
BENIGN
TUMOUR
TURP

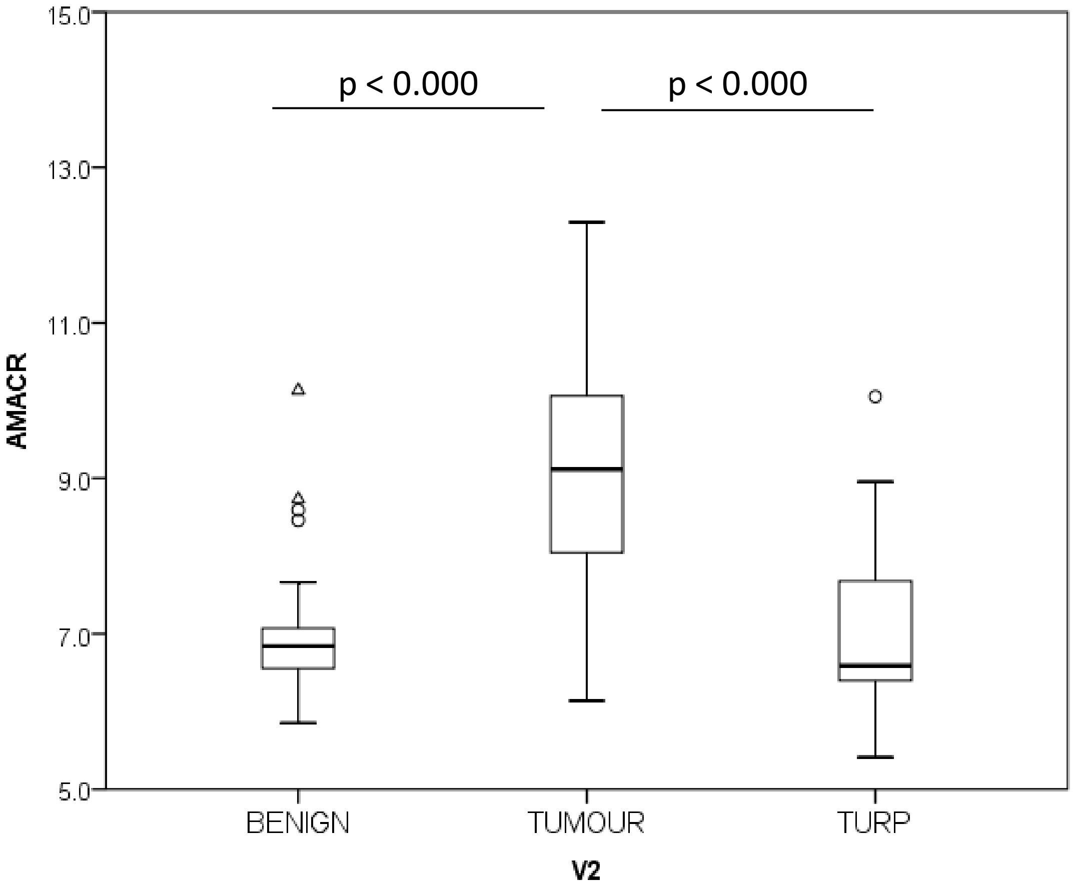

p < 0.000
p < 0.000
AMACR
15.0
13.0
11.0
9.0
7.0
5.0
BENIGN
TUMOUR
TURP
V2


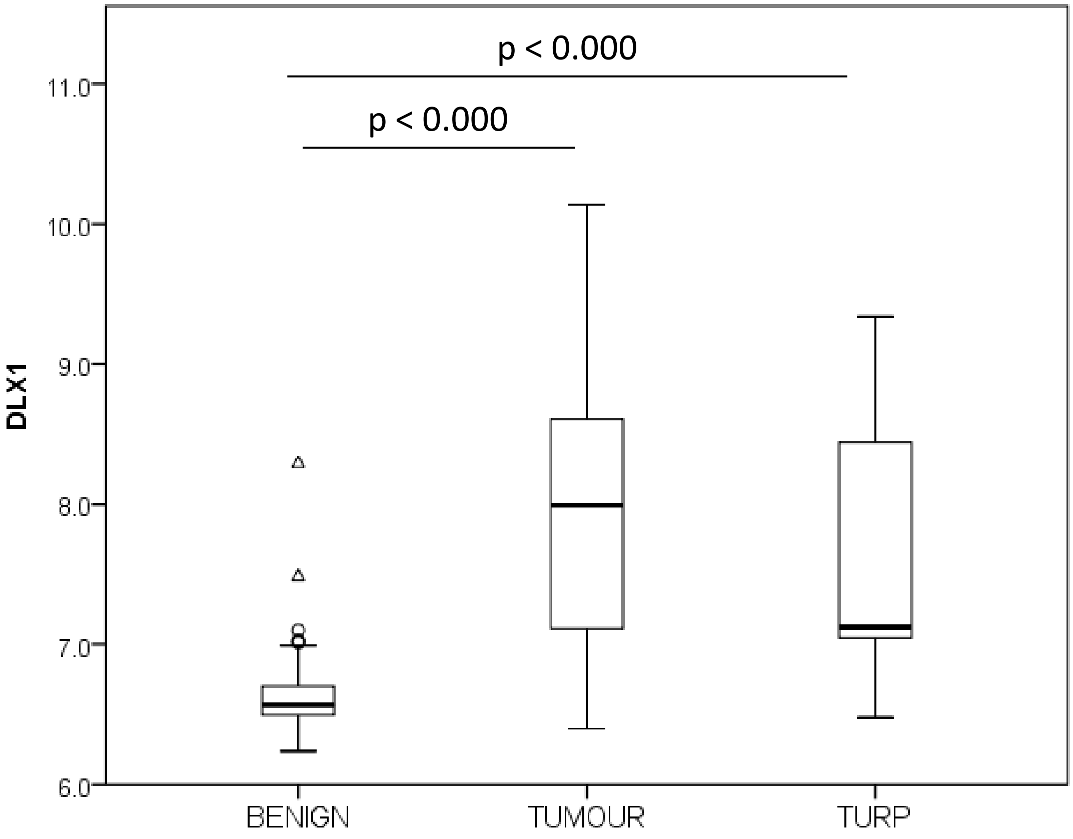

p < 0.000
p < 0.000
DLX1
11.0
10.0
9.0
8.0
7.0
6.0
BENIGN
TUMOUR
TURP

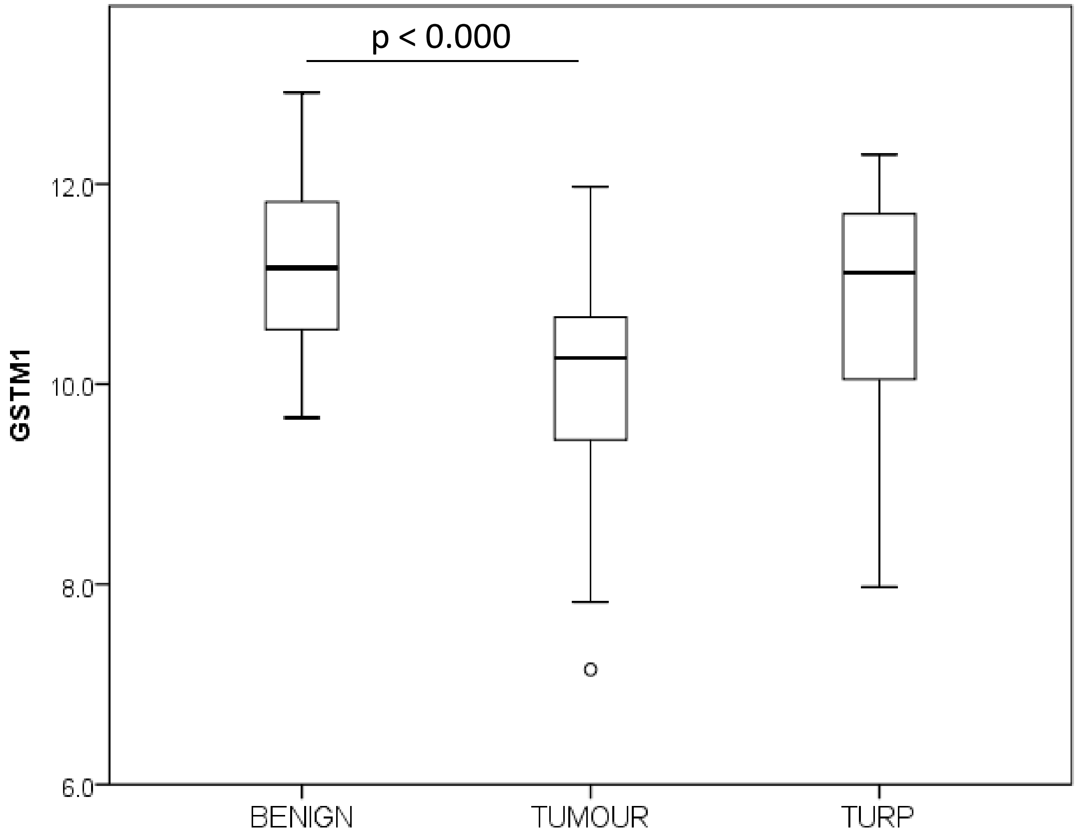
p < 0.000
12.0
10.0
8.0
6.0
GSTM1
BENIGN
TUMOUR
TURP

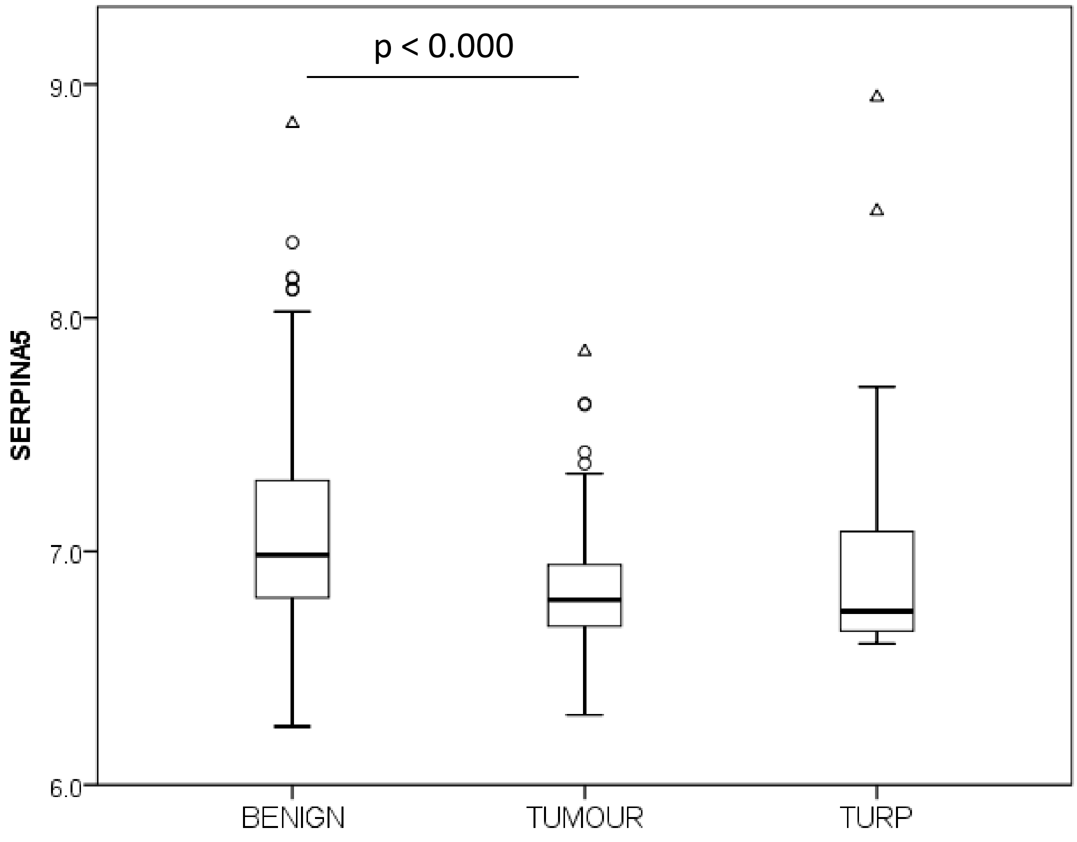
p < 0.000
9.0
8.0
7.0
6.0
SERPINA5
BENIGN
TUMOUR
TURP

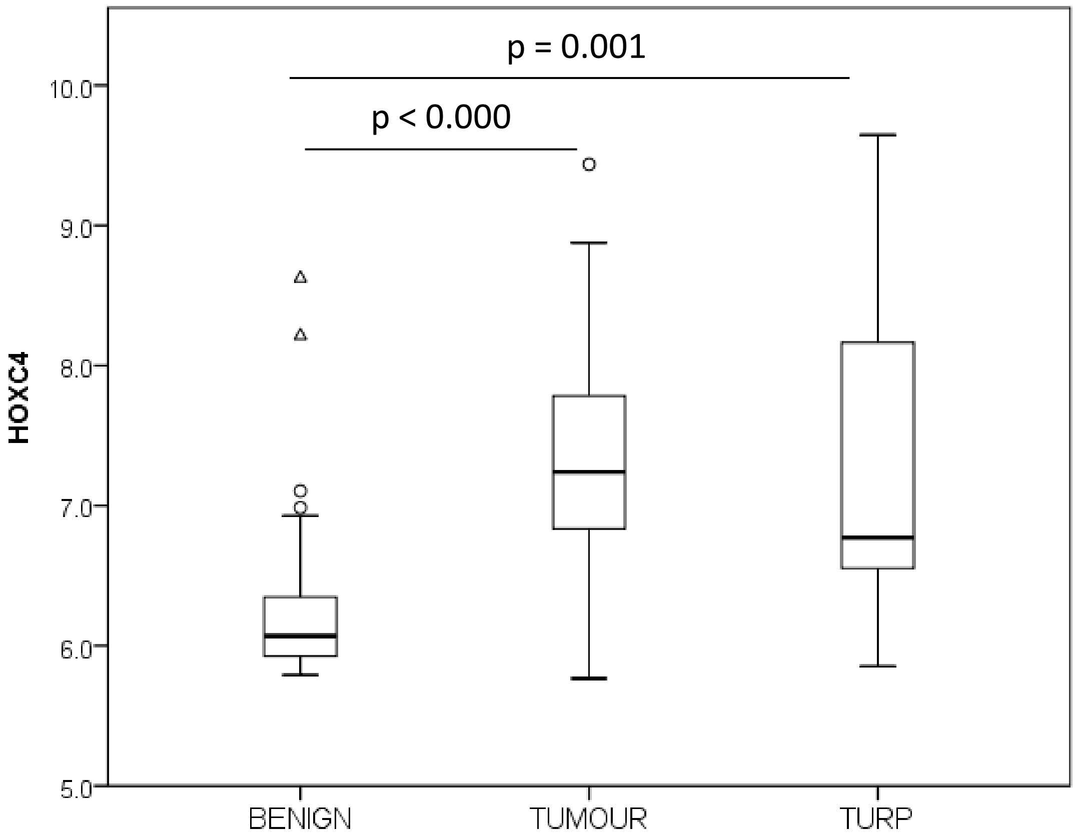
p = 0.001
p < 0.000
HOXC4
10.0
9.0
8.0
7.0
6.0
5.0
BENIGN
TUMOUR
TURP

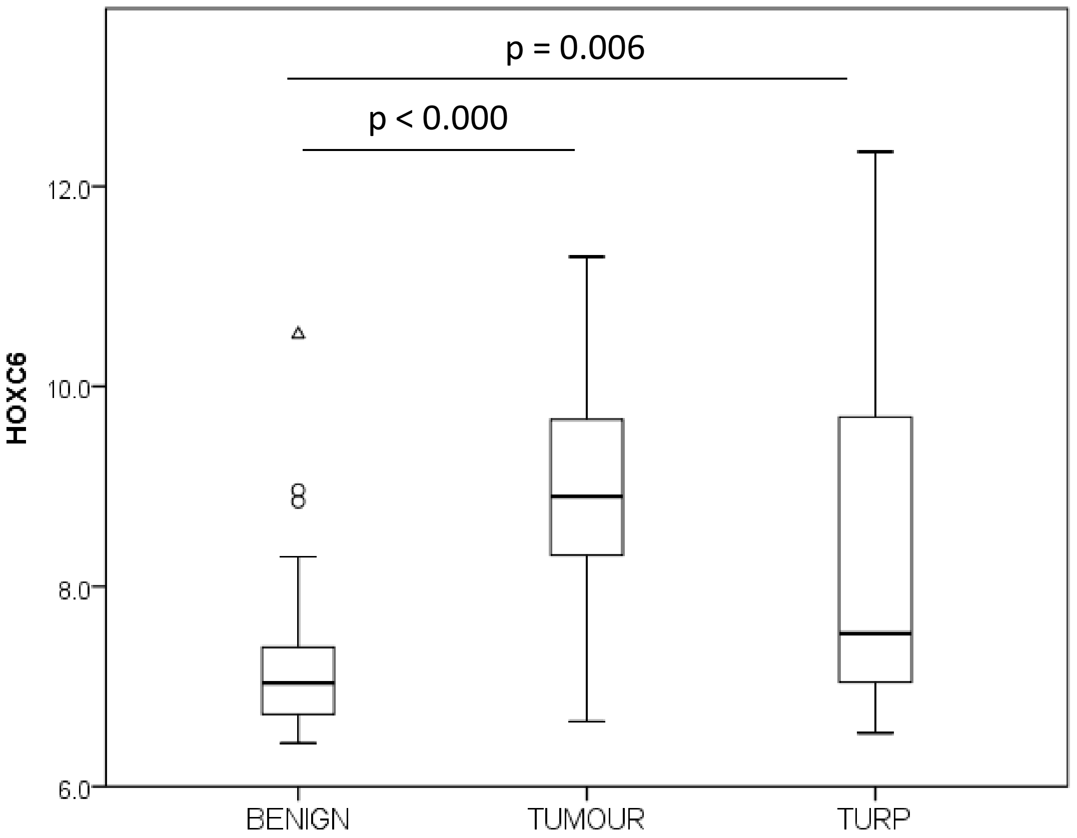
p = 0.006
p < 0.000
HOXC6
12.0
10.0
8.0
6.0
BENIGN
TUMOUR
TURP

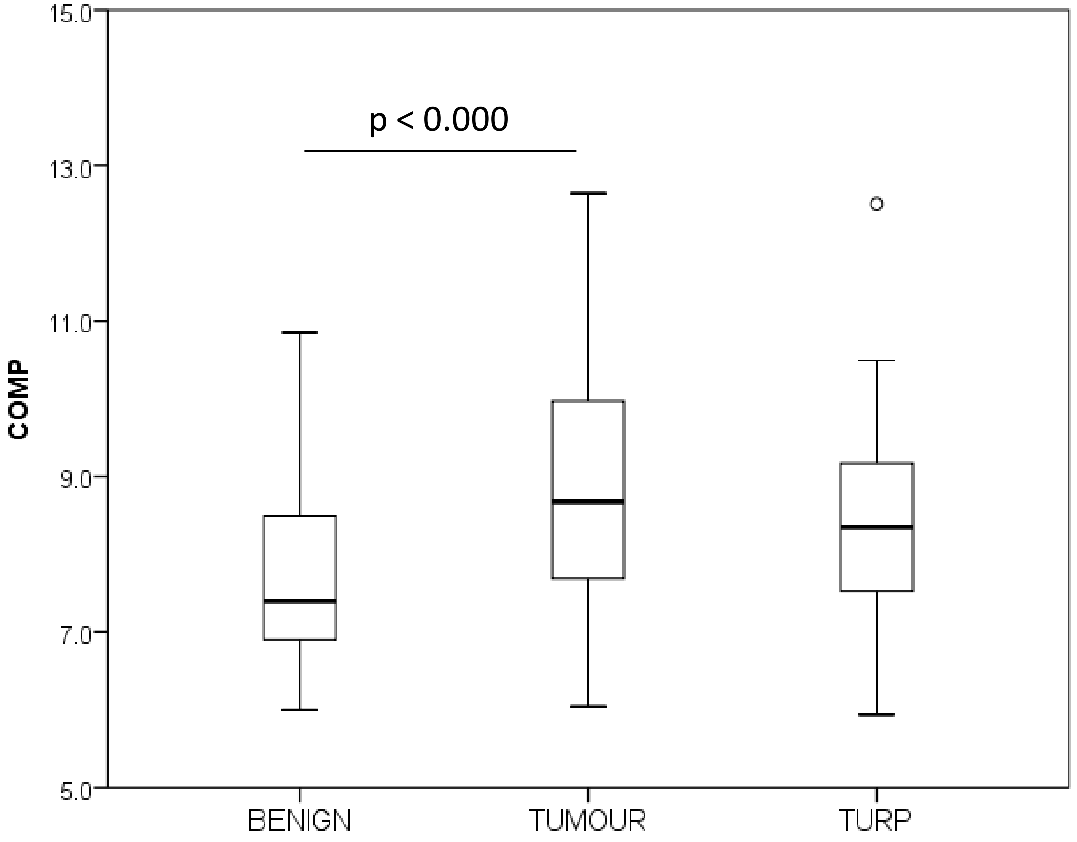

p < 0.000
COMP
15.0
13.0
11.0
9.0
7.0
5.0
BENIGN
TUMOUR
TURP


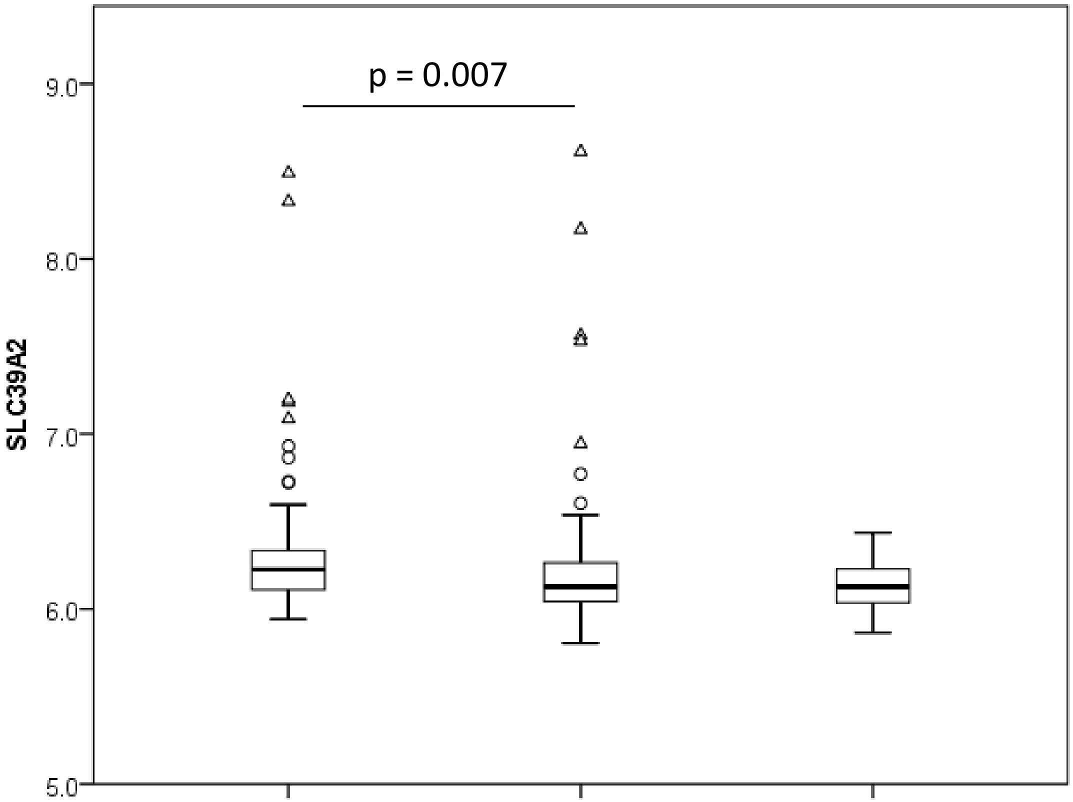

p = 0.007
SLC39A2
9.0
8.0
7.0
6.0
5.0
BENIGN
TUMOUR
TURP

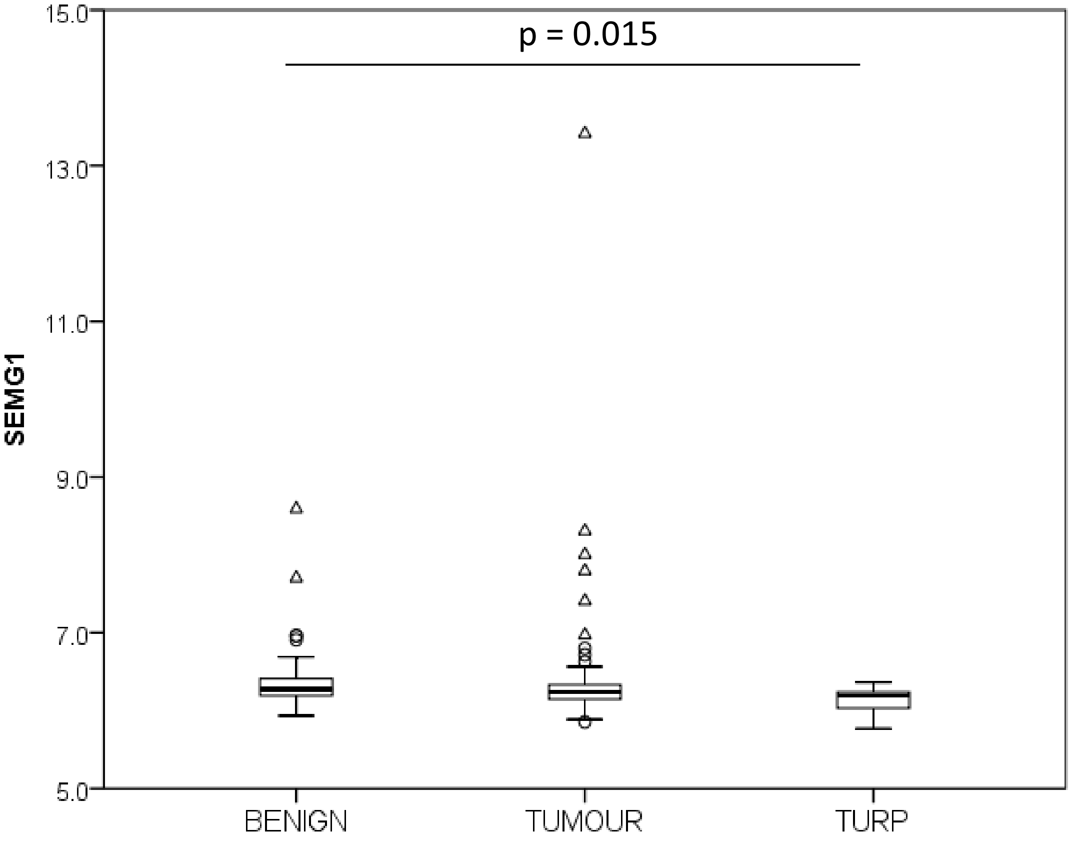

p = 0.015
SEMG1
15.0
13.0
11.0
9.0
7.0
5.0
BENIGN
TUMOUR
TURP


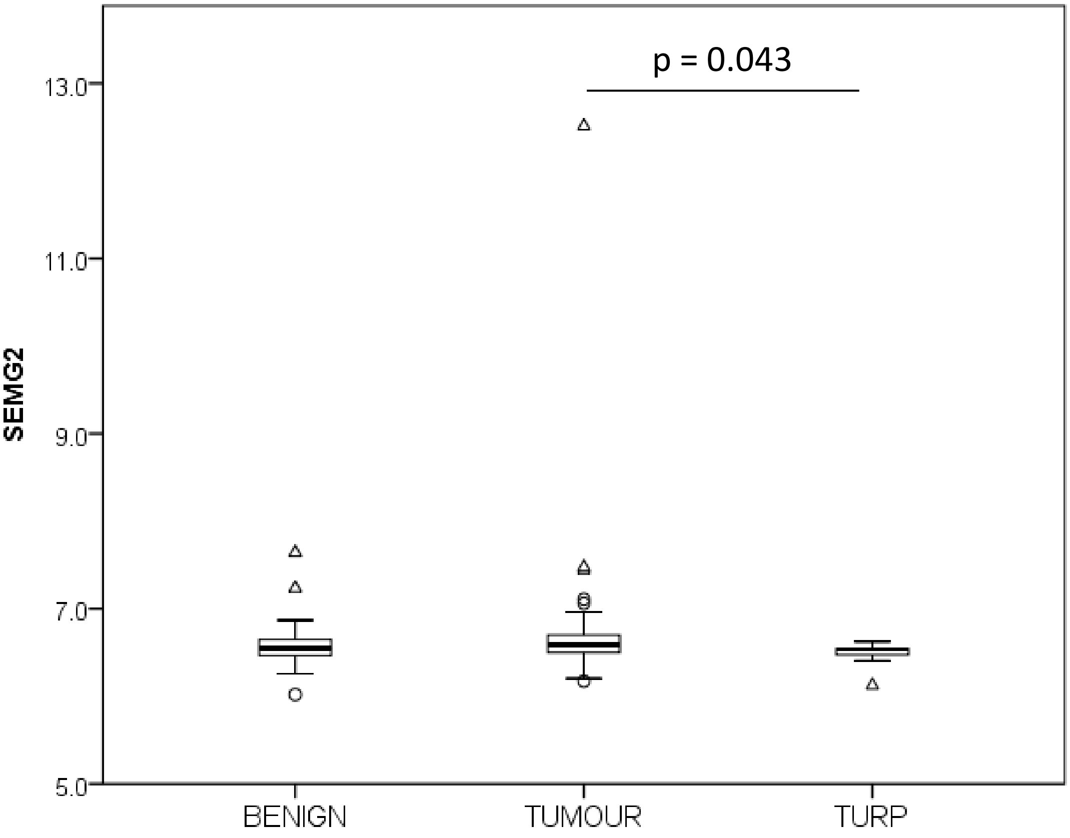

p = 0.043
SEMG2
13.0
11.0
9.0
7.0
5.0
BENIGN
TUMOUR
TURP

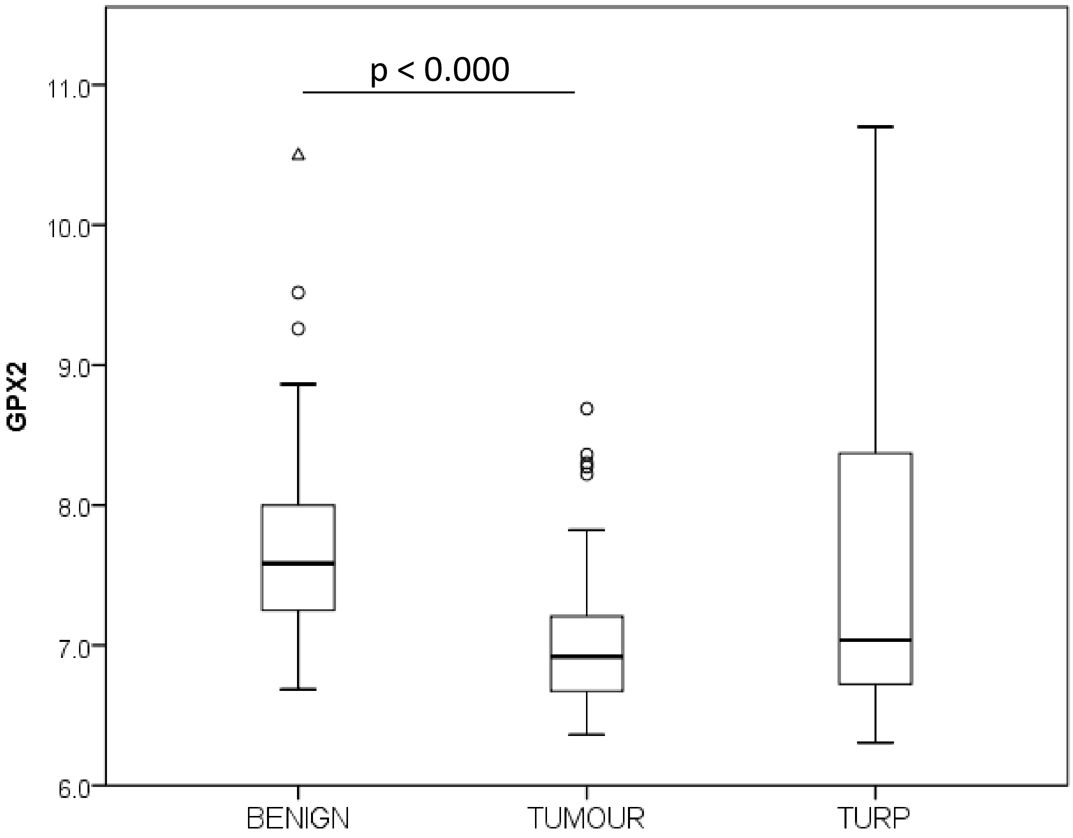
p < 0.000
GPX2
11.0
10.0
9.0
8.0
7.0
6.0
BENIGN
TUMOUR
TURP

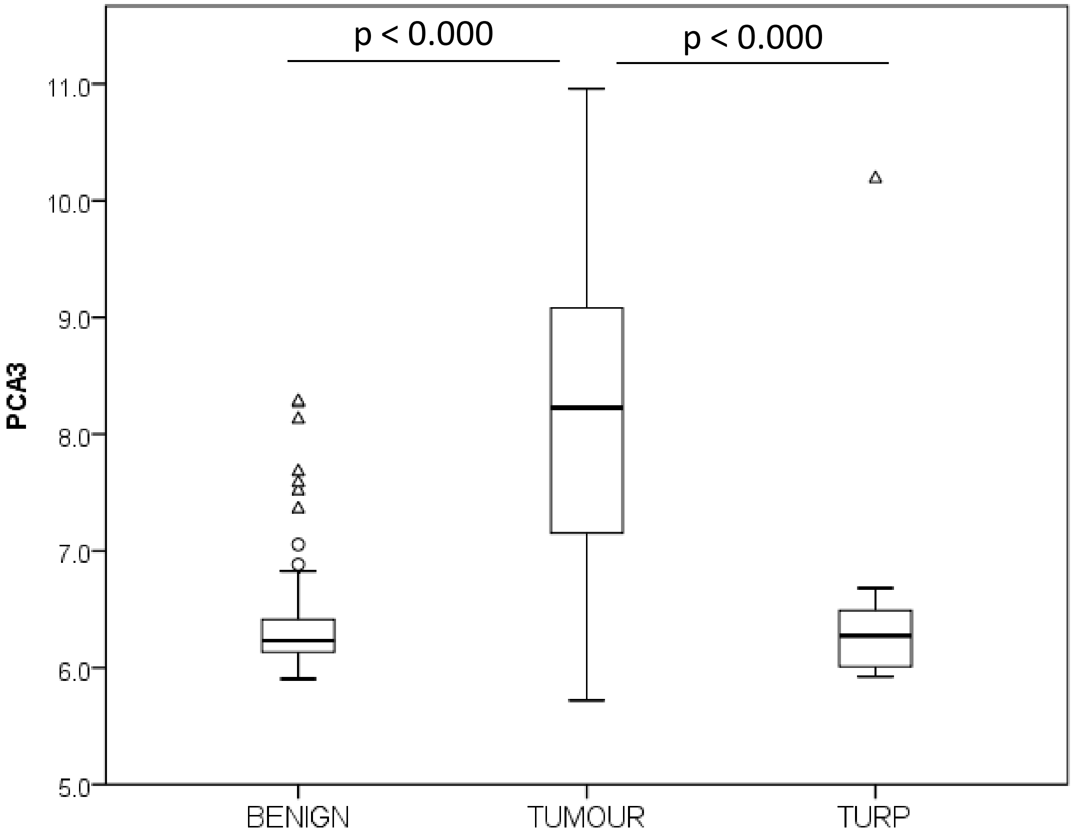
p < 0.000
p < 0.000
PCA3
11.0
10.0
9.0
8.0
7.0
6.0
5.0
BENIGN
TUMOUR
TURP

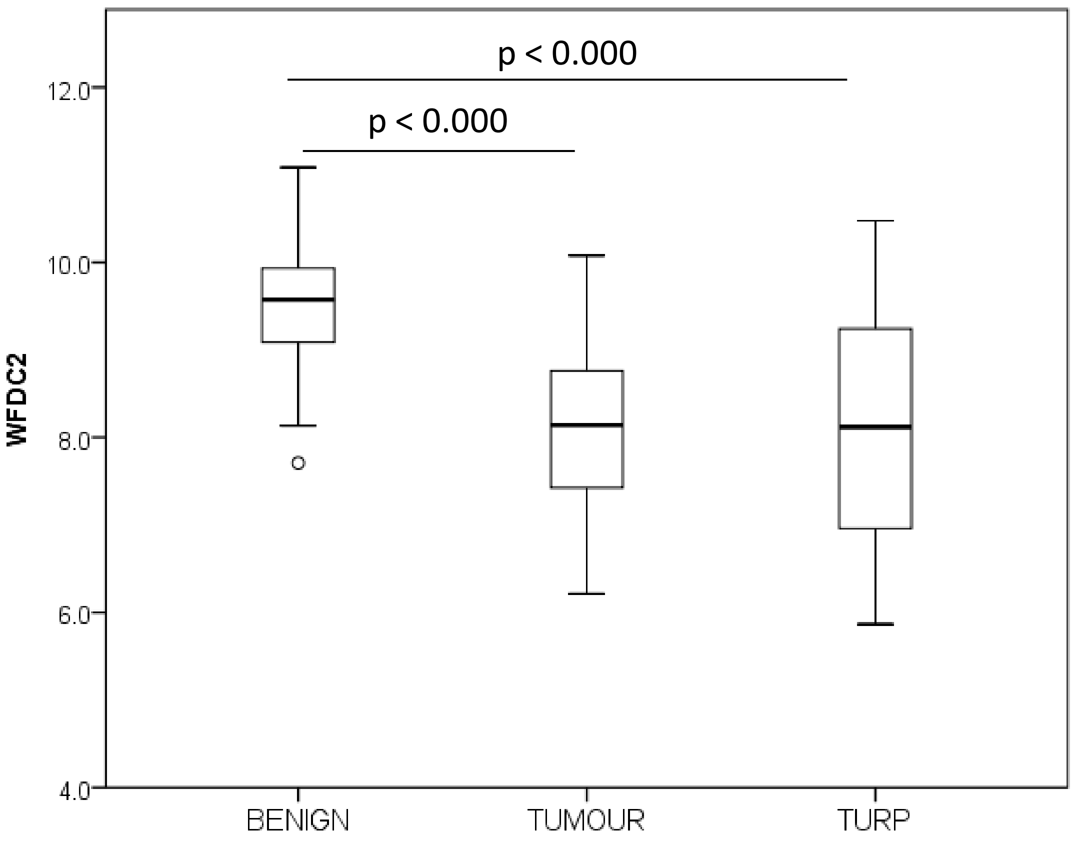

p < 0.000
p < 0.000
12.0
10.0
8.0
6.0
4.0
WFDC2
BENIGN
TUMOUR
TURP

### Ross-Adams Dataset
### Summary of Results

| Gene | Benign vs. RP | Benign vs. TURP | RP vs. TURP |
|---|---|---|---|
| | P | P | P |
| OR51E2 | <0.000 | | <0.000 |
| SIM2 | <0.000 | | |
| ACTC1 | <0.000 | <0.000 | |
| HPN | <0.000 | | <0.000 |
| SLC45A2 | <0.000 | | 0.003 |
| TDRD1 | 0.028 | | 0.001 |
| AMARC | <0.000 | | <0.000 |
| DLX1 | <0.000 | | <0.000 |
| GSTM1 | <0.000 | | |
| SERPINA5 | <0.000 | | |
| HOXC4 | <0.000 | 0.001 | |
| HOXC6 | <0.000 | 0.006 | |
| COMP | <0.000 | | |
| SLC39A2 | 0.007 | | |
| SEMG1 | | 0.015 | |
| SEMG2 | | | 0.043 |
| GPX2 | <0.000 | | |
| PCA3 | <0.000 | | <0.000 |
| WFDC2 | <0.000 | <0.000 | |

Green: Upregulation (e.g . High in primary vs. Benign)
Red: Downregulation (e.g . Low in primary vs. Benign)

### 3- Taylor dataset

Benign vs. primary tumours vs. diverse metastases

Statistics: Kruskal-Wallis test with Bonferroni correction for multiple tests

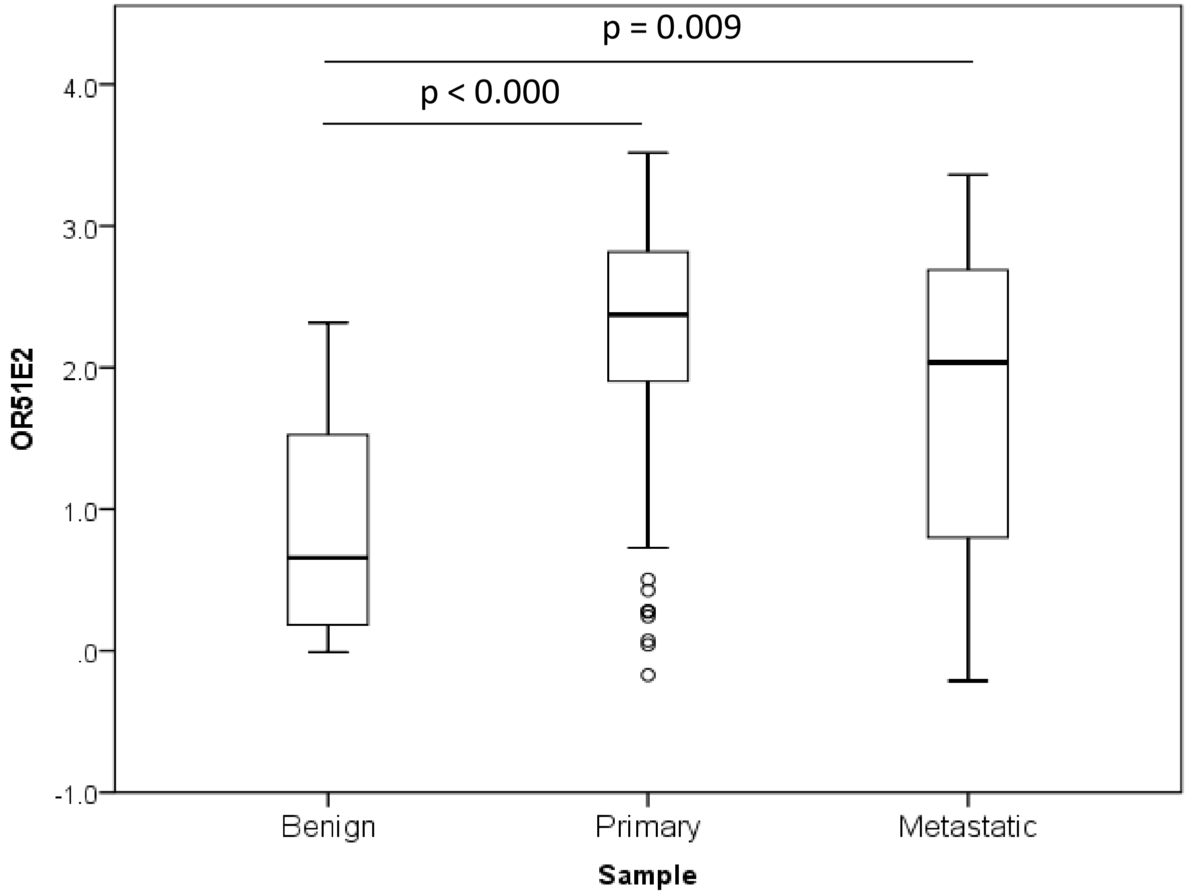
p = 0.009
p < 0.000
4.0
3.0
2.0
1.0
.0
-1.0
OR51E2
Benign
Primary
Metastatic
Sample

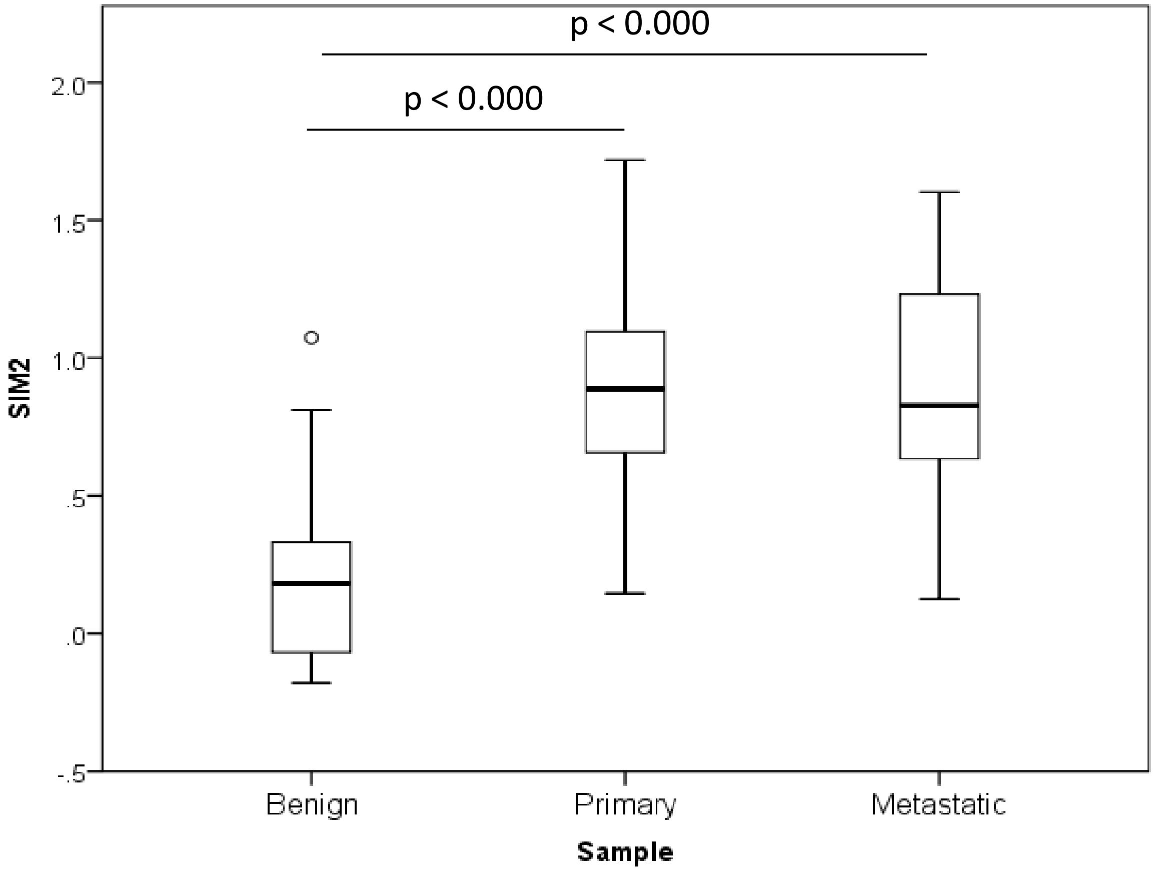
p < 0.000
p < 0.000
2.0
1.5
1.0
.5
.0
-.5
SIM2
Benign
Primary
Metastatic
Sample

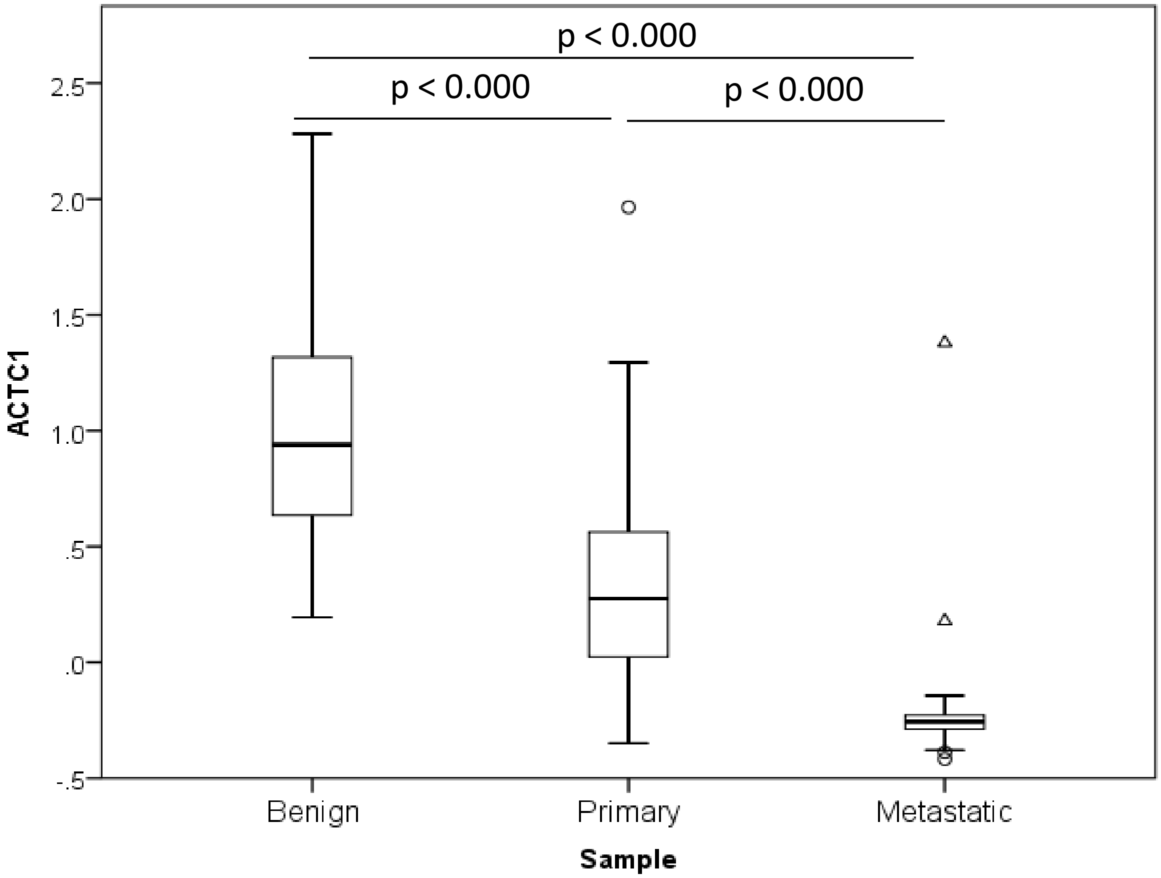
p < 0.000
p < 0.000
p < 0.000
2.5
2.0
1.5
1.0
.5
.0
-.5
ACTC1
Benign
Primary
Metastatic
Sample

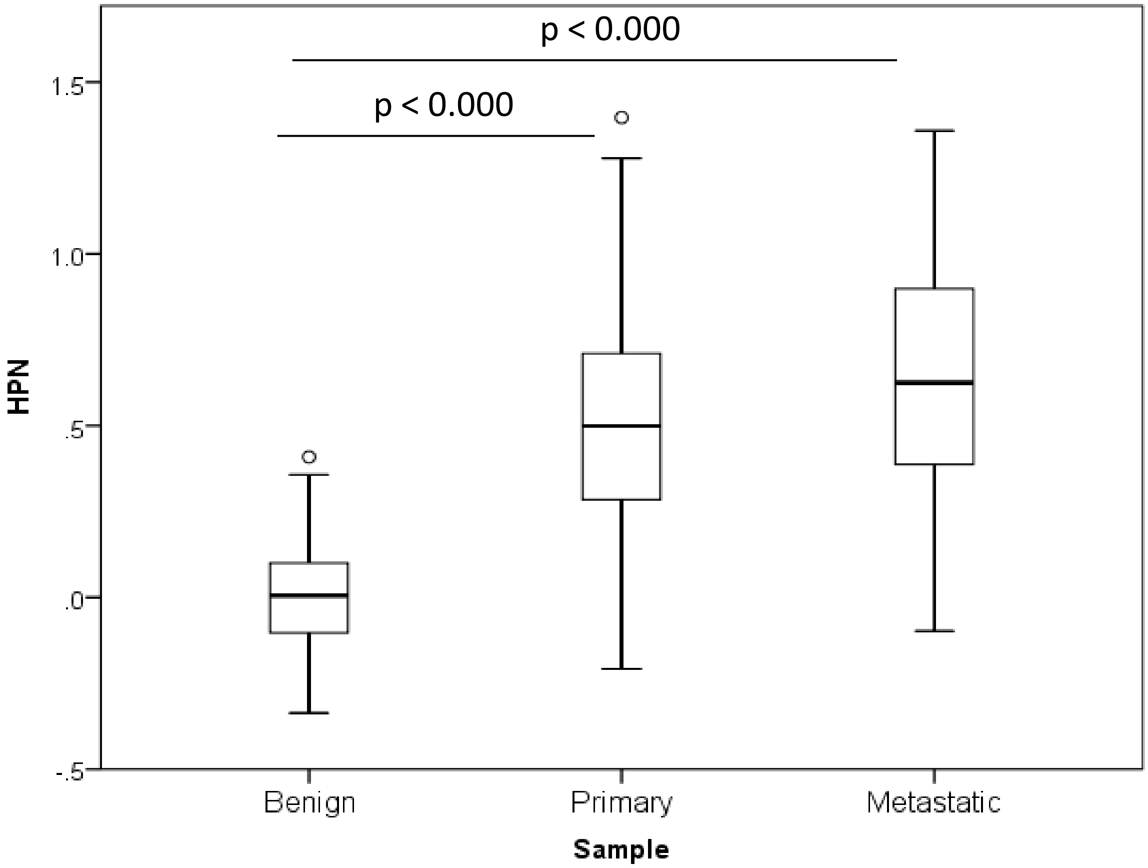
p < 0.000
p < 0.000
1.5
1.0
.5
.0
-.5
HPN
Benign
Primary
Metastatic
Sample

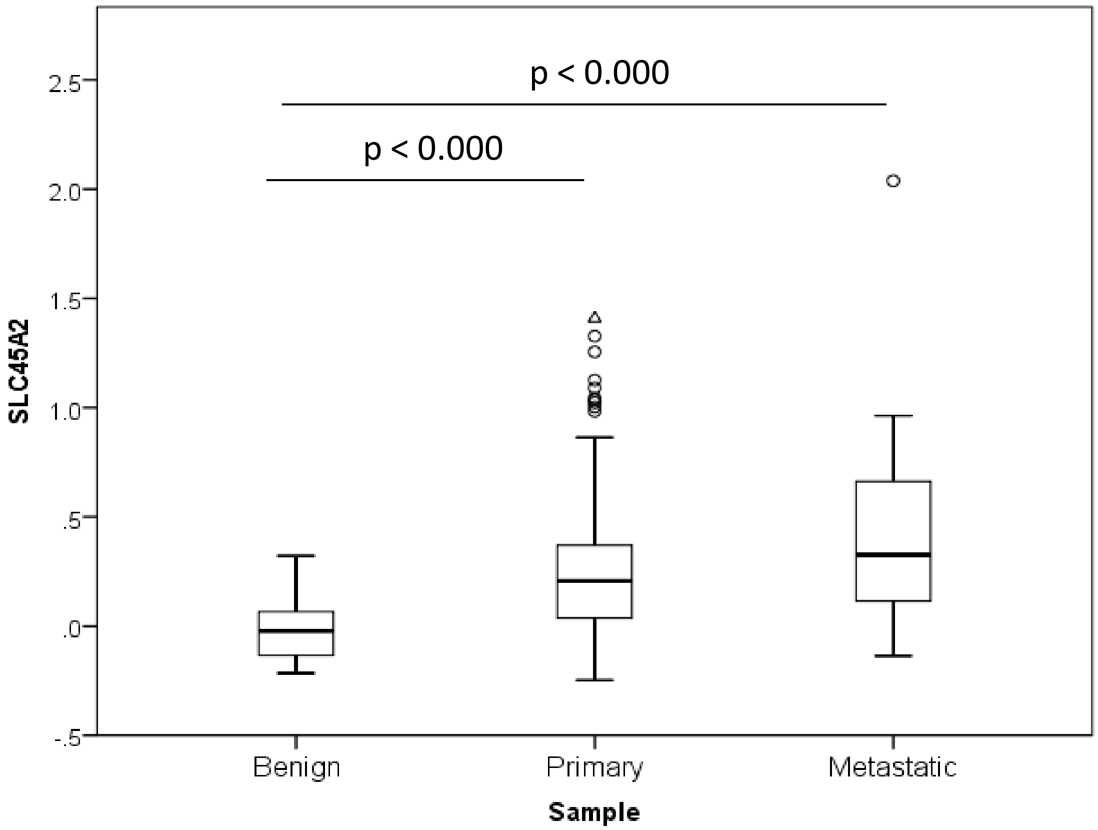
p < 0.000
p < 0.000
SLC45A2
2.5
2.0
1.5
1.0
.5
.0
-.5
Benign
Primary
Metastatic
Sample

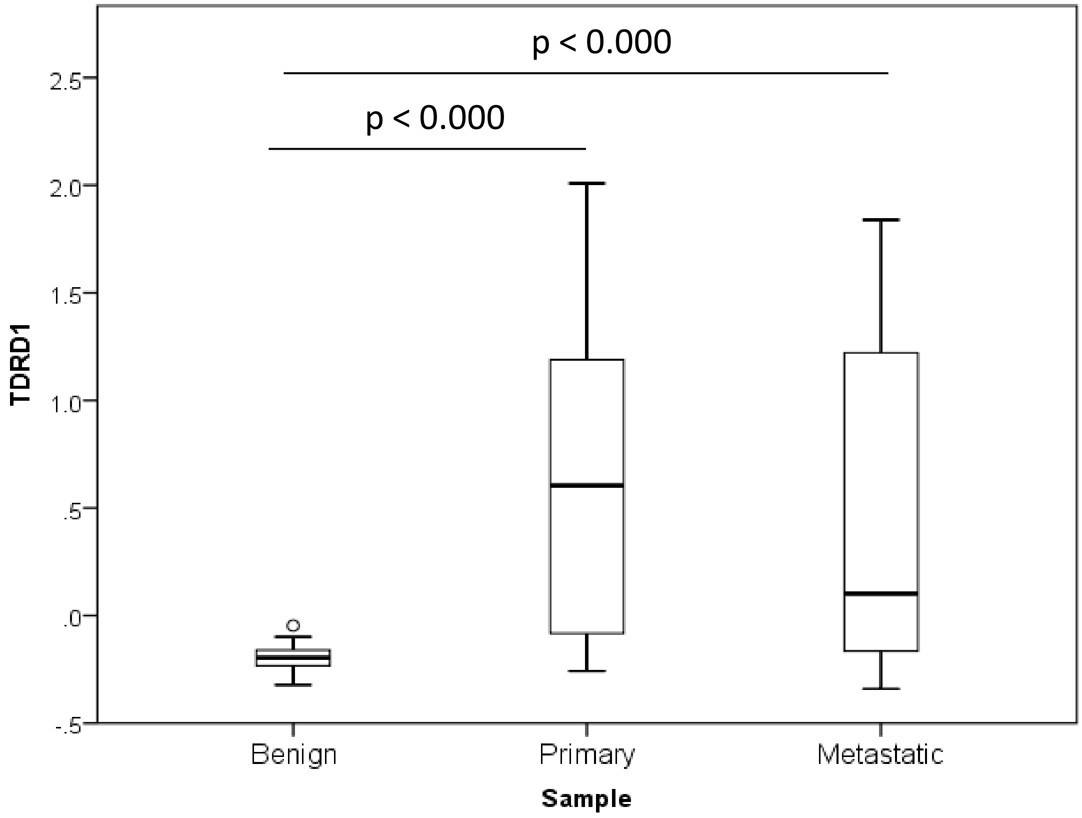
p < 0.000
p < 0.000
TDRD1
2.5
2.0
1.5
1.0
.5
.0
-.5
Benign
Primary
Metastatic
Sample

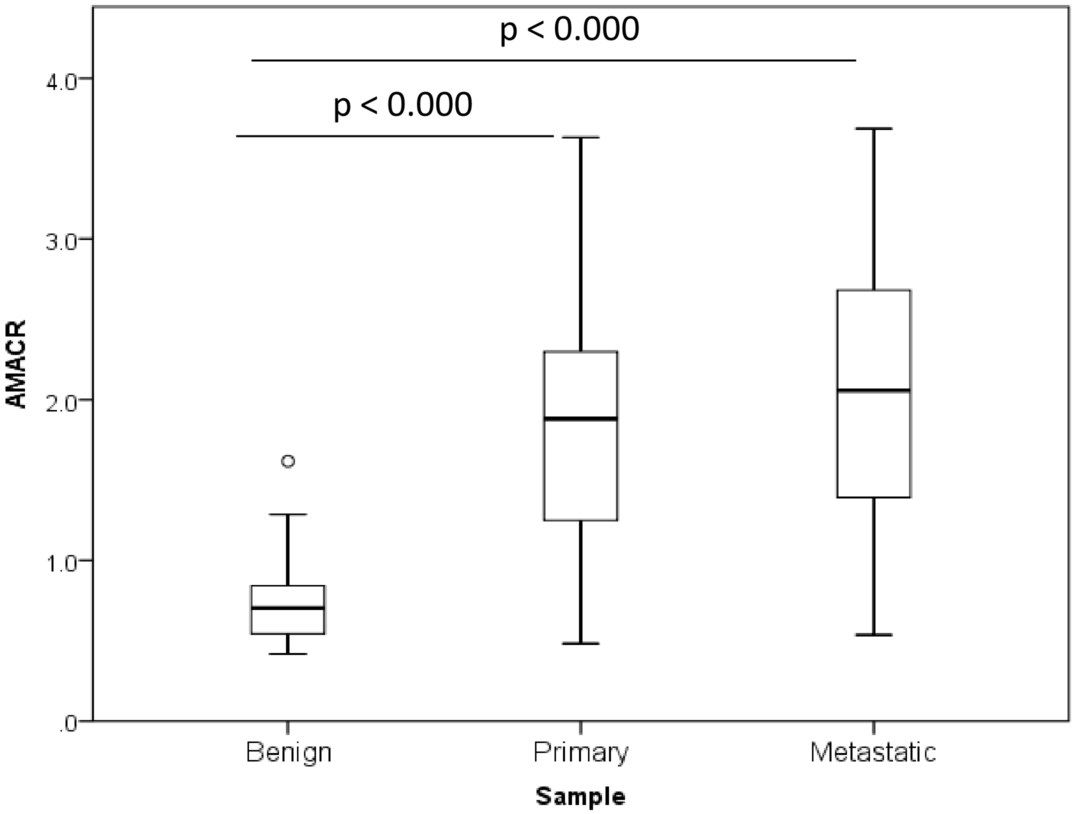
p < 0.000
p < 0.000
4.0
3.0
2.0
1.0
.0
AMACR
Benign
Primary
Metastatic
Sample

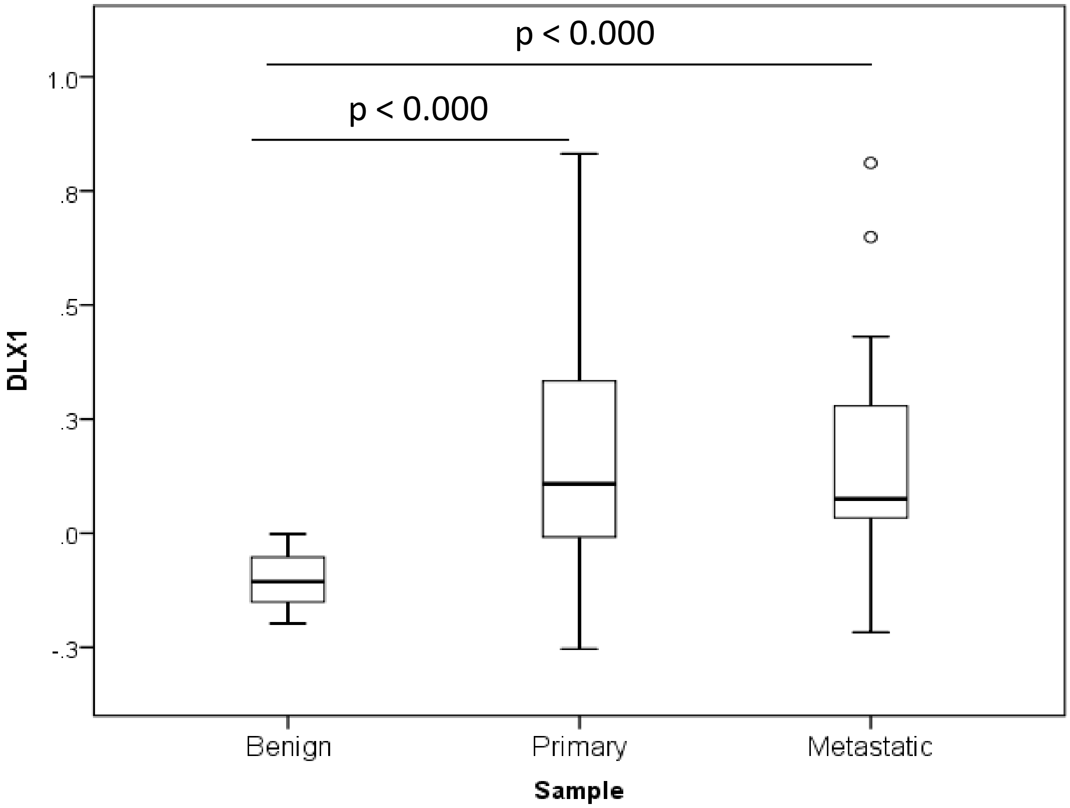
p < 0.000
p < 0.000
1.0
.8
.5
.3
.0
-.3
DLX1
Benign
Primary
Metastatic
Sample

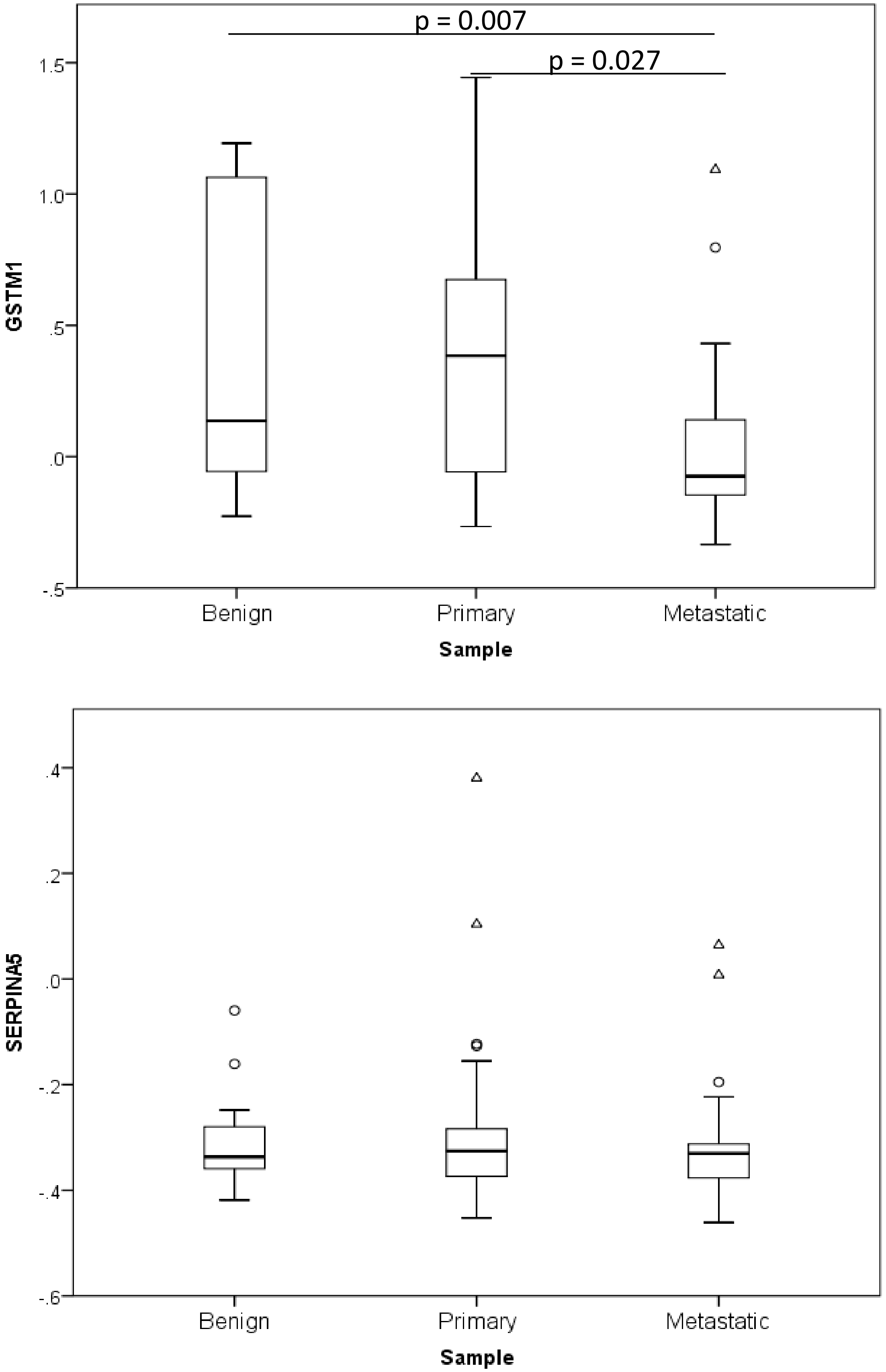
p = 0.007
p = 0.027
1.5
1.0
.5
.0
-.5
GSTM1
Benign
Primary
Metastatic
Sample
.4
.2
.0
-.2
-.4
-.6
SERPINA5
Benign
Primary
Metastatic
Sample

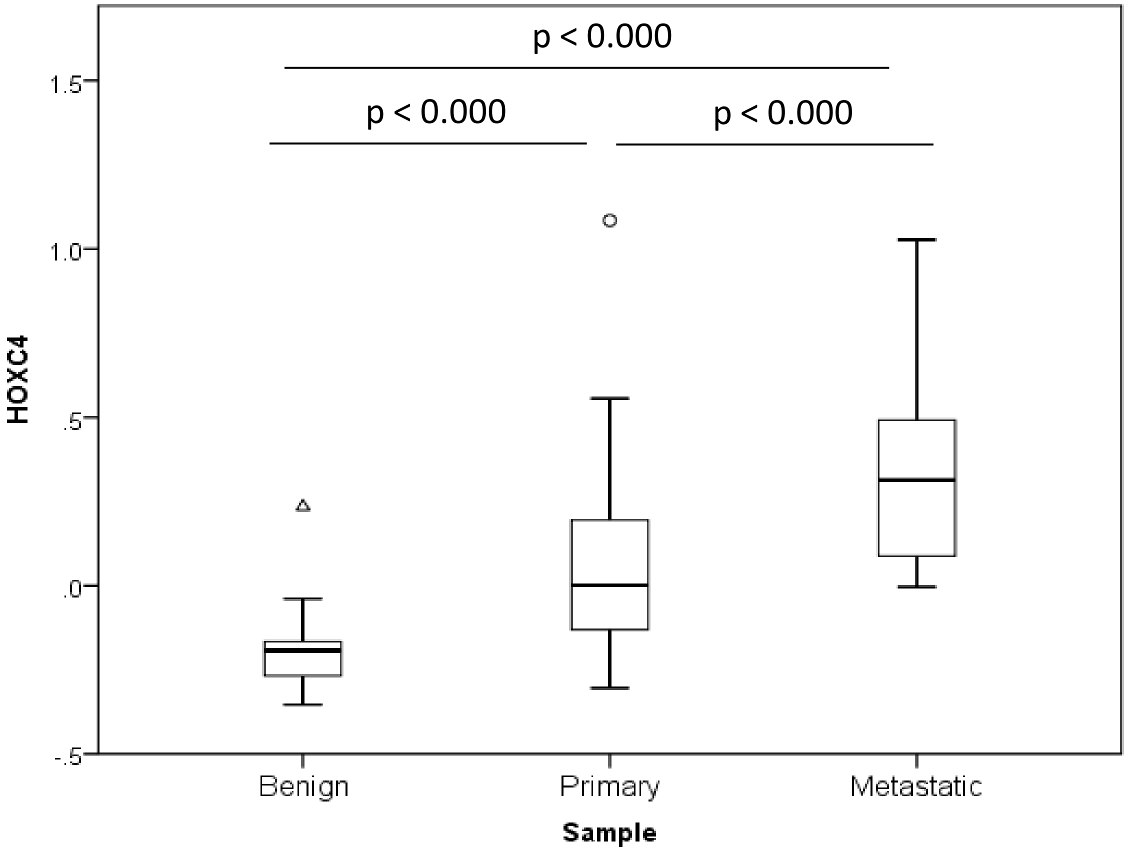
p < 0.000
p < 0.000
p < 0.000
1.5
1.0
.5
.0
-.5
HOXC4
Benign
Primary
Metastatic
Sample

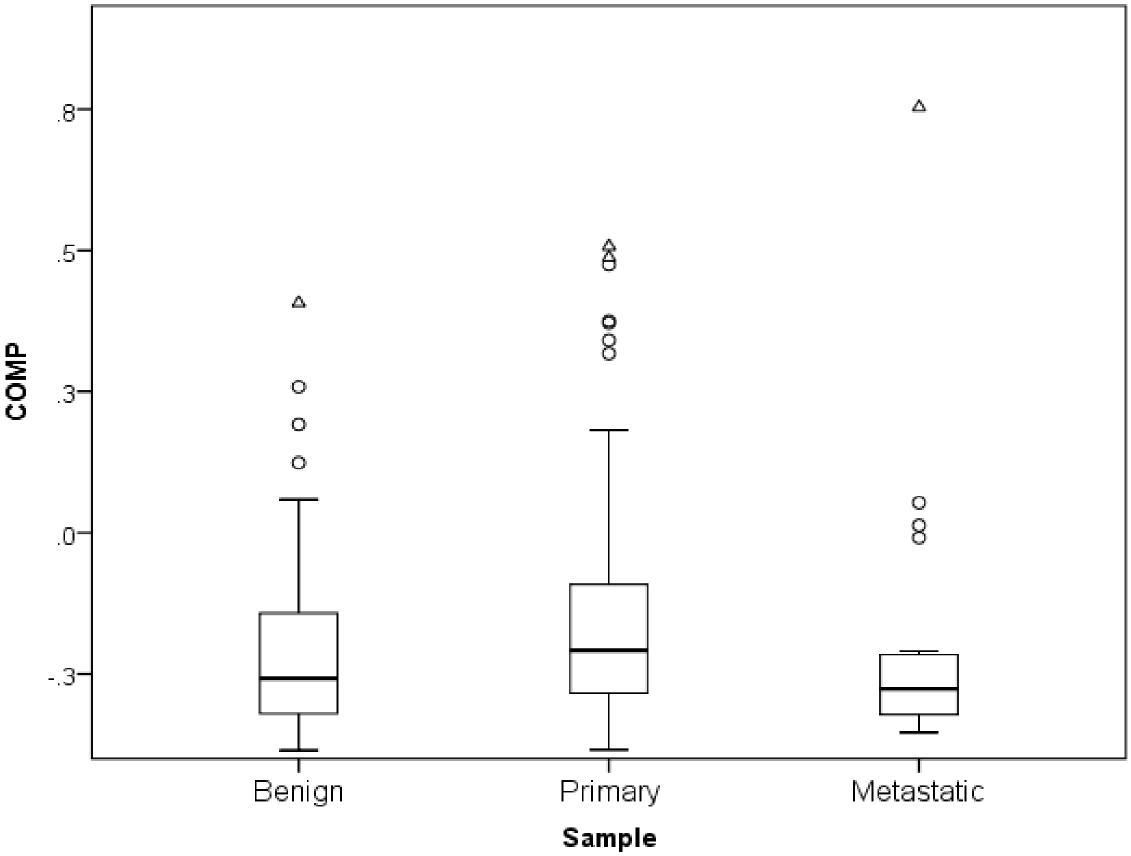
.8
.5
.3
.0
-.3
COMP
Benign
Primary
Metastatic
Sample

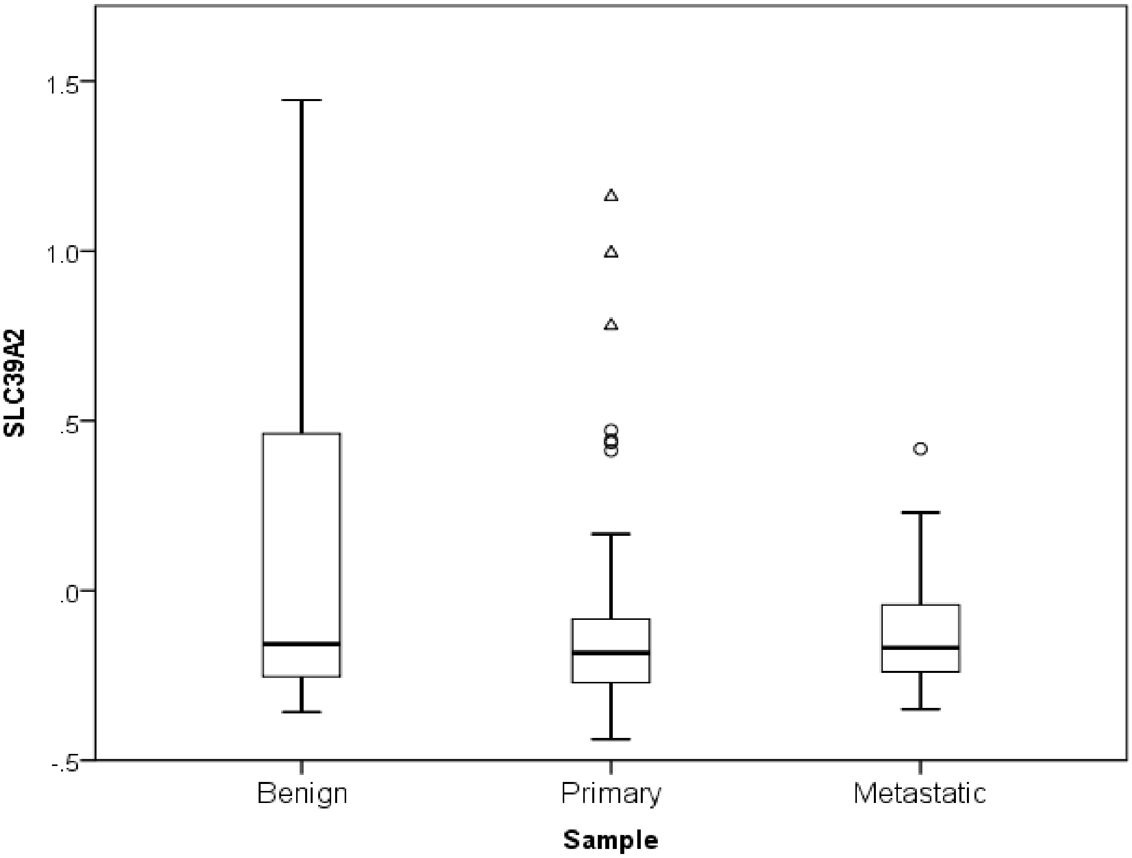
1.5
1.0
.5
.0
-.5
SLC39A2
Benign
Primary
Metastatic
Sample

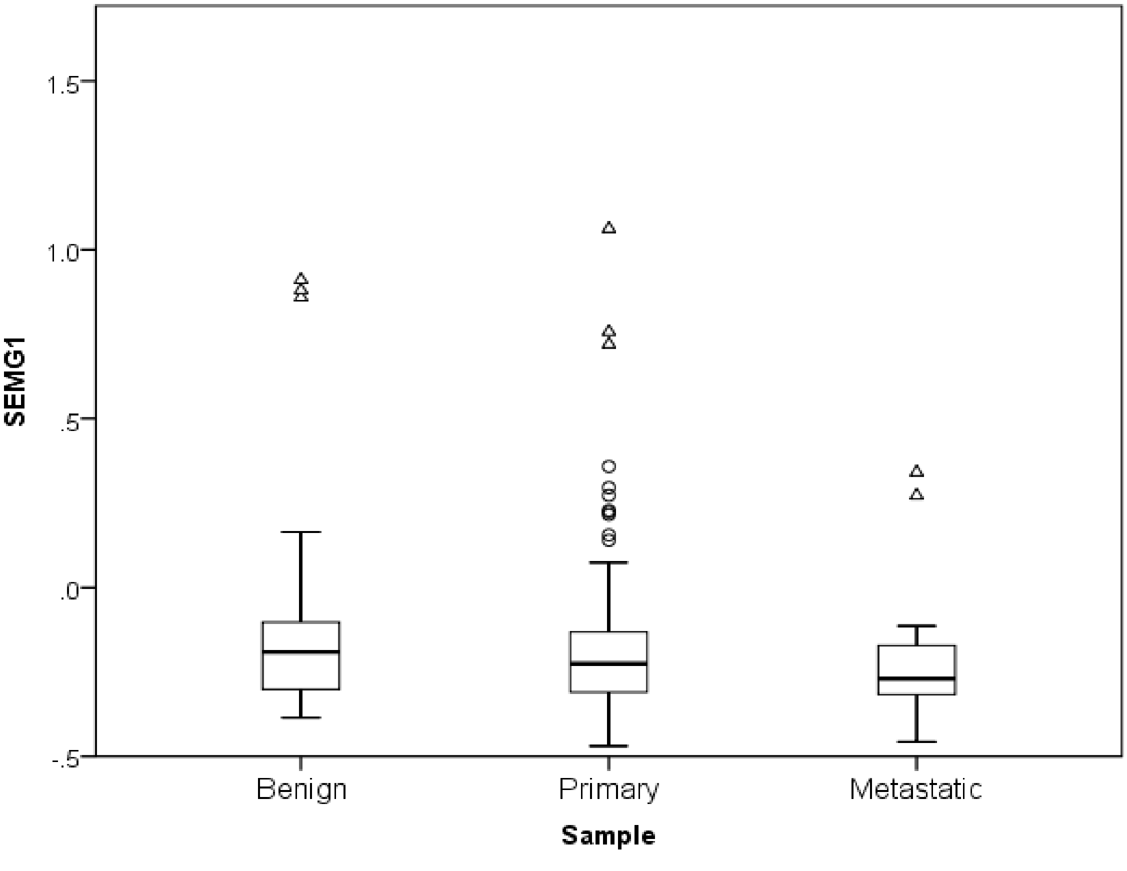
1.5
1.0
.5
.0
-.5
SEMG1
Benign
Primary
Metastatic
Sample

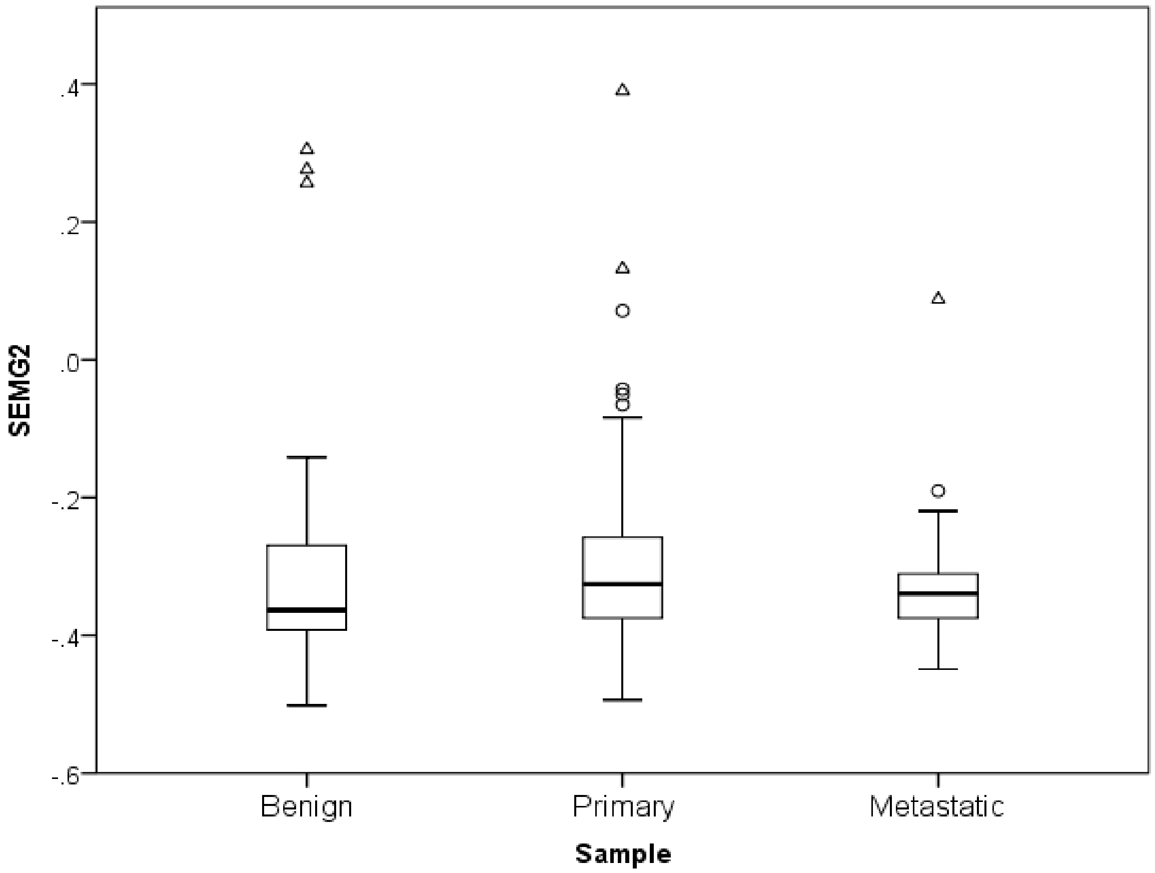

.4
.2
.0
-.2
-.4
-.6
SEMG2
Benign
Primary
Metastatic
Sample


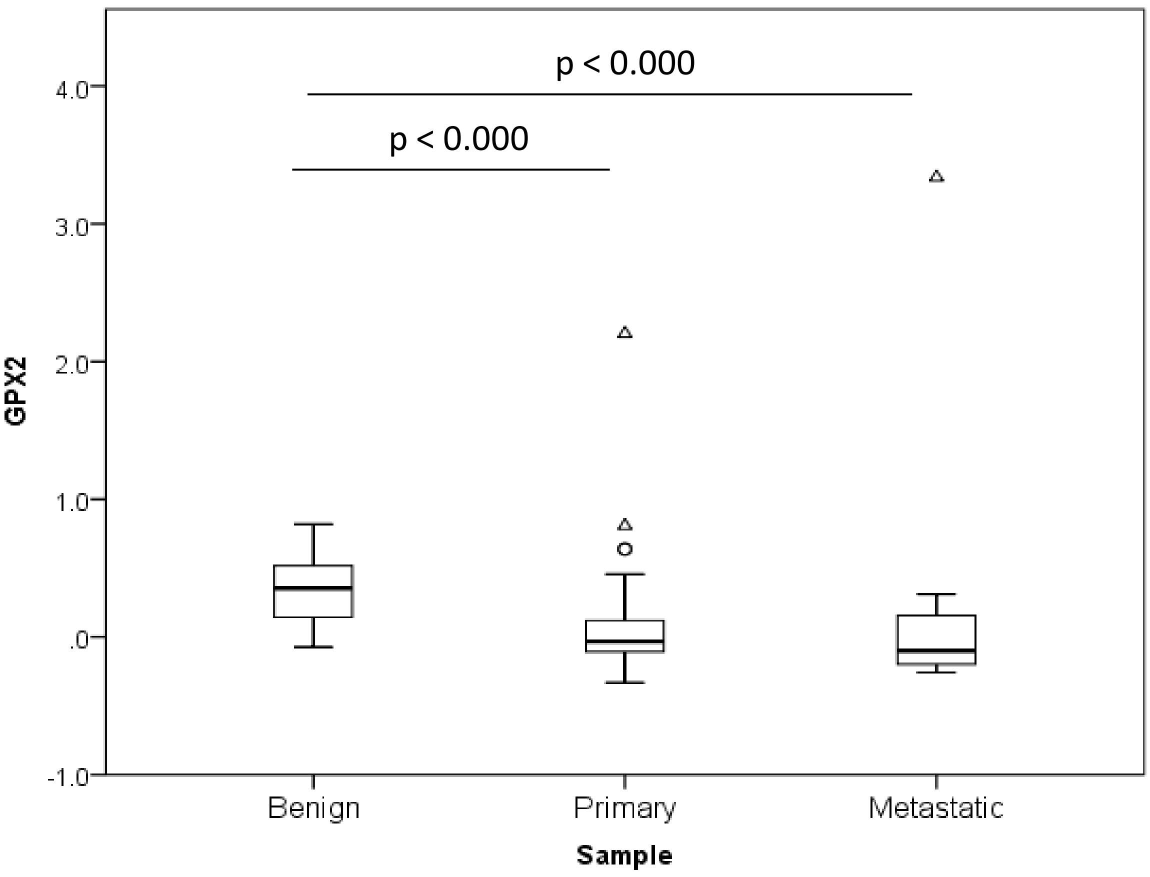

p < 0.000
p < 0.000
4.0
3.0
2.0
1.0
.0
-1.0
GPX2
Benign
Primary
Metastatic
Sample

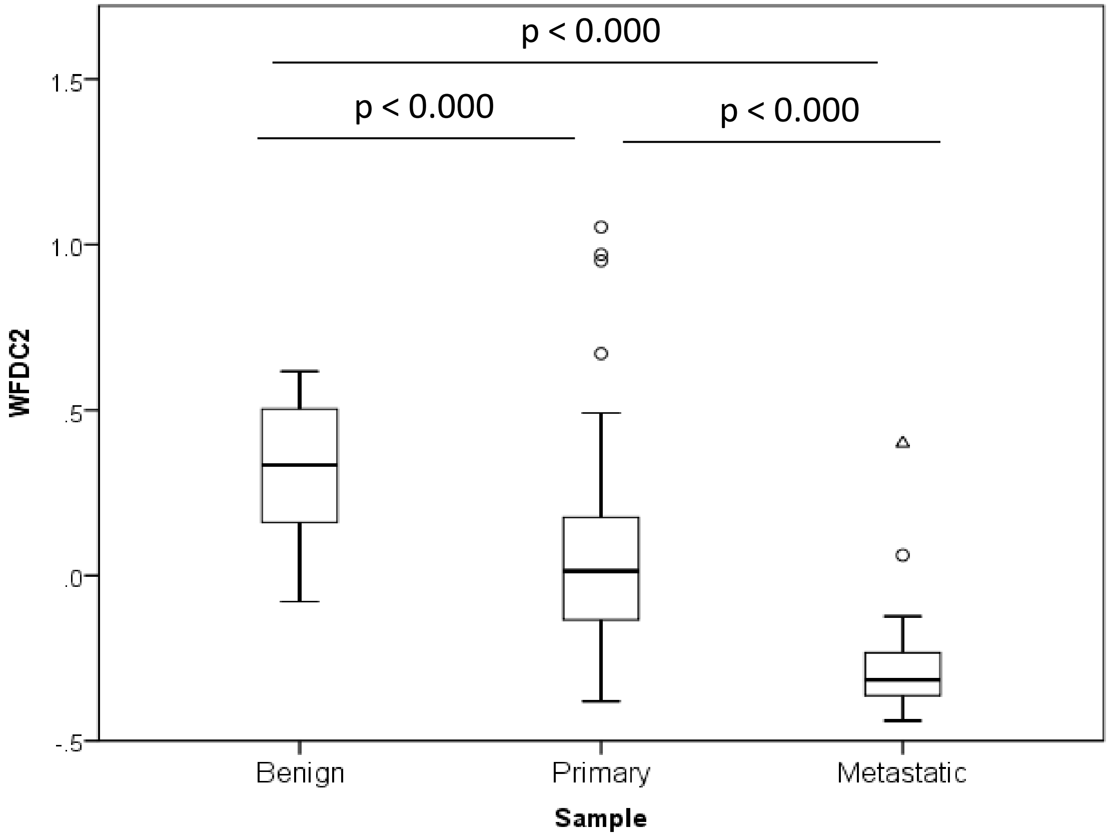

p < 0.000
p < 0.000
p < 0.000
1.5
1.0
.5
.0
-.5
WFDC2
Benign
Primary
Metastatic
Sample

### Taylor Dataset
### Summary of Results

| P values | Benign vs. Primary | Benign vs. Metastases | Primary vs. Metastases |
|---|---|---|---|
| ORE51E2 | **0.009** | **< 0.000** | |
| SIM2 | **< 0.000** | **< 0.000** | |
| ACTC1 | **< 0.000** | **< 0.000** | **< 0.000** |
| HPN | **< 0.000** | **< 0.000** | |
| SLC45A2 | **< 0.000** | **< 0.000** | |
| TDRD1 | **< 0.000** | **< 0.000** | |
| AMARC | **< 0.000** | **< 0.000** | |
| DLX1 | **< 0.000** | **< 0.000** | |
| GSTM1 | | **0.07** | **0.027** |
| SERPINA5 | | | |
| HOXC4 | **< 0.000** | **< 0.000** | **< 0.000** |
| COMP | | | |
| SLC39A2 | | | |
| SEMG1 | | | |
| SEMG2 | | | |
| GPX2 | **< 0.000** | **< 0.000** | |
| WFDC2 | **< 0.000** | **< 0.000** | **< 0.000** |

Green: Upregulation (e.g . High in primary vs. Benign)
Red: Downregulation (e.g . Low in primary vs. Benign)

### Summary for genes listed

| Gene name | Lapointe | Ross-Adams | Taylor | Up vs. down* |
|---|---|---|---|---|
| ORE51E2 | ✓ | ✓ | ✓ | Up |
| SIM2 | ✓ | ✓ | ✓ | Up |
| ACTC1 |  | ✓ | ✓ | Down |
| HPN | ✓ | ✓ | ✓ | Up |
| SLC45A2 | ✓ | ✓ | ✓ | Up |
| TDRD1 |  | ✓ | ✓ | Up |
| AMARC | ✓ | ✓ | ✓ | Up |
| DLX1 |  | ✓ | ✓ | Up |
| GSTM1 |  | ✓ | X | Down |
| SERPINA5 | ✓ | ✓ | X | Down |
| HOXC4 | ✓ | ✓ | ✓ | Up |
| HOXC6 |  | ✓ |  | Up |
| COMP | X | ✓ | X | Up |
| SLC39A2 | ✓ | ✓ | X | Down |
| SEMG1 | X | X | X | --- |
| SEMG2 | X | X | X | --- |
| GPX2 | ✓ | ✓ | ✓ | Down |
| PCA3 | ✓ | ✓ |  | Up |
| WFDC2 | ✓ | ✓ | ✓ | Down |

Green: significant difference in expression between benign and primary tumours at p lower than 0.05; Red: no significant difference; Black: not available.* Consistant up vs. down-regulation in primary tumours vs. benign
